# Hmx3a has essential functions in zebrafish spinal cord, ear and lateral line development

**DOI:** 10.1101/2020.01.23.917468

**Authors:** Samantha J. England, Gustavo A. Cerda, Angelica Kowalchuk, Taylor Sorice, Ginny Grieb, Katharine E. Lewis

## Abstract

Transcription factors that contain a homeodomain DNA-binding domain have crucial functions in most aspects of cellular function and embryonic development in both animals and plants. Hmx proteins are a sub-family of NK homeodomain-containing proteins that have fundamental roles in development of sensory structures such as the eye and the ear. However, Hmx functions in spinal cord development have not been analyzed. Here we show that zebrafish (*Danio rerio*) *hmx2* and *hmx3a* are co-expressed in spinal dI2 and V1 interneurons, whereas *hmx3b*, *hmx1* and *hmx4* are not expressed in spinal cord. Using mutational analyses, we demonstrate that, in addition to its previously reported role in ear development, *hmx3a* is required for correct specification of a subset of spinal interneuron neurotransmitter phenotypes, as well as correct lateral line progression and survival to adulthood. Surprisingly, despite similar expression patterns of *hmx2* and *hmx3a* during embryonic development, zebrafish *hmx2* mutants are viable and have no obviously abnormal phenotypes in sensory structures or neurons that require *hmx3a*. In addition, embryos homozygous for deletions of both *hmx2* and *hmx3a* have identical phenotypes to severe *hmx3a* single mutants. However, mutating *hmx2* in hypomorphic *hmx3a* mutants that usually develop normally, results in abnormal ear and lateral line phenotypes. This suggests that while *hmx2* cannot compensate for loss of *hmx3a*, it does function in these developmental processes, although to a much lesser extent than *hmx3a*. More surprisingly, our mutational analyses suggest that Hmx3a may not require its homeodomain DNA-binding domain for its roles in viability or embryonic development.

## Introduction

Homeobox-containing genes and the Homeodomain-containing transcription factors that they encode, have crucial functions in most aspects of cellular function and embryonic development in both animals and plants (Burglin and Affolter 2016). They were also some of the first examples discovered of invertebrate developmental genes that are highly conserved in vertebrates (Carrasco *et al*. 1984; Gehring 1985). One important subclass of homeodomain proteins are NK proteins. *NK* genes are evolutionarily ancient and are part of the ANTP megacluster, which also includes *Hox* and *ParaHox* genes. NK proteins have fundamental roles in the development of mesoderm, endoderm, the nervous system and the heart in all bilaterian animals examined so far (Wotton *et al*. 2010; Holland 2013; Treffkorn *et al*. 2018) and they are also found in sponges, one of the most basal animals still alive, and potentially the sister group to all other animals (Larroux *et al*. 2007; Fortunato *et al*. 2014; Pisani *et al*. 2015; Simion *et al*. 2017).

Hmx proteins (H6 Family Homeodomain proteins, previously called Nk5 or Nkx5 proteins, see Table S1) are a key sub-family of NK proteins. In vertebrates there are usually three or four different *Hmx* genes as *Hmx4* is only found in some species (Wotton *et al*. 2010). Interestingly, *Hmx2* and *Hmx3* are usually located adjacent to each other on the same chromosome and this is also the case for *Hmx1* and *Hmx4*, suggesting that both pairs of genes arose from tandem duplication events rather than the two rounds of whole genome duplication that occurred at the base of the vertebrates (Wotton *et al*. 2010). In teleosts, there are occasionally extra duplicates of one or more of these genes as the result of the additional genome duplication in this lineage, although interestingly, the retained genes are not consistent between different teleost species (Wotton *et al*. 2010). In zebrafish there are two *hmx3* genes, *hmx3a* and *hmx3b,* but only one *hmx1*, *hmx2* and *hmx4* gene.

Previous research has shown that Hmx2 and Hmx3 have crucial functions in ear development in mouse and our recent work shows that this is also the case for Hmx3a in zebrafish (Wang *et al*. 1998; Wang *et al*. 2001; Wang *et al*. 2004; Wang and Lufkin 2005; Hartwell *et al*. 2019). In mouse, both *Hmx2* and *Hmx3* mutants have ear defects and these are more severe in double mutants (Wang *et al*. 2001; Wang *et al*. 2004). *Hmx2* and *Hmx3* are also required for correct specification of the mouse hypothalamus (Wang *et al*. 2004) and morpholino knock-down experiments have suggested that they are required for correct lateral line development in zebrafish (Feng and Xu 2010). *Hmx2* and *Hmx3* are also expressed in two distinct domains in mouse spinal cord but the spinal cord functions of these genes are unknown (Bober *et al*. 1994; Wang *et al*. 2000; Wang *et al*. 2004).

Here we show that zebrafish *hmx2* and *hmx3a* are co-expressed in spinal dI2 and V1 interneurons, whereas *hmx3b*, *hmx1* and *hmx4* are not expressed in spinal cord. Using knock-down and mutational analyses, we demonstrate that, in addition to its role in ear development, *hmx3a* is required for correct specification of a subset of spinal cord interneuron neurotransmitter phenotypes as well as lateral line progression and viability (survival to adulthood). Our data suggest that in the absence of functional Hmx3a protein, a subset of dI2 spinal interneurons switch their neurotransmitter phenotype from glutamatergic (excitatory) to GABAergic (inhibitory). This is important because currently very little is known about how dI2 spinal interneuron neurotransmitter phenotypes are specified, or indeed how spinal cord excitatory neurotransmitter phenotypes in general are specified, and if neurons do not acquire the correct neurotransmitter phenotypes, they cannot function appropriately in spinal cord circuitry.

Surprisingly, despite the fact that *hmx2* and *hmx3a* have similar expression patterns during embryonic development and both genes are required for correct ear development in mouse, our mutational analyses did not uncover any requirement for *hmx2*, by itself, in viability, ear development, lateral line progression or specification of spinal cord interneuron neurotransmitter phenotypes in zebrafish. This is surprising, especially given that embryos injected with a *hmx2* morpholino have reduced numbers of spinal cord glutamatergic neurons and a corresponding increase in the number of inhibitory spinal cord neurons and that embryos injected with both *hmx2* and *hmx3a* morpholinos have more severe spinal cord phenotypes than single knock-down embryos. Zebrafish *hmx2* mutants are viable and have no obviously abnormal phenotypes in these sensory structures and neurons that require *hmx3a*, even when almost all of the *hmx2* locus is deleted. (In our most severe mutant allele, *hmx2^SU39^,* only 84 nucleotides of 5’ and 60 nucleotides of 3’ coding sequence remain). In addition, zebrafish embryos homozygous for deletions of both *hmx2* and *hmx3a* have identical phenotypes to severe *hmx3a* single mutants. However, mutating *hmx2* in hypomorphic *hmx3a^SU42^* mutants, that usually develop normally, results in abnormal ear and lateral line progression phenotypes, suggesting that while *hmx2* cannot compensate for mutations in *hmx3a*, it does function in these developmental processes, although to a much lesser extent than *hmx3a*. Our analyses of homozygous mutant phenotypes for several different *hmx3a* mutant alleles also suggest that Hmx3a may not require its homeodomain for its roles in viability or embryonic development. This is surprising, as homeodomain proteins usually function by binding DNA through their homeodomain and regulating gene expression. In contrast, our mutational analyses suggest that Hmx3a may only require its N terminal-domain for its vital functions in viability and sensory organ and spinal cord interneuron development.

## Materials and Methods

### Ethics statement

All zebrafish experiments in this research were carried out in accordance with the recommendations and approval of either the UK Home Office or the Syracuse University IACUC committee.

### Zebrafish husbandry and fish lines

Zebrafish (*Danio rerio*) were maintained on a 14-h light/10-h dark cycle at 28.5^◦^C. Embryos were obtained from natural paired and/or grouped spawnings of wild-type (WT; AB, TL or AB/TL hybrid) fish, heterozygous or homozygous *hmx2*, *hmx3a* or *hmx2/3a* mutants (created as part of this study and reported here, see Fig. 4), *Tg*(*evx1:EGFP*)*^SU1^* or *Tg*(*evx1:EGFP*)*^SU2^* transgenic fish (Juárez-Morales *et al*. 2016), *Tg(UAS:mRFP*) transgenic fish (Balciuniene *et al*. 2013), heterozygous *mindbomb1 (mib1^ta52b^)* mutants (Jiang *et al*. 1996) or heterozygous or homozygous *hmx3a^sa23054^* mutants (Kettleborough *et al*. 2013).

### CRISPR mutagenesis and screening

The *hmx3a^SU3^* allele was described previously (Hartwell *et al*. 2019). With the exception of *hmx3a^sa23054^* (generated in the Zebrafish Mutation Project and obtained from ZIRC), we created all of the other *hmx2, hmx3a* and *hmx2/3a* double deletion mutants described in this paper using CRISPR mutagenesis. For all alleles, other than the *hmx2^MENTHU^* allele, we designed and synthesized single guide RNA (sgRNA) and *Cas9* mRNA as in (Hartwell *et al*. 2019). For the *hmx2^MENTHU^* allele, we designed the crRNA using the Microhomology-mediated End joining kNockout Target Heuristic Utility (MENTHU) tool (version 2.1.2), in the Gene Sculpt Suite (Ata *et al*. 2018; Mann *et al*. 2019). The MENTHU allele crRNA design was verified with CHOPCHOP (version 3.0.0) (Montague *et al*. 2014; Labun *et al*. 2016; Labun *et al*. 2019) and the CRISPR-Cas9 gRNA design checker tool (Integrated DNA Technologies). The *hmx2^MENTHU^* crRNA was purchased together with a universal 67mer tracrRNA (1072533) and Alt-R S.p. Cas9 Nuclease V3 (1081058) from integrated DNA Technologies. See Table S2 for gRNA sequences and Figure 4 for their genomic locations. *hmx2^SU35^, hmx2^SU36^, hmx2^SU37^* and *hmx3a^SU42^* alleles were all generated with a single sgRNA (sgRNA E for *hmx2^SU35^, hmx2^SU36^* and *hmx2^SU37^*, and sgRNA B for *hmx3a^SU42^*, Table S2, Figure 4). *hmx2^MENTHU^* was generated with a single crRNA (sgRNA D, Table S2, Figure 4). *hmx2^SU38^, hmx2^SU39^, hmx2;hmx3a^SU44^* and *hmx2;hmx3a^SU45^* alleles are all deletions, generated by combinatorial use of two sgRNAs. The *hmx2^SU38^* sgRNAs (sgRNAs E and F, Table S2, Figure 4) flank the homeodomain. For *hmx2^SU39^* we used the same 3’ sgRNA and a more 5’ sgRNA (sgRNAs C and F, Table S2, Figure 4). To make *hmx2;hmx3a* double deletion alleles we designed sgRNAs that flanked the two genes, which are adjacent on chromosome 17 (sgRNAs A and F, Table S2, Figure 4). In all cases except the *hmx2^MENTHU^* mutant, we injected 2-4 nl of a mixture of 200 ng/µl of each sgRNA + 600 ng/µl *nls-ZCas9-nls* mRNA into the single-cell of a one-cell stage AB WT embryo. To create the *hmx2^MENTHU^* mutant allele, we injected 1 nl of a 5 µM crRNA:tracrRNA:Cas9 ribonucleoprotein complex into the single-cell of a very early one-cell stage embryo from an incross of heterozygous *hmx3a^SU42^* fish. The 5 µM crRNA:tracrRNA:Cas9 ribonucleoprotein complex was synthesized as described in (Hoshijima *et al*. 2019). *hmx3a^SU43^* is a *hsp70:Gal4* knock-in allele. We co-injected a donor template containing *Gal4*, under the control of a minimal *hsp70* promoter, (*pMbait-hsp70:Gal4*, a kind gift of Dr Shin-ichi Higashijima (Kimura *et al*. 2014)) with two sgRNA molecules: one specifically targeting *hmx3a* (sgRNA B, see Table S2 and Figure 4), and one, (*Mbait* sgRNA (GGCTGCTGCGGTTCCAGAGG)), specifically linearizing the donor template *in vivo*, into the single-cell of one-cell stage embryos from an incross of heterozygous *Tg(UAS:mRFP)* fish. For these experiments, embryos were injected with 2-4 nl of a mixture of 130 ng/µl of each sgRNA + 180 ng/µl *nlz-ZCas9-nls* mRNA + 66 ng/µl *pMbait-hsp70:Gal4* donor DNA. We screened injected embryos for RFP fluorescence in patterns consistent with *hmx3a* expression (i.e. ear, lateral line primordium and/or spinal cord) from 1 day post fertilization (d) onwards and raised injected embryos displaying appropriate expression patterns to adulthood. We then assessed germline transmission by outcrossing to heterozygous or homozygous *Tg(UAS:mRFP)* fish. Gal4 expression in *hmx3a^SU43^* recapitulates *hmx3a* spinal expression but is not expressed in the ear or lateral line primordium (data not shown).

We identified founder fish for ***hmx2^SU35^****, **hmx2******^SU36^****, **hmx2******^SU37^*** and ***hmx3a^SU42^*** alleles using high resolution melt analysis (HRMA), and the supermix and amplification programs described in (Hartwell *et al*. 2019). For the PCRs described below, we used Phusion High-Fidelity DNA Polymerase (M0530L, NEB) unless otherwise stated. HRMA primers and PCR primers for sequencing are provided in Table S2.

We used the following PCR conditions to identify ***hmx2^SU38^*** founder fish: 98.0°C for 30 seconds, 35 cycles of: 98.0°C for 10 seconds, 67.0°C for 30 seconds, 72.0°C for 40 seconds, followed by a final extension at 72.0°C for 5 minutes. We distinguished the mutant allele by gel electrophoresis on a 1% TAE agarose gel (110V for 30 minutes). The WT allele generated a 1098 bp product, compared with a 671 bp mutant allele product. The PCR primer sequences are provided in Table S2.

We used nested PCR to identify ***hmx2^SU39^*** founder fish, with the following conditions: <u>Nested PCR 1:</u> 98.0°C for 30 seconds, 35 cycles of: 98.0°C for 10 seconds, 69.0°C for 20 seconds, 72.0°C for 75 seconds, followed by a final extension at 72.0°C for 5 minutes. The mutant allele was distinguished by gel electrophoresis on a 1% TAE agarose gel (110V for 30 minutes). The WT allele generated a 2012 bp product (which may or may not be detected on the gel), compared with a 576 bp mutant allele product. We then diluted the nested 1 PCR product 1:5 in sterile distilled water and performed the <u>Nested PCR 2</u> reaction using the following conditions: 98.0°C for 30 seconds, 35 cycles of: 98.0°C for 10 seconds, 66.0°C for 20 seconds, 72.0°C for 60 seconds, followed by a final extension at 72.0°C for 5 minutes. The mutant allele was distinguished by gel electrophoresis on a 1% TAE agarose gel (110V for 30 minutes). The WT allele generated a 1758 bp product compared with a 322 bp mutant allele product. All PCR primer sequences are provided in Table S2.

We identified ***hmx2^MENTHU^*** F0 embryos by PCR, followed by sequencing with the FW primer that generated the amplicon (Table S2). The PCR was performed on DNA extracted from individual embryos using the following conditions: 98.0°C for 30 seconds, 35 cycles of: 98.0°C for 10 seconds, 64.0°C for 20 seconds, 72.0°C for 15 seconds, followed by a final extension at 72.0°C for 5 minutes. We assayed that the PCR was successful by gel electrophoresis on a 2.5% TBE agarose gel (100V for 40 minutes). The PCR generates a 155 bp product. The PCR product was purified using EZ-10 Spin Column PCR Products Purification Kit (BS664, Bio Basic) and eluted in 30 µl sterile water prior to sequencing.

We used either assessment of germline transmission, as described above, or PCR to identify ***hmx3a^SU43^*** founder fish. PCR conditions were: 98.0°C for 30 seconds, 35 cycles of: 98.0°C for 10 seconds, 69.0°C for 20 seconds, 72.0°C for 60 seconds, followed by a final extension at 72.0°C for 5 minutes. The mutant allele was distinguished by gel electrophoresis on a 1% TAE agarose gel (110V for 30 minutes). A 1471 bp PCR product was only generated by fish heterozygous for the allele. It was not produced from WT animals since the reverse primer only recognizes the inserted donor DNA sequence. The PCR primer sequences are provided in Table S2

We identified ***hmx2;hmx3a^SU44^*** and ***hmx2;hmx3a^SU45^*** founder fish by nested PCR, using the following conditions: <u>Nested PCR 1:</u> 98.0°C for 30 seconds, 35 cycles of: 98.0°C for 10 seconds, 67.0°C for 20 seconds, 72.0°C for 30 seconds, followed by a final extension at 72.0°C for 5 minutes. The mutant allele was distinguished by gel electrophoresis on a 1% TAE agarose gel (110V for 30 minutes). The WT product was too large to be generated by these PCR conditions, so only heterozygous animals are detected by the presence of a 514 bp product on the gel. We then diluted the nested 1 PCR 1:5 in sterile distilled water and performed the <u>Nested PCR 2</u> reaction using the following conditions: 98.0°C for 30 seconds, 35 cycles of: 98.0°C for 10 seconds, 66.0°C for 20 seconds, 72.0°C for 30 seconds, followed by a final extension at 72.0°C for 5 minutes. The mutant allele was distinguished by gel electrophoresis on a 1% TAE agarose gel (110V for 30 minutes). Again, the WT product was too large to be generated by these PCR conditions, so only heterozygous animals were detected by the presence of a 445 bp product.

Once stable lines were established, we identified ***hmx2^SU3^****^5^* fish by PCR, followed by sequencing (the mutation introduces a 1 bp insertion that cannot be resolved by restriction digestion, and we cannot distinguish heterozygotes from homozygotes using HRMA). We performed this PCR using Taq DNA Polymerase (M0320S, NEB) and the following conditions: 95.0°C for 30 seconds, 35 cycles of: 95.0°C for 20 seconds, 52.0°C for 30 seconds, 68.0°C for 45 seconds, followed by a final extension at 68.0°C for 5 minutes. The PCR primer sequences are provided in Table S2. We used HRMA and the conditions described above to identify ***hmx2^SU36^*** stable mutants. Homozygous mutants segregate from heterozygous animals by the scale of their deflection in the HRMA plot. We identified ***hmx2^SU37^*** mutants by performing the PCR used to sequence *hmx2^SU35^* stable mutants (see above and Table S2). When we analyzed the products on a 1% TAE gel (110V for 30 minutes), the WT allele generated a 580 bp product, compared with a 528 bp mutant product. We identified ***hmx2^SU38^*** stable mutants using the same PCR conditions initially used to identify founders (see above and Table S2). We used the same nested PCR conditions to identify ***hmx2^SU39^*** mutants. However, the WT product was not always visible on the gel. Therefore, we also performed a WT amplicon PCR identical to that described above for identifying stable *hmx2^SU35^* fish, as this genomic region is only present in WT and heterozygous animals (see also Table S2).

We identified stable ***hmx3a^SU42^*** mutants by PCR, using Taq DNA Polymerase (M0320S, NEB) and the following conditions: 94.0°C for 2 minutes, 35 cycles of: 94.0°C for 30 seconds, 64.9°C for 30 seconds, 72.0°C for 30 seconds, followed by a final extension at 72.0°C for 2 minutes. The PCR primer sequences are provided in Table S2. Whilst the mutant PCR product (321 bp) could sometimes be distinguished from the WT product (331 bp) by running on a 2% TBE gel (100V for 55 minutes), the mutation also deletes a BanI restriction site. Following digestion with BanI (R0118S, NEB), the products were run on a 2% TBE gel (100V for 40 minutes). The WT amplicon digested to completion, producing 120 bp + 211 bp bands, whereas the mutant product did not cut. We identified stable ***hmx3a^SU^****^3^* mutants by running the same PCR used to identify *hmx3a^SU42^* mutants (see above and Table S2). The insertion in *hmx3a^SU3^* was easily visualized on a 2% TBE gel. The WT product was 331 bp, compared to a mutant product of 400 bp. Since the PCR used to detect ***hmx3a^SU43^*** mutants was specific to the inserted donor DNA, and the WT amplicon in *hmx2;hmx3a^SU44^* and *hmx2;hmx3a^SU45^* mutants was too large to detect using the nested PCR conditions, for these alleles we also performed a WT amplicon PCR to distinguish WTs from heterozygotes. The WT amplicon PCR was identical to that performed prior to BanI digestion on *hmx3a^SU42^* mutants (see above and Table S2). For ***hmx3a^SU4^****^3^*, the WT amplicon PCR results were compared to the PCR results (identical PCR to that first used to identify founders, see above), and for ***hmx2;hmx3a^SU44^*** and ***hmx2;hmx3a^SU45^***, the WT amplicon PCR results were compared to the Nested 2 PCR results (identical Nested 2 PCR to that first used to identify founders – see above and Table S2).

In all cases, stable F1 heterozygous fish were confirmed by sequencing. See Fig. 4 for details of how individual mutant alleles differ from one another. To further confirm the mutant sequences of *hmx2^SU39^* and *hmx3a^SU42^*, we extracted total RNA from embryos produced by incrosses of homozygous viable adults using TRIzol Reagent (15596018, ThermoFisher Scientific) and the RNEasy Mini Kit (74104, Qiagen). Total RNA was converted to cDNA using the iScript cDNA synthesis kit (1708891, Bio-Rad). We performed transcript-specific PCRs using the following primers and conditions: *hmx2*-FW: TGAACTGTTATGAGACGAGAATGAA and *hmx2*-RV: GTGTATTTTGTACGTCTTAGTGTGTGT (PCR: 98.0°C for 30 seconds; 35 cycles of: 98.0°C for 10 seconds, 64.2°C for 20 seconds, 72.0°C for 30 seconds, followed by final extension at 72.0°C for 5 minutes) or *hmx3a*-FW: AACCGCGTTTAAGTTCCCATTG and *hmx3a*-RV: GTGCGAGTAGTAAACCGGATGAG (PCR: 98.0°C for 30 seconds; 35 cycles of: 98.0°C for 10 seconds, 71.0°C for 20 seconds, 72.0°C for 30 seconds, followed by final extension at 72.0°C for 5 minutes). We then confirmed these homozygous mutant transcript sequences by sequencing.

### Morpholino injections

For single knockdown (SKD) translation-blocking experiments (Fig. 3), 3.5 nl of a mixture containing either 2 ng/nl of a translation-blocking *hmx2* morpholino (5’ TTCCGCTGTCCTCCGAATTATTCAT) or 2 ng/nl of a translation-blocking *hmx3a* morpholino (5’ ACGTATCCTGTGTTGTTTCGGGCAT) plus 5 ng/nl of a control zebrafish *p53* morpholino (5’ GCGCCATTGCTTTGCAAGAATTG) was injected into the single-cell of a one-cell stage WT embryo. For double knockdown (DKD) experiments with translation-blocking morpholinos (Fig. 3), 3.5 nl of a mixture containing 2 ng/nl of both translation-blocking *hmx* morpholinos plus 5 ng/nl of the control zebrafish *p53* morpholino was injected. For DKD splice-blocking experiments (Fig. S1), 4 nl of a mixture containing 5 ng/nl of both a splice-blocking *hmx2* morpholino (5’ GGCACCTGCAACCAA TGCGACACAC) and a splice-blocking *hmx3a* morpholino (5’ TGCTGCTACAGTAATAGAGGCCAAA), plus 7 ng/nl of the control zebrafish *p53* morpholino was injected (all morpholinos obtained from Gene Tools). DKD but not SKD embryos exhibit delayed development from somitogenesis stages onwards when compared to uninjected controls. To circumvent this, they were incubated at 32°C from 9 hours post fertilization (h) onwards, whereas the uninjected controls remained at 28.5°C. This ensured that control and injected embryos reached the desired developmental stage of 27 h at approximately the same time. The lateral line primordium does not migrate in DKD animals (Fig. 3B), so this could not be used to stage injected embryos. Instead, these embryos were visually inspected and fixed when they displayed the same head-trunk angle, head size and eye size as prim-staged uninjected control embryos (Kimmel 1995). Migration of the lateral line primordium is unaffected in SKD embryos, so these were prim-staged prior to fixing for experiments (Kimmel 1995). Morpholino injections always produce a spectrum of phenotypes, since it is hard to ensure that every cell receives the same dose. Therefore, prior to fixing at 27 h, we removed any embryos with severely abnormal morphology (stunted length and/or severely developmentally delayed, likely caused by receiving too much morpholino). Embryos injected with *hmx2/3a* morpholinos (SKD and DKD) display a slight curled-tail-down morphology. Embryos that lacked this morphology (and may therefore not have received any or sufficient morpholino) were also removed before fixing.

For the mRNA + morpholino rescue experiments, we co-injected either each individual or both translation-blocking *hmx* morpholinos (at the same volume and dose described above), together with a total dose of up to 500 pg of morpholino-resistant (MOR) full-length *hmx2* or *hmx3a* mRNA. Both *hmx* mRNAs had 7 nucleotides altered in the morpholino recognition sequence. Each change was in the third nucleotide of a codon. This codon wobble was used so that the same amino acid was encoded in each case, but the mRNA would not be recognized by the morpholino. The protein encoded by the injected RNA is therefore the same as either endogenous Hmx2 or Hmx3a.

WT *hmx2*: ATG AAT AAT TCG GAG GAC AGC (Met, Asn, Asn, Ser, Glu, Asp, Ser)
*MOR-hmx2*: ATG AAC AAC TCC GAA GAT AGT (Met, Asn, Asn, Ser, Glu, Asp, Ser)
WT *hmx3a*: ATG CCC GAA ACA ACA CAG GAT ACG (Met, Pro, Glu, Thr, Thr, Gln, Asp, Thr)
*MOR-hmx3a*: ATG CCG GAG ACT ACT CAA GAC ACC (Met, Pro, Glu, Thr, Thr, Gln, Asp, Thr)

Reverse transcription PCR (RT-PCR) was performed to assess the efficiency of *hmx2;hmx3a* DKD by splice-blocking morpholinos (Fig. S1A, C). At 27 h, separate pools of 25 injected embryos (injected at the one-cell stage with the morpholino dose and volume described above) and 25 uninjected control embryos were homogenised in 200 µl of Tri Reagent Solution (AM9738, ThermoFisher Scientific). Total RNA was extracted and purified as per the manufacturer’s instructions, prior to resuspending in 20 µl of sterile distilled water. To remove genomic DNA, 2.4 µl of RQ1 DNase Buffer and 2 µl of RQ1 RNase-Free DNase (M6101, Promega) was added to each RNA sample and incubated for 15 minutes at 37°C. Heat-inactivation of the DNase was performed for 10 minutes at 65°C. 20 µl RT-PCRs were performed as per the manufacturer’s instructions using the Qiagen One-Step RT-PCR kit (210210, Qiagen) and the following primers: *hmx2* RT-PCR E1-2 FW: TCAAGTTTCACGATCCAGTCTA and *hmx2* RT-PCR E1-2 RV: ATAAACCTGACTCCGAGAGAAA, *hmx3a* RT-PCR E1-2 FW: GTCAAAGCCTAAGCCTATTTTG and *hmx3a* RT-PCR E1-2 RV: TCACTCTTCTTCCAGTCGTCTA, and *actb1* RT-PCR E3-4 FW: GAGGTATCCTGACCCTCAAATA and *actb1* RT-PCR E3-4 RV: TCATCAGGTAGTCTGTCAGGTC and universal PCR program: 50°C for 30 min; 95°C for 15min; 35 cycles of: 95°C for 30 seconds, 57°C for 45 seconds and 72°C for 1 minute; followed by a final extension for 10 minutes at 72°C. Parallel reactions, omitting reverse transcriptase and performed on non DNase-treated samples were used to verify the non-spliced (genomic) PCR product. 10 µl of each RT-PCR product was assessed by electrophoresing for 40 minutes at 100 V on a 2% TBE agarose gel. The *hmx2* RT-PCR E1-2 primers generate either a 1204 bp genomic (unspliced) or 426 bp spliced product. The *hmx3a* RT-PCR E1-2 primers generate either a 779 bp genomic (unspliced) or 393 bp spliced product. The *actb1* RT-PCR E3-4 primers generate either a 697 bp genomic (unspliced) or 387 bp spliced product (Fig. S1A-D).

To assess whether genetic compensation occurs in either *hmx2^SU39^* or *hmx3a^SU42^* mutants, which lack obvious phenotypes, or *hmx3a^SU3^* mutants, which have milder spinal cord phenotypes than *hmx2/3a* DKD embryos, we injected the same dose of either *hmx2* + *p53* MOs (*hmx2^SU39^*) or *hmx3a* + *p53* MOs (*hmx3a^SU42^, hmx3a^SU3^*) as described above, into the single-cell of one-cell stage embryos generated from incrosses of heterozygous *hmx2^SU39^, hmx3a^SU42^* or *hmx3a^SU3^* parents respectively. If genetic compensation is occurring, the upregulated compensating gene(s) will not be knocked-down by the *hmx* morpholino and the phenotype of homozygous mutants should be unchanged. In contrast, WT and heterozygous animals, which contain at least one WT copy of the respective *hmx* gene will be susceptible to the *hmx* morpholino and should exhibit stronger, morphant-like phenotypes. For these experiments, while we removed any embryos with severely abnormal morphology, we did not remove embryos that lacked the curled-tail-down morphology, incase these were morpholino-resistant mutant embryos. After fixing, we performed an *in situ* hybridization for the glutamatergic marker, *slc17a6a/b*. We visually inspected the embryos on a dissecting microscope and categorized them as either the stronger, morphant-like phenotype (large reduction in the number of *slc17a6a/b*-expressing cells) or a more subtle phenotype (WT-like in the case of *hmx2^SU39^* and *hmx3a^SU42^*, or a smaller reduction in the number of *slc17a6a/b*- expressing cells in the case of *hmx3a^SU3^* embryos). Embryos within each class were then genotyped as described in the CRISPR mutagenesis section above.

### Genotyping

We isolated DNA for genotyping from both anesthetized adult fish and fixed embryos via fin biopsy or head dissections, respectively. For assaying ear phenotypes, we dissected tail tips instead. We genotyped the *hmx* CRISPR mutants as described above. For *mib1^ta52b^* and *hmx3a^sa23054^* mutants, we used KASP assays designed by LGC Genomics LLC. KASP assays use allele-specific PCR primers, which differentially bind fluorescent dyes that we quantified with a Bio-Rad CFX96 real-time PCR machine to distinguish genotypes. The proprietary primers used were: *mib_ta52b* and *hmx3a_sa23054*. Heads or tail tips of fixed embryos were dissected in 70% glycerol/30% distilled water with insect pins. Embryo trunks were stored in 70% glycerol/30% distilled water at 4^◦^C for later analysis. For all experiments except Phalloidin staining experiments, DNA was extracted via the HotSHOT method (Truett *et al*. 2000) using 10 μl of 50 mM NaOH and 1 μl of 1M Tris-HCl (pH-7.4). For Phalloidin staining experiments, the tail up until the end of the yolk extension was dissected in 70% glycerol/30% distilled water as described above and transferred to PBST. The PBST was then replaced with 50 µl of DNA extraction buffer (10 mM Tris, pH 8.0, 10 mM EDTA, 200 mM NaCl, 0.5% SDS, 200 µg/ml Proteinase K (Proteinase K, recombinant, PCR Grade, 3115879001, Sigma Aldrich)), before incubating for 3 hours in a 55°C water bath. The samples were vortexed periodically to ensure thorough digestion of the tissue. Subsequently, the Proteinase K was inactivated by heating the samples for 10 minutes at 100°C, prior to centrifuging for 20 minutes at 14,000 rpm at room temperature to pellet debris. The supernatant was transferred to sterile microcentrifuge tubes before adding 20 µg UltraPure Glycogen (10814010, ThermoFisher Scientific) and 2 volumes of ice-cold RNase-free ethanol. Samples were precipitated at -20°C overnight. Genomic DNA was recovered by centrifugation at +4°C, followed by washing with 70% RNase-free ethanol and further centrifugation at +4°C. After carefully removing the ethanolic supernatant, the pellets were air dried for 5-10 minutes at room temperature before resuspending in 15 µl of sterile distilled water.

### *in situ* hybridization and immunohistochemistry

We fixed embryos in 4% paraformaldehyde / phosphate-buffered saline (PBS) and performed single and double *in situ* hybridizations and immunohistochemistry plus *in situ* hybridization double labeling experiments as previously described (Concordet *et al*. 1996; Batista *et al*. 2008). Sources of *in situ* hybridization probes are provided in Table S1. To amplify *in situ* hybridization probe templates for *hmx1* and *hmx3b*, we created cDNA from 27 h WT zebrafish embryos. We extracted total RNA by homogenizing 50–100 mg of embryos in 1 mL of TRIzol reagent (Ambion, 15596-026). We confirmed RNA integrity (2:1 ratio of 28S:18S rRNA bands) and quality (A260/A280 ratio of ∼2.0) using agarose gel electrophoresis and spectrophotometry respectively. We synthesized cDNA using Bio-Rad iScript Reverse Transcription Supermix kit (Bio-Rad, 170-8891). We amplified *hmx1* sequence from the cDNA using Phusion High-Fidelity DNA Polymerase (M0530L, NEB), primers *hmx1*-FW: CTGGTATATTTGCTCAAGACATGC and *hmx1-*RV: GCTTCTGCTGAACACAGTTCG and PCR conditions: 98.0°C for 10 seconds, 30 cycles of: 98.0°C for 60 seconds, 57.0°C for 30 seconds and 72.0°C for 30 seconds, followed by a final extension for 45 seconds at 72.0°C. The PCR product was assessed on a 1% TAE gel, before purifying with QIAquick PCR Purification Kit (28104, Qiagen). We used Taq DNA Polymerase (M0320S, NEB) to add 3’A overhangs prior to TOPO TA-cloning (K4600-01, Invitrogen). We then performed colony PCR using the same PCR primers and conditions used to amplify the *hmx1* sequence from cDNA. We extracted plasmid DNA from positive colonies using QIAprep Spin Miniprep Kit (27104, Qiagen) and then verified the sequence using standard SP6 and T7 primers for Sanger sequencing. To make the anti-sense RNA riboprobe, we linearized DNA with HindIII-HF (R3104S, NEB) and transcribed with T7 RNA Polymerase (10881767001, Roche). We used a PCR-based DNA template to make the *hmx3b* ISH probe. The reverse primer contains the T3 RNA Polymerase minimal promoter sequence (underlined). We used primers *hmx3b*-FW: GTGTGCCCGTCATCTACCAC and *hmx3b*-RV: <u>AATTAACCCTCACTAAAGGGA</u>TGAAGATGATGAAGATGCGCAAC, 27 h WT cDNA, Phusion High-Fidelity DNA Polymerase (M0530L, NEB) and PCR conditions: 94.0°C for 3 minutes, 35 cycles of: 94.0°C for 30 seconds, 56.5°C for 30 seconds and 72.0°C for 1.5 minutes, followed by a final extension step of 72.0°C for 10 minutes. We purified the template through phenol:chloroform:isoamyl alcohol extraction and precipitation with 0.2 M NaCl and ice-cold ethanol prior to *in situ* probe synthesis using 1 µg purified PCR product, T3 RNA Polymerase (11031171001, Roche) and DIG RNA Labeling Mix (11277073910, Roche).

Embryos older than 24 h were usually incubated in 0.003% 1-phenyl-2-thiourea (PTU) to prevent pigment formation. For some experiments we added 5% of dextran sulfate to the hybridization buffer (* in Table 1B). Dextran sulfate can increase specific staining in *in situ* hybridization experiments as it facilitates molecular crowding (Ku *et al*. 2004; Lauter *et al*. 2011).

In cases where we did not detect expression of a particular gene in the spinal cord, we checked for low levels of expression by exposing embryos to prolonged staining. In some cases, this produced higher background (diffuse, non-specific staining), especially in the hindbrain, where ventricles can sometimes trap anti-sense riboprobes.

To determine neurotransmitter phenotypes we used probes for genes that encode proteins that transport or synthesize specific neurotransmitters as these are some of the most specific molecular markers of these cell fates (Higashijima *et al*. 2004b; Higashijima *et al*. 2004c and references therein). A mixture of probes to *slc17a6a* and *slc17a6b* (previously called *vglut*), which encode glutamate transporters, was used to label glutamatergic neurons (Higashijima *et al*. 2004b; Higashijima *et al*. 2004c). GABAergic neurons were labeled using probes to *gad1b* (probes previously called *gad67a* and *gad67b*) (Higashijima *et al*. 2004b; Higashijima *et al*. 2004c). The *gad1b* gene encodes for a glutamic acid decarboxylase, which is necessary for the synthesis of GABA from glutamate. A mixture of probes (*glyt2a* and *glyt2b*) for *slc6a5* (previously called *glyt2*) was used to label glycinergic cells (Higashijima *et al*. 2004b; Higashijima *et al*. 2004c). *slc6a5* encodes for a glycine transporter necessary for glycine reuptake and transport across the plasma membrane.

The antibodies that we used for fluorescent *in situ* hybridization were mouse anti-Dig (200-002-156, Jackson ImmunoResearch, 1:5000) and rabbit anti-Flu (A889, Invitrogen, 1:2500). These were detected using secondary antibodies: goat anti-rabbit-HRP (G-21234, ThermoFisher Scientific, 1:750) and goat anti-mouse-HRP (G-21040, ThermoFisher Scientific, 1:750) and Tyramide SuperBoost Kits B40922 and B40915 (ThermoFisher Scientific).

For double fluorescent *in situ* hybridization and immunohistochemistry, after detection of the *in situ* hybridization reaction using Tyramide SuperBoost Kit B40915 (with HRP, Goat anti-mouse IgG and Alexa Fluor 594 Tyramide), embryos were washed 8 × 15min in PBST (PBS with 0.1% Tween-20) and incubated in Image-iT FX Signal Enhancer (ThermoFisher Scientific, I36933) for 30 mins at room temperature. Immunohistochemistry was performed using chicken polyclonal anti-GFP primary antibody (Ab13970, Abcam, 1:500) and a Goat anti-chicken IgY (H+L), Alexa Fluor 488 secondary antibody (A-11039, ThermoFisher Scientific, 1:1000).

### Phalloidin staining

4 d old embryos generated from incrosses of heterozygous *hmx2^SU39^* or *hmx2;hmx3a^SU44^* parents were fixed and processed for phalloidin staining as described in (Hartwell *et al*. 2019). Stained embryos were stored in DABCO (2% w/v 1,4-Diazabicyclo[2.2.2]octane (D27802, Sigma Aldrich) in 80% glycerol in sterile distilled water).

### qPCR analyses

We collected embryos from incrosses of AB wild-type parents and flash-froze them at 16-cell, 6 h, 14 h, 27 h and 48 h stages. We collected 40-50 embryos per biological replicate per developmental stage and performed duplicate biological replicates. We isolated total RNA by homogenizing each sample in 1 mL of TRIzol reagent (Ambion, 15596-026). Following chloroform extraction, we added 20 µg UltraPure Glycogen (10814010, ThermoFisher Scientific) to the aqueous phase followed by one volume of RNase-free ethanol. We performed RNA purification and genomic DNA removal using the Monarch Total RNA Miniprep Kit (T2010S, NEB) following manufacturer’s instructions for purifying TRIzol-extracted samples. RNA concentration was measured using Nanodrop 2000 (ND2000, ThermoFisher Scientific), prior to synthesizing cDNA using the Bio-Rad iScript Reverse Transcription Supermix kit (Bio-Rad, 170-8891). We also included controls lacking Reverse-Transcriptase to assay for the presence of genomic DNA contamination. qPCR was performed in triplicate for each sample using iTaq Universal SYBR Green Supermix (1725121, Bio-Rad) and a Bio-Rad CFX96 real-time PCR machine. The following qPCR primers were used:

*hmx2*-qPCR-FW: CCCATTTCAAGTTTCACGATCCAGTC,
*hmx2*-qPCR-RV: TGCTCCTCTTTGTAATCCGGTAG,
*hmx3a*-qPCR-FW: TTGATGGCAGCTTCTCCCTTTC,
*hmx3a*-qPCR-RV: ACTCTTCTTCCAGTCGTCTATGC,
*mob4*-qPCR-FW: CACCCGTTTCGTGATGAAGTACAA,
*mob4*-qPCR-RV: GTTAAGCAGGATTTACAATGGAG.

The *hmx2* and *hmx3a* primers were generated in this study. The *mob4* primers were generated by Hu and colleagues (Hu *et al*. 2016). They demonstrated that *mob4* is a more effective reference gene than *actb2* across a broad range of zebrafish developmental stages, including early stages where only maternal RNAs should be present (Hu *et al*. 2016). To generate amplification data the program used was: 95.0°C for 30 seconds, 40 cycles of: 95.0°C for 5 seconds, 63.3°C (*hmx2*)/64.5°C (*hmx3a*)/60.0°C (*mob4*) for 30 seconds, with imaging after each cycle. To assay amplification specificity and exclude false positives from primer dimers we then generated melt data using: 65.0°C for 30 seconds, 40 cycles of: 65.0°C-95.0°C, + 0.5°C/second increment, with each increment held for 5 seconds prior to imaging, 95.0°C for 15 seconds.

### Screening lateral line and otolith phenotypes

We examined whether any of the *hmx* mutants generated in this study had lateral line and/or fused otolith phenotypes, as reported for *hmx2;hmx3a* double-knockdown embryos (Feng and Xu 2010). To assay live lateral line phenotypes, we anaesthetized embryos from incrosses of heterozygous mutant fish in 0.016% tricaine (A5040, Sigma Aldrich) in embryo medium (EM, 5 mM NaCl, 0.17 mM KCl, 0.33 mM CaCl_2_·2H_2_O, 0.33 mM MgSO_4_·7H_2_O, 0.017% w/v (0.7mM) HEPES pH 7.8 and 0.00004% methylene blue in autoclaved reverse osmosis water) and mounted them on coverslip bridges (2 x 22 mm square glass coverslips (16004-094, VWR) glued together on either side of a 24 x 60 mm glass cover slip (12460S, ThermoFisher Scientific), overlaid with a third 22 mm square glass coverslip). Using a Zeiss Axio Imager M1 compound microscope, we located the tip of the lateral line primordium and counted the somite number adjacent to this position. We also used this method routinely to determine the developmental stage of embryos prior to fixing for *in situ* hybridization. To assay lateral line phenotypes in fixed embryos, we performed *in situ* hybridizations for *hmx3a* or *krt15* (both of which label the migrating primordium and neuromasts) and then determined the lateral line position as in live embryos. To examine live otolith phenotypes, embryos were raised until 3 d, before anaesthetizing (as for assessing live lateral line phenotypes) and examining the spatial location of otoliths in both ears. WT embryos have two otoliths in each ear: one smaller, anterior (utricular) otolith, and one larger, posterior (saccular) otolith. These are separate from each other and spatially distinct. We classified otoliths as fused if only one large, amalgamated otolith was visible in a mid-ventral position within the otic vesicle (see Fig. 5U).

### Imaging

Embryos were mounted in 70% glycerol:30% distilled water and Differential Interference Contrast (DIC) pictures were taken using an AxioCam MRc5 camera mounted on a Zeiss Axio Imager M1 compound microscope. Fluorescent images were taken on a Zeiss LSM 710 confocal microscope. Images were processed using Adobe Photoshop software (Adobe, Inc) and Image J software (Abramoff et al., 2004). In some cases, different focal planes were merged to show labeled cells at different medio-lateral positions in the spinal cord. All images were processed for brightness-contrast and color balance using Adobe Photoshop software (Adobe, Inc.). Images of control and mutant embryos from the same experiment were processed identically. Figures were assembled using Adobe Photoshop and Adobe Illustrator (Adobe, Inc.).

### Cell counts and statistics

In all cases except where noted to the contrary, cell counts are for both sides of a five-somite length of spinal cord adjacent to somites 6-10. Embryos were mounted laterally with the somite boundaries on each side of the embryo exactly aligned and the apex of the somite over the middle of the notochord. This ensures that the spinal cord is straight along its dorsal-ventral axis and that cells in the same dorsal/ventral position on opposite sides of the spinal cord will be directly above and below each other. Embryos from mutant crosses were counted blind to genotype. Labeled cells in embryos analyzed by DIC were counted while examining embryos on a Zeiss Axio Imager M1 compound microscope. We identified somites 6–10 in each embryo and counted the number of labeled cells in that stretch of the spinal cord. We adjusted the focal plane as we examined the embryo to count cells at all medio-lateral positions (both sides of the spinal cord; also see Batista *et al*. 2008; Batista and Lewis 2008; England *et al*. 2011; Hilinski *et al*. 2016; Juárez-Morales *et al*. 2016).

In some cases, cell count data were pooled from different experiments. Prior to pooling, all pairwise combinations of data sets were tested to determine if there were any statistically significant differences between them, as described below. Data were only pooled if none of the pairwise comparisons were statistically significantly different from each other. In addition, as *in situ* hybridization staining can vary slightly between experiments, we only compared different mutant results when the counts from their corresponding WT sibling embryos were not statistically significantly different from each other.

To determine whether differences in values are statistically significant, data were first analyzed for normality using the Shapiro-Wilk test. Data sets with non-normal distributions were subsequently analyzed using the Wilcoxon-Mann-Whitney test (also called the Mann Whitney U test). For data sets with normal distributions, the F-test for equal variances was performed, prior to conducting either a type 2 (for equal variances) or type 3 (for non-equal variances) student’s *t*-test. P values generated by Wilcoxon-Mann-Whitney, type 2 student’s *t*-test and type 3 student’s *t*-test are indicated by ^^^, ^+^ and ^§^ respectively. To control for type I errors, when comparing three or more experimental conditions, a one-way analysis of variance (ANOVA) test was performed. Prior to conducting ANOVA tests, data were first analysed for normality using the Shapiro-Wilk test, as described above. All data sets for ANOVA analysis had normal distributions and so were subsequently assessed for homogeneity of variances using Bartlett’s test. All of the data sets also had homogeneous (homoscedastic, Bartlett’s test p value >0.05) variances and so standard ANOVA analysis was performed. ANOVA results are reported as F(dfn,dfd) = f-ratio, p value = *x*, where F is the F-statistic, dfn = degree of freedom for the numerator of the F-ratio, dfd = degree of freedom for the denominator of theR-ratio, and *x* = the p value. For statistically significant ANOVA, to determine which specific experimental groups or groups differed, post hoc testing was performed. Since all ANOVA data sets had homogeneous (homoscedastic) variances, Tukey’s honestly significant difference post hoc test for multiple comparisons was performed. P values generated by Tukey’s honestly significant difference test are indicated by ^‡.^ Data are depicted as individual value plots and the *n*-values for each experimental group are also shown. For each plot, the wider red horizontal bar depicts the mean and the red vertical bars depict the standard error of the mean (standard error of the mean (S.E.M.) values are listed in Tables 1A & B). Individual data value plots were generated using Prism version 8.4.3 (GraphPad Software, San Diego, California USA, www.graphpad.com). To assess whether mutant phenotypes occurred at Mendelian frequencies, we performed Chi-squared tests. To test whether a small number of embryos with abnormal phenotypes was statistically significantly different from zero we performed a binomial distribution test, using the cumulative distribution function, the number of embryos without mutant phenotypes, the total number of embryos examined (n) and a probability argument of n-1/n. P values >0.05 support the null hypothesis that the number of embryos with abnormal phenotypes is not statistically significantly different from zero. Shapiro-Wilk and Wilcoxon-Mann-Whitney testing was performed in R version 3.5.1 (R Development Core Team 2005). The F-test, student’s t-test, Chi-squared test and binomial distribution test were performed in Microsoft Excel version 16.41. Bartlett’s testing, standard ANOVA, and Tukey’s honestly significant difference testing were performed in Prism version 8.4.3 (GraphPad Software, San Diego, California USA, www.graphpad.com).

**Table 1A.**
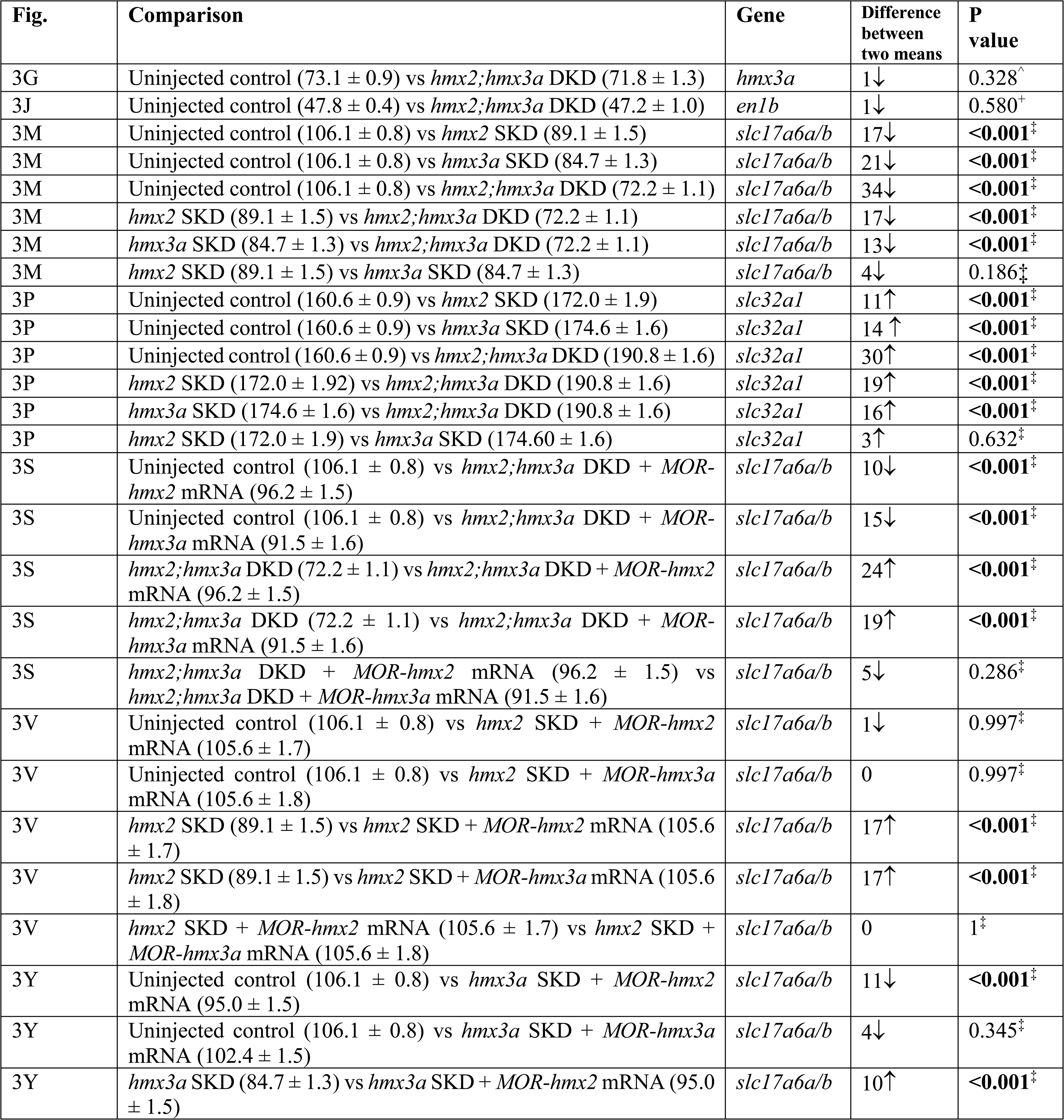
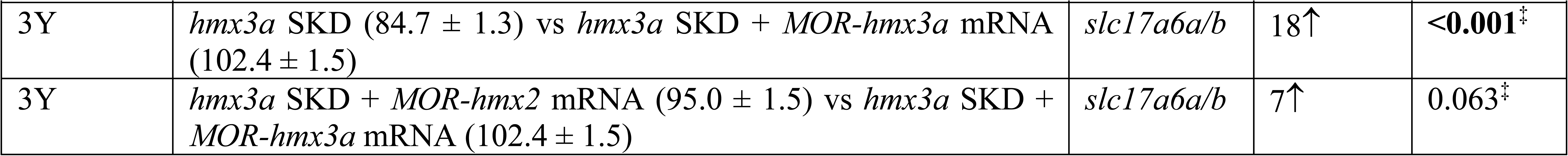
**<u>Statistical comparisons of numbers of cells expressing particular genes in morpholino knock-down experiments.</u>** Statistical comparisons between uninjected WT control and knockdown embryos. First column indicates the figure panel that contains the relevant individual value plots for the comparison. Second column states the genotypes being compared. Numbers within parentheses indicate mean numbers of cells ± S.E.M. In all cases, numbers are an average of at least 5 embryos and cells were counted in all dorsal-ventral spinal cord rows. All of the experiments were conducted on 27 h embryos. 27 h embryos fixed on different days varied slightly in stage from prim stage 9 to 12. This explains the small differences in numbers of cells labelled with a particular probe in uninjected WT control embryos in different experiments. Data from different days were only combined if there was no statistically significant difference between uninjected WT control embryos for each day (see materials and methods). Column three lists the gene that the cell counts and statistical comparison refer to. The fourth column indicates the difference between the two mean values for the embryos being compared. All values are rounded to the nearest whole number. ↑ = increase, ↓ = decrease. Last column shows the P value for the comparison, rounded to three decimal places. Statistically significant (P<0.05) values are indicated in bold. Statistical test used is indicated by superscript symbol: Wilcoxon-Mann-Whitney test (^^^), type 2 Student’s t-test (^+^), type 3 Student’s t-test (^§^) and Tukey’s honestly significant post-hoc test after ANOVA (^‡^). For a discussion of why particular tests were used, see materials and methods.

| Fig. | Comparison | Gene | Difference between two means | P value |
| --- | --- | --- | --- | --- |
| 3G | Uninjected control ($73.1 \pm 0.9$) vs *hmx2;hmx3a* DKD ($71.8 \pm 1.3$) | *hmx3a* | 1↓ | 0.328<sup>^</sup> |
| 3J | Uninjected control ($47.8 \pm 0.4$) vs *hmx2;hmx3a* DKD ($47.2 \pm 1.0$) | *en1b* | 1↓ | 0.580<sup>+</sup> |
| 3M | Uninjected control ($106.1 \pm 0.8$) vs *hmx2* SKD ($89.1 \pm 1.5$) | *slc17a6a/b* | 17↓ | <0.001<sup>‡</sup> |
| 3M | Uninjected control ($106.1 \pm 0.8$) vs *hmx3a* SKD ($84.7 \pm 1.3$) | *slc17a6a/b* | 21↓ | <0.001<sup>‡</sup> |
| 3M | Uninjected control ($106.1 \pm 0.8$) vs *hmx2;hmx3a* DKD ($72.2 \pm 1.1$) | *slc17a6a/b* | 34↓ | <0.001<sup>‡</sup> |
| 3M | *hmx2* SKD ($89.1 \pm 1.5$) vs *hmx2;hmx3a* DKD ($72.2 \pm 1.1$) | *slc17a6a/b* | 17↓ | <0.001<sup>‡</sup> |
| 3M | *hmx3a* SKD ($84.7 \pm 1.3$) vs *hmx2;hmx3a* DKD ($72.2 \pm 1.1$) | *slc17a6a/b* | 13↓ | <0.001<sup>‡</sup> |
| 3M | *hmx2* SKD ($89.1 \pm 1.5$) vs *hmx3a* SKD ($84.7 \pm 1.3$) | *slc17a6a/b* | 4↓ | 0.186<sup>‡</sup> |
| 3P | Uninjected control ($160.6 \pm 0.9$) vs *hmx2* SKD ($172.0 \pm 1.9$) | *slc32a1* | 11↑ | <0.001<sup>‡</sup> |
| 3P | Uninjected control ($160.6 \pm 0.9$) vs *hmx3a* SKD ($174.6 \pm 1.6$) | *slc32a1* | 14↑ | <0.001<sup>‡</sup> |
| 3P | Uninjected control ($160.6 \pm 0.9$) vs *hmx2;hmx3a* DKD ($190.8 \pm 1.6$) | *slc32a1* | 30↑ | <0.001<sup>‡</sup> |
| 3P | *hmx2* SKD ($172.0 \pm 1.92$) vs *hmx2;hmx3a* DKD ($190.8 \pm 1.6$) | *slc32a1* | 19↑ | <0.001<sup>‡</sup> |
| 3P | *hmx3a* SKD ($174.6 \pm 1.6$) vs *hmx2;hmx3a* DKD ($190.8 \pm 1.6$) | *slc32a1* | 16↑ | <0.001<sup>‡</sup> |
| 3P | *hmx2* SKD ($172.0 \pm 1.9$) vs *hmx3a* SKD ($174.60 \pm 1.6$) | *slc32a1* | 3↑ | 0.632<sup>‡</sup> |
| 3S | Uninjected control ($106.1 \pm 0.8$) vs *hmx2;hmx3a* DKD + MOR-*hmx2* mRNA ($96.2 \pm 1.5$) | *slc17a6a/b* | 10↓ | <0.001<sup>‡</sup> |
| 3S | Uninjected control ($106.1 \pm 0.8$) vs *hmx2;hmx3a* DKD + MOR-*hmx3a* mRNA ($91.5 \pm 1.6$) | *slc17a6a/b* | 15↓ | <0.001<sup>‡</sup> |
| 3S | *hmx2;hmx3a* DKD ($72.2 \pm 1.1$) vs *hmx2;hmx3a* DKD + MOR-*hmx2* mRNA ($96.2 \pm 1.5$) | *slc17a6a/b* | 24↑ | <0.001<sup>‡</sup> |
| 3S | *hmx2;hmx3a* DKD ($72.2 \pm 1.1$) vs *hmx2;hmx3a* DKD + MOR-*hmx3a* mRNA ($91.5 \pm 1.6$) | *slc17a6a/b* | 19↑ | <0.001<sup>‡</sup> |
| 3S | *hmx2;hmx3a* DKD + MOR-*hmx2* mRNA ($96.2 \pm 1.5$) vs *hmx2;hmx3a* DKD + MOR-*hmx3a* mRNA ($91.5 \pm 1.6$) | *slc17a6a/b* | 5↓ | 0.286<sup>‡</sup> |
| 3V | Uninjected control ($106.1 \pm 0.8$) vs *hmx2* SKD + MOR-*hmx2* mRNA ($105.6 \pm 1.7$) | *slc17a6a/b* | 1↓ | 0.997<sup>‡</sup> |
| 3V | Uninjected control ($106.1 \pm 0.8$) vs *hmx2* SKD + MOR-*hmx3a* mRNA ($105.6 \pm 1.8$) | *slc17a6a/b* | 0 | 0.997<sup>‡</sup> |
| 3V | *hmx2* SKD ($89.1 \pm 1.5$) vs *hmx2* SKD + MOR-*hmx2* mRNA ($105.6 \pm 1.7$) | *slc17a6a/b* | 17↑ | <0.001<sup>‡</sup> |
| 3V | *hmx2* SKD ($89.1 \pm 1.5$) vs *hmx2* SKD + MOR-*hmx3a* mRNA ($105.6 \pm 1.8$) | *slc17a6a/b* | 17↑ | <0.001<sup>‡</sup> |
| 3V | *hmx2* SKD + MOR-*hmx2* mRNA ($105.6 \pm 1.7$) vs *hmx2* SKD + MOR-*hmx3a* mRNA ($105.6 \pm 1.8$) | *slc17a6a/b* | 0 | 1<sup>‡</sup> |
| 3Y | Uninjected control ($106.1 \pm 0.8$) vs *hmx3a* SKD + MOR-*hmx2* mRNA ($95.0 \pm 1.5$) | *slc17a6a/b* | 11↓ | <0.001<sup>‡</sup> |
| 3Y | Uninjected control ($106.1 \pm 0.8$) vs *hmx3a* SKD + MOR-*hmx3a* mRNA ($102.4 \pm 1.5$) | *slc17a6a/b* | 4↓ | 0.345<sup>‡</sup> |
| 3Y | *hmx3a* SKD ($84.7 \pm 1.3$) vs *hmx3a* SKD + MOR-*hmx2* mRNA ($95.0 \pm 1.5$) | *slc17a6a/b* | 10↑ | <0.001<sup>‡</sup> |
| 3Y | *hmx3a* SKD ($84.7 \pm 1.3$) vs *hmx3a* SKD + *MOR-hmx3a* mRNA ($102.4 \pm 1.5$) | *slc17a6a/b* | 18↑ | **<0.001**<sup>‡</sup> |
| 3Y | *hmx3a* SKD + *MOR-hmx2* mRNA ($95.0 \pm 1.5$) vs *hmx3a* SKD + *MOR-hmx3a* mRNA ($102.4 \pm 1.5$) | *slc17a6a/b* | 7↑ | 0.063<sup>‡</sup> |

**Table 1B.** <u>Statistical comparisons of numbers of cells expressing particular genes in mutant experiments.</u> Statistical comparisons between WT sibling embryos and mutant embryos. First column indicates the figure panel that contains the relevant individual value plots for the comparison. NS = not shown in a figure. Second column states the genotypes being compared. Single mutant and double deletion embryos have to be obtained from different parents because *hmx2* and *hmx3a* are adjacent to each other on chromosome 17. Therefore, these were always analyzed as separate experiments and single mutants were only compared to double deletion mutants when there was no statistically significant difference between the WT sibling cell counts in the two experiments. Numbers within parentheses indicate mean numbers of cells ± S.E.M. In all cases, numbers are an average of at least 5 embryos and cells were counted in all dorsal-ventral spinal cord rows. All of the experiments were conducted on 27 h embryos except those indicated with (48 h) next to the gene name, which used 48 h embryos. 27 h embryos fixed on different days varied slightly in stage from prim stage 9 to 12. This explains the small differences in numbers of cells labelled with a particular probe in WT embryos in different experiments. Column three lists the gene that the cell counts and statistical comparison refer to. Asterisks indicate experiments performed with dextran sulfate (see materials and methods). The fourth column indicates the difference between the two mean values for the embryos being compared. All values are rounded to the nearest whole number. ↑ = increase, ↓ = decrease. Last column shows the P value for the comparison, rounded to three decimal places. Statistically significant (P<0.05) values are indicated in bold. Statistical test used is indicated by superscript symbol: Wilcoxon-Mann-Whitney test (^^^), type 2 Student’s t-test (^+^) or type 3 Student’s t-test (^§^). For a discussion of why particular tests were used, see materials and methods.

| Fig. | Comparison | Gene | Difference between two means | P value |
| --- | --- | --- | --- | --- |
| NS | WT ($105.5 \pm 2.2$) vs *hmx2*<sup>SU37</sup> ($103.3 \pm 0.7$) | *slc17a6a/b* | 2↓ | 0.582<sup>^</sup> |
| NS | WT ($102.8 \pm 0.9$) vs *hmx2*<sup>SU38</sup> ($103.6 \pm 1.8$) | *slc17a6a/b* | 1↓ | 0.700<sup>+</sup> |
| 5K | WT ($96.4 \pm 1.3$) vs *hmx2*<sup>SU39</sup> ($94.7 \pm 1.1$) | *slc17a6a/b* | 2↓ | 0.351<sup>+</sup> |
| 5G | WT ($106.0 \pm 0.9$) vs *hmx3a*<sup>SU42</sup> ($108.0 \pm 3.1$) | *slc17a6a/b* | 2↑ | 0.673<sup>^</sup> |
| 5H | WT ($102.2 \pm 1.1$) vs *hmx3a*<sup>sa23054</sup> ($101.0 \pm 1.7$) | *slc17a6a/b* | 1↓ | 0.567<sup>+</sup> |
| NS | WT ($97.0 \pm 1.2$) vs *hmx3a*<sup>SU43</sup> ($85.2 \pm 1.2$) | *slc17a6a/b* | 12↓ | **0.019**<sup>^</sup> |
| 5I | WT ($99.8 \pm 1.9$) vs *hmx3a*<sup>SU3</sup> ($86.4 \pm 0.8$) | *slc17a6a/b* | 13↓ | **<0.001**<sup>+</sup> |
| 5J | WT ($102.2 \pm 1.9$) vs *hmx2;hmx3a*<sup>SU44</sup> ($89.0 \pm 1.9$) | *slc17a6a/b* | 13↓ | **0.001**<sup>+</sup> |
| NS | WT ($102.0 \pm 1.7$) vs *hmx2;hmx3a*<sup>SU45</sup> ($87.8 \pm 2.2$) | *slc17a6a/b* | 14↓ | **<0.001**<sup>+</sup> |
| 5I&J | *hmx3a*<sup>SU3</sup> ($86.4 \pm 0.8$) vs *hmx2;hmx3a*<sup>SU44</sup> ($89.0 \pm 1.9$) | *slc17a6a/b* | 3↑ | 0.243<sup>+</sup> |
| NS | *hmx3a*<sup>SU3</sup> ($86.4 \pm 0.8$) vs *hmx2;hmx3a*<sup>SU45</sup> ($87.8 \pm 2.2$) | *slc17a6a/b* | 1↑ | 0.559<sup>§</sup> |
| NS | *hmx3a*<sup>SU43</sup> ($85.2 \pm 1.2$) vs *hmx2;hmx3a*<sup>SU44</sup> ($89.0 \pm 1.9$) | *slc17a6a/b* | 4↑ | 0.126<sup>+</sup> |
| NS | *hmx3a*<sup>SU43</sup> ($85.2 \pm 1.2$) vs *hmx2;hmx3a*<sup>SU45</sup> ($87.8 \pm 2.2$) | *slc17a6a/b* | 3↑ | 0.309<sup>§</sup> |
| 6K | WT ($69.6 \pm 0.7$) vs *hmx3a*<sup>SU3</sup> ($70.2 \pm 0.9$) | *hmx3a* | 1↑ | 0.613<sup>+</sup> |
| 6L | WT ($47.6 \pm 2.0$) vs *hmx3a*<sup>SU3</sup> ($45.0 \pm 1.7$) | *en1b* | 3↓ | 0.356<sup>+</sup> |
| 6M | WT ($154.4 \pm 2.9$) vs *hmx3a*<sup>SU3</sup> ($167.8 \pm 2.3$) | *slc32a1* | 13↑ | **0.004**<sup>^</sup> |
| 6N | WT ($158.0 \pm 2.0$) vs *hmx2;hmx3a*<sup>SU44</sup> ($169.2 \pm 1.7$) | *slc32a1* | 11↑ | **0.002**<sup>+</sup> |
| NS | WT ($158.0 \pm 1.3$) vs *hmx3a*<sup>SU43</sup> ($167.6 \pm 2.7$) | *slc32a1* | 10↑ | **0.012**<sup>+</sup> |
| 6O | WT ($130.4 \pm 1.4$) vs *hmx3a*<sup>SU3</sup> ($130.0 \pm 3.0$) | *slc6a5* | 0 | 0.907<sup>+</sup> |
| NS | WT ($111.8 \pm 3.9$) vs *hmx2;hmx3a*<sup>SU44</sup> ($118.3 \pm 3.0$) | *slc6a5* | 7↑ | 0.234<sup>+</sup> |
| 6AB | WT ($79.0 \pm 2.1$) vs *hmx3a*<sup>SU3</sup> ($90.4 \pm 2.4$) | *gad1b* | 11↑ | **0.022**<sup>^</sup> |
| 6AC | WT ($77.5 \pm 2.5$) vs *hmx2;hmx3a*<sup>SU44</sup> ($87.4 \pm 2.3$) | *gad1b* * | 10↑ | **0.014**<sup>+</sup> |
| 6AB&AC | *hmx3a*<sup>SU3</sup> ($90.4 \pm 2.4$) vs *hmx2;hmx3a*<sup>SU44</sup> ($87.4 \pm 2.3$) | *gad1b* * | 3↓ | 0.382<sup>+</sup> |
| 6AD | WT ($151.4 \pm 2.7$) vs *hmx3a*<sup>SU3</sup> ($175.7 \pm 3.5$) | *gad1b*<br>(48h)* | 24↑ | **<0.001**<sup>+</sup> |
| 6AE | WT ($143.5 \pm 6.7$) vs *hmx2;hmx3a*<sup>SU44</sup> ($170.2 \pm 8.5$) | *gad1b*<br>(48h)* | 27↑ | **0.024**<sup>+</sup> |
| 6AF | WT ($152.4 \pm 4.8$) vs *hmx2*<sup>SU39</sup> ($153.6 \pm 2.5$) | *gad1b*<br>(48h)* | 1↑ | 0.608<sup>^</sup> |
| 6AD&AE | *hmx3a*<sup>SU3</sup> ($175.7 \pm 3.5$) vs *hmx2;hmx3a*<sup>SU44</sup> ($170.2 \pm 8.5$) | *gad1b*<br>(48h)* | 5↓ | 0.562<sup>§</sup> |

### Microarray expression profiling experiments

These experiments are described in detail in (Cerda *et al*. 2009). P values were corrected for multiple testing (Benjamini and Hochberg 1995; Gentleman *et al*. 2004; Tarraga *et al*. 2008). These data have been deposited in the NCBI Gene Expression Omnibus with accession number GSE145916.

### Data and Reagent Availability

Plasmids and zebrafish strains are available upon request. Two supplemental figures, two supplemental tables and a supplementary materials reference list are available at FigShare. Figure S1 contains RT-PCR and cell-count data demonstrating the efficacy of *hmx2;hmx3a* DKD with splice-blocking morpholinos. Figure S2 shows an alignment of mouse and zebrafish Hmx2 and Hmx3a protein sequences. Table S1 includes gene names, ZFIN identifiers and references for *in situ* hybridization probes. Table S2 contains the sgRNA and primer sequences used for *hmx*2, *hmx3a* and *hmx2;hmx3a* CRISPR mutagenesis and genotyping. Microarray data have been previously deposited in the NCBI Gene Expression Omnibus with accession number GSE145916.

## Results

### *hmx2* and *hmx3a* are the only *hmx* genes expressed in spinal cord

While the expression and functions of zebrafish *hmx* genes have been analyzed during the development of sensory structures such as the eye and the ear, the expression of *hmx1*, *hmx2*, *hmx3a* and *hmx4* in the developing spinal cord has not been investigated and no expression data has previously been reported for *hmx3b,* which only appeared in more recent versions of the zebrafish genome sequence (Zv9 and above). Therefore, to determine which of the *hmx* genes are expressed in the spinal cord we performed *in situ* hybridizations for *hmx1*, *hmx2*, *hmx3a*, *hmx3b* and *hmx4* at different developmental stages (Fig. 1). At all of these stages, we observed no spinal cord expression of *hmx1*, *hmx3b* or *hmx4* (Fig. 1). However, consistent with previous reports, both *hmx1* and *hmx4* were expressed in the developing eye, ear and anterior lateral line neuromasts (Fig. 1; French *et al*. 2007; Feng and Xu 2010; Gongal *et al*. 2011; Boisset and Schorderet 2012; Marcelli *et al*. 2014). In contrast, the only expression of *hmx3b* that we observed was weak hindbrain expression at later stages of development (36-48 h; Fig. 1S’, X’ & AC’).

**Figure 1.**
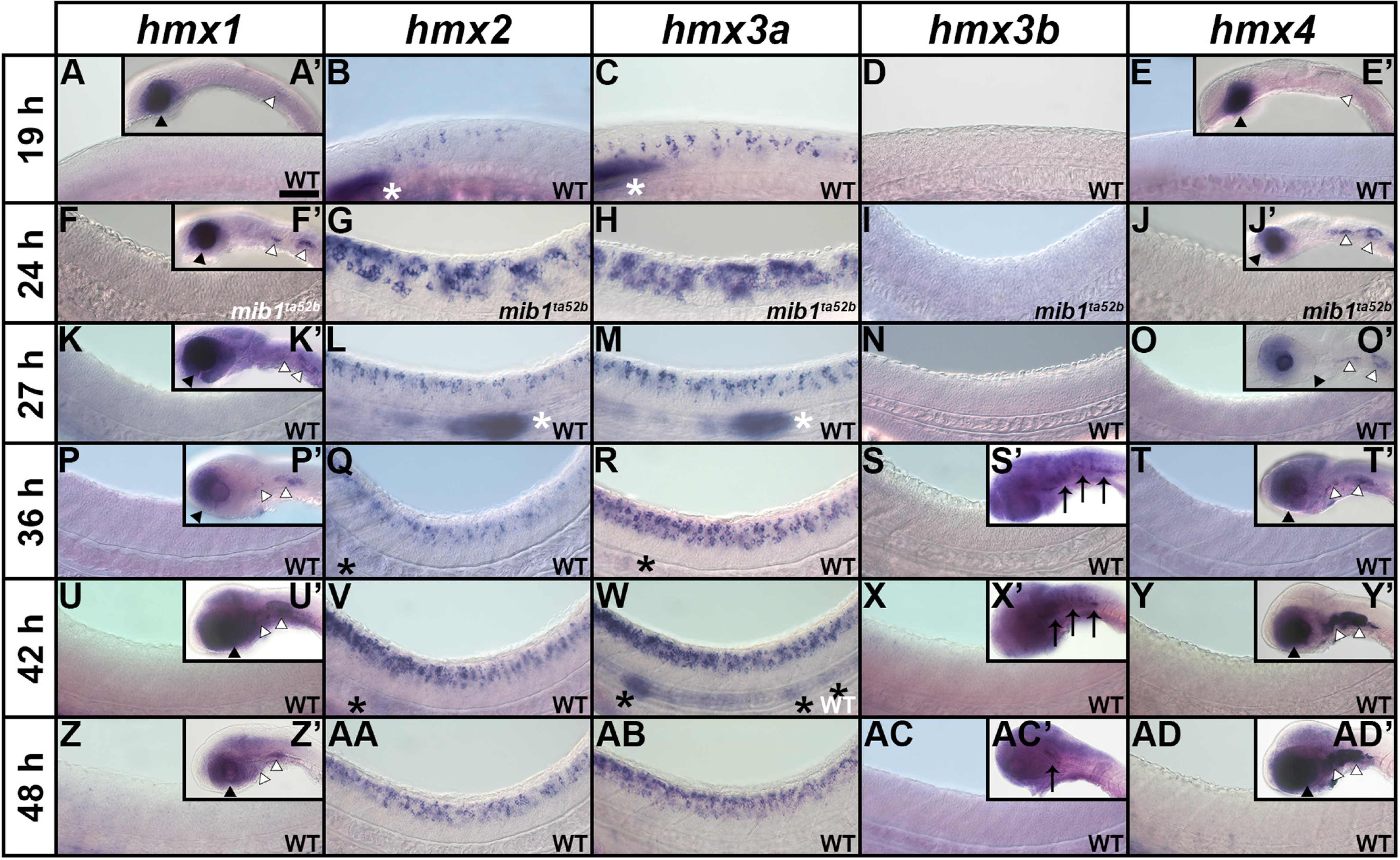
<u>Expression of *hmx* genes in WT zebrafish embryos</u>. Lateral views of *hmx* expression in spinal cord (**A-AD**), hindbrain (**S’, X’,AC’**), eye and ear (**A’, E’, F’, J’, K’, O’, P’, T’, U’, Y’, Z, AD’**) and lateral line primordium and neuromasts (**B-C, L-M, Q-R, V-W**) at 19 h (**A-E, A’, E’**), 24 h (**F-J, F’, J’**), 27 h (**K-O, K’, O’**), 36 h (**P-T, P’, S’, T’**), 42 h (**U-Y, U’, X’, Y’**) and 48 h (**Z-AD, Z’, AC’, AD’**). Rostral, left; Dorsal, up. (**B-C, G-H, L-M, Q-R, V-W, AA-AB**) *hmx2* and *hmx3a* are expressed in spinal cord, lateral line primordium (white asterisks), neuromasts (black asterisks), and anterior ear (data not shown) at all stages examined, although *hmx2* spinal cord expression initially appears weaker than *hmx3a* and does not extend as far caudally. Whilst there is expression of both *hmx2* and *hmx3a* in the lateral line primordium at 24 h (data not shown), the lateral line primordium has not yet migrated into the field of view shown in **G-H**. For consistency, the specific region of spinal cord shown (adjacent to somites 6-10) is identical in panels **F-AD**. At 19 h, expression is found only in the very anterior spinal cord and so a more rostral region of spinal cord is shown in **A-F**. (**A, A’, E, E’, F, F’, J, J’, K, K’, O, O’, P, P’, T, T’, U, U’, Y, Y’, Z, Z’, AD, AD’**) *hmx1* and *hmx4* are not expressed in WT spinal cord at any of these stages but are expressed in the eye (black arrowheads) and posterior-ventral ear and adjacent ganglion of the anterior lateral line (white arrowheads). (**D, I, N, S, S’, X, X’, AC, AC’**) *hmx3b* is not expressed in WT spinal cord at any of these stages. The only expression we observed was in the hindbrain between 36 and 48 h (black arrows). (**G, H**) The expression pattern of *hmx2* and *hmx3a* is expanded in the spinal cord of *mib1^ta52b^* mutants but is unaltered in the ear and lateral line primordium (data not shown). (**F, F’, J, J’**) Neither *hmx1* (**F**) nor *hmx4* (**J**) are expressed in the spinal cord of *mib1^ta52b^* mutants, although the expression of both genes persists in the eye (black arrowheads), posterior-ventral ear and adjacent ganglion of the anterior lateral line (white arrowheads) (**F’, J’**). (**I**) *hmx3b* is not expressed in *mib1^ta52b^* mutants – either in the spinal cord or in any other tissue. (**F, I, J, K, N, O, P, S, S’, T, U’, X, X’, Y, Z, AC, AC’, AD**) The background (diffuse, non-specific staining) in these pictures is higher because we exposed the embryos to prolonged staining to ensure that there was no weak spinal cord expression. Especially in the brain, this can lead to background staining as the large ventricles of the hindbrain trap anti-sense riboprobes. Scale Bar = 50 µm (**A-AD**), 120 µm (**A’, E’, F’, J’, K’, O’, P’, S’, T’, U’, X’, Y’, Z’, AC’, AD’**).

In contrast, *hmx2* and *hmx3a* are expressed in the spinal cord at all of the stages that we examined (Fig. 1). The spinal cord expression patterns of these two genes are very similar, with the exception that initially, *hmx2* expression appears to be weaker than *hmx3a* and it does not extend as far caudally (Fig. 1B & C). Consistent with previous reports, both of these genes are also expressed in the lateral line and developing ear (Fig. 1; Adamska *et al*. 2000; Feng and Xu 2010; Hartwell *et al*. 2019) as well as distinct regions of the brain (data not shown).

As *hmx3a* is expressed in the spinal cord, and teleost duplicate genes often have similar expression patterns, we wanted to further test whether there was any spinal cord expression of *hmx3b*. Therefore, we performed *in situ* hybridization on *mindbomb1 (mib1^ta52b^)* mutants at 24 h. *mib1* encodes an E3-ubiquitin protein ligase required for efficient Notch signaling. Consequently, Notch signaling is lost in *mib1^ta52b^* mutants and this causes most spinal progenitor cells to precociously differentiate into early-forming classes of spinal neurons at the expense of later-forming classes of neurons and glia (Jiang *et al*. 1996; Schier *et al*. 1996; Itoh *et al*. 2003; Park and Appel 2003; Batista *et al*. 2008). As a result, weak expression in spinal neurons is often expanded and stronger, and hence easier to observe, in 24 h *mib1^ta52b^* mutants (Batista *et al*. 2008; England *et al*. 2017). However, even in *mib1^ta52b^* mutants we detected no expression of *hmx3b* (Fig. 1I). We also analyzed the expression of the other *hmx* genes in 24 h *mib1^ta52b^* mutants. The expression patterns of both *hmx2* and *hmx3a* are expanded in the spinal cord of these mutants (Fig. 1G & H) but are unaltered in the ear and lateral line primordium (data not shown). The expanded spinal cord expression suggests that at least some of the spinal cord neurons expressing *hmx2* and *hmx3a* differentiate precociously in *mib1^ta52b^* mutants. In contrast, whilst the expression of *hmx1* and *hmx4* persists in the eye, posterior-ventral ear and adjacent ganglion of the anterior lateral line in *mib1^ta52b^* mutants (Fig. 1F’ & J’), we still did not observe any expression in the spinal cord (Fig. 1F & J).

### *hmx2* and *hmx3a* are expressed in V1 and dI2 interneurons in the spinal cord

To identify the spinal cord neurons that express *hmx2* and *hmx3a*, we performed several different double-labeling experiments. Double *in situ* hybridization with *hmx2* and *hmx3a* confirmed that these genes are co-expressed in the exact same cells in the spinal cord (Fig. 2A). Approximately half of these *hmx2* and *hmx3a (hmx2/3a)* co-expressing spinal cells also co-express *slc32a1*, which is only expressed by inhibitory (glycinergic and GABAergic) interneurons (Jellali *et al*. 2002), and approximately half co-express *slc17a6a/b,* which are only expressed by excitatory (glutamatergic) interneurons (Serrano-Saiz *et al*. 2013) (Fig. 2B & D; see materials and methods for a more detailed description of probes used to determine neurotransmitter phenotypes and additional references). In addition, the inhibitory *hmx2/3a*-expressing cells are generally more ventral than the excitatory double-labelled cells. Our previous expression-profiling of V1 interneurons, suggested that these cells might be the ventral inhibitory neurons that express *hmx3a* (Fig. 2G, for a description of these experiments see (Cerda *et al*. 2009)). Results from our lab and others, have established that V1 interneurons are the only spinal cord cells that express *engrailed1b* (*en1b*) (Higashijima *et al*. 2004a; Batista and Lewis 2008). Therefore, to confirm that V1 interneurons also express *hmx3a* we performed double *in situ* hybridizations for *hmx3a* and *en1b*. These experiments showed that all of the *en1b*-expressing spinal cells co-express *hmx3a,* and that approximately half of the *hmx2/3a*-expressing spinal cells co-express *en1b* (Fig. 2C). Taken together, these data clearly identify the inhibitory *hmx2/3a*-expressing spinal cells as V1 interneurons.

**Figure 2.**
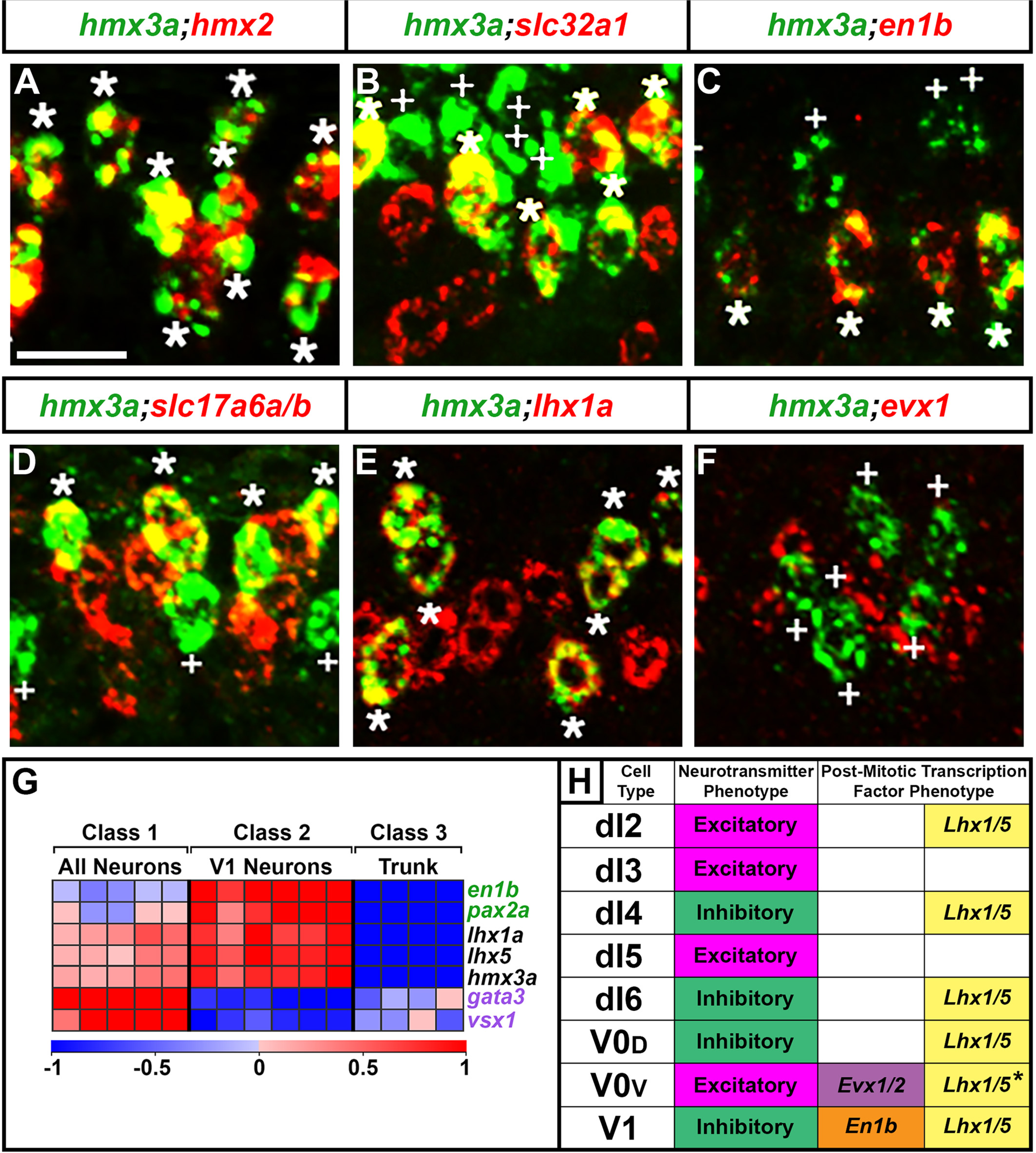
*<u>hmx2</u>* <u>and *hmx3a* are expressed in V1 and dI2 interneurons.</u> (**A-F**) Lateral views of spinal cord at 27 h. Rostral, left; Dorsal, up. *hmx2* and *hmx3a* are co-expressed (A) in V1 and dI2 interneurons. (**B-F**) V1 interneurons are inhibitory (*slc32a1*-expressing) (Jellali *et al*. 2002; Goulding *et al*. 2014) (**B**) and express *en1b* (**C**). dI2 interneurons are glutamatergic (excitatory, express *slc17a6a/b*) (Alaynick *et al*. 2011; Serrano-Saiz *et al*. 2013) (**D**) and express *lhx1a* (**E**) but not *evx1* (**F**). Asterisks = double-labeled and white crosses = single-labeled *hmx3a*-expressing cells (green). Expression of other genes is red. Scale bar = 20 µm. (**G**) Heatmap analysis of gene expression profiling of V1 interneurons. A three-class ANOVA analysis of differential expression was performed on different FAC-sorted populations of cells. Class 1: All post-mitotic spinal neurons. Class 2: V1 interneurons. Class 3: All trunk cells. Each column is a different biological replicate. Rows show relative expression levels for a single transcription factor gene as normalized data transformed to a mean of 0, with standard deviation of +1 (highly expressed) or -1 (weakly/not expressed) sigma units. Adjusted P values corrected for multiple testing are <0.000001 for all genes shown. For more information on these experiments see (Cerda *et al*. 2009). Expression profiles are included for positive control genes (green), *en1b* and *pax2a,* that are expressed by V1 cells and negative control genes (purple), *gata3* and *vsx1,* that are expressed by other spinal cord interneurons but not V1 cells. Data for *lhx1a, lhx5* and *hmx3a* shows that they are co-expressed in V1 interneurons. (**H**) Schematic showing neurotransmitter and post-mitotic transcription factor phenotypes of spinal cord interneurons found in the dorsal-ventral spinal cord region shown in panels A-F. It is currently unclear whether V0v interneurons express Lhx1/5 (*, see discussion in the results).

As mentioned above, the glutamatergic *hmx2/3a*-expressing cells are generally located more dorsal to the inhibitory *hmx2/3a*-expressing cells. Therefore, these excitatory cells could be V0v, dI5, dI3, dI2 or dI1 interneurons (Fig. 2; Cheng *et al*. 2005; Grossmann *et al*. 2010; Satou *et al*. 2012; Talpalar *et al*. 2013). The zebrafish embryonic spinal cord is relatively small. For example, at 27h, the dorsal-ventral axis is only about 10 cells high. As a result, the different neuronal populations are often intermingled, rather than clearly separated as they are in amniotes (e.g. see Batista *et al*. 2008; England *et al*. 2011). In addition, studies in amniotes suggest that many dorsal neurons migrate dorsally or ventrally soon after they are born (e.g. see Gross *et al*. 2002; Müller *et al*. 2002). Taken together, this means that it is hard to accurately identify dorsal spinal cord cell types by position alone. Therefore, to identify the excitatory *hmx2/3a*-expressing neurons we performed double labeling experiments with markers of different dorsal excitatory cell types. We found that all *hmx2/3a*-expressing spinal cells co-express *lhx1a* and *lhx5* (Fig. 2E & G). Lhx1a and Lhx5 are predominantly expressed by inhibitory spinal cord interneurons, but they are also expressed by dI2 interneurons, which are excitatory (Gowan *et al*. 2001; Moran-Rivard *et al*. 2001; Gross *et al*. 2002; Muller *et al*. 2002; Cheng *et al*. 2004; Wine-Lee *et al*. 2004; Muller *et al*. 2005; Alaynick *et al*. 2011; Satou *et al*. 2012). The only other excitatory neurons that might express these two *lhx* genes are V0v interneurons. Recent single-cell RNA-Seq (scRNA-Seq) data from mouse spinal cord identified possible V0 cells that expressed *lhx1a* and *lhx5*, although, given the relatively small size of this recovered population, it is not clear whether these cells were excitatory V0v and/or inhibitory V0_D_ cells (Delile *et al*. 2019). V0v cells are the only spinal cord cells that express *evx1* and *evx2* (Juárez-Morales *et al*. 2016 and references therein). Therefore, we tested whether there was any co-expression of *hmx3a* and *evx1/2* using double *in situ* hybridization as well as immunohistochemistry for GFP and *in situ* hybridization for *hmx3a* in *Tg*(*evx1:EGFP*)*^SU1^* and *Tg*(*evx1:EGFP*)*^SU2^* embryos. However, we did not observe any co-expression in any of these experiments (Fig. 2F and data not shown). Therefore, we are confident that the excitatory *hmx2/3a*-expressing spinal cells are dI2 interneurons. Consistent with this, recent mouse scRNA-Seq spinal cord data suggests that mouse *Hmx2* and *Hmx3* are also expressed in V1 and dI2 spinal cord interneurons (Delile *et al*. 2019).

### Knock-down experiments suggest that *hmx2* and *hmx3a* may be redundantly required for correct specification of a subset of spinal interneuron glutamatergic phenotypes

As an initial step to try and identify the function(s) of *hmx2* and *hmx3a* in spinal cord development we performed morpholino knock-down experiments. As previous analyses suggested that these genes have redundant roles in ear and lateral line development (Feng and Xu 2010), we designed and injected translation-blocking morpholinos for both of these genes (see materials and methods). In embryos co-injected with the two translation-blocking morpholinos (*hmx2;hmx3a* double knock-down (DKD) animals), we observed stalled lateral line progression and fused otoliths in the ears (Fig. 3A-D). Normally there is an anterior (utricular) and a posterior (saccular) otolith in each ear, but in DKD embryos there was just one fused otolith in a medio-ventral region of each ear (Fig 3D). When we analyzed spinal cord phenotypes, we detected no change in the number of *hmx3a*- or *en1b*-expressing cells in DKD embryos, suggesting that dI2 and V1 interneurons still form in normal numbers (Fig. 3E-J, Table 1A). However, when we examined markers of neurotransmitter phenotypes, we observed a reduction in the number of spinal excitatory (glutamatergic, *slc17a6-*expressing) cells and a corresponding increase in inhibitory (*slc32a1*-expressing) cells (Fig. 3K-P, Table 1A). As *hmx2* and *hmx3a* are only expressed by dI2 neurons (which are glutamatergic) and V1 neurons (which are inhibitory) in the spinal cord, this suggested that at least some dI2 interneurons had switched their neurotransmitter phenotype from glutamatergic to inhibitory. Consistent with the idea that the two genes act redundantly, both of these spinal cord phenotypes were less severe in SKD embryos (Fig. 3M, P, Table 1A). Interestingly, we also did not see abnormal lateral line progression or ear phenotypes in SKD embryos.

**Figure 3.**
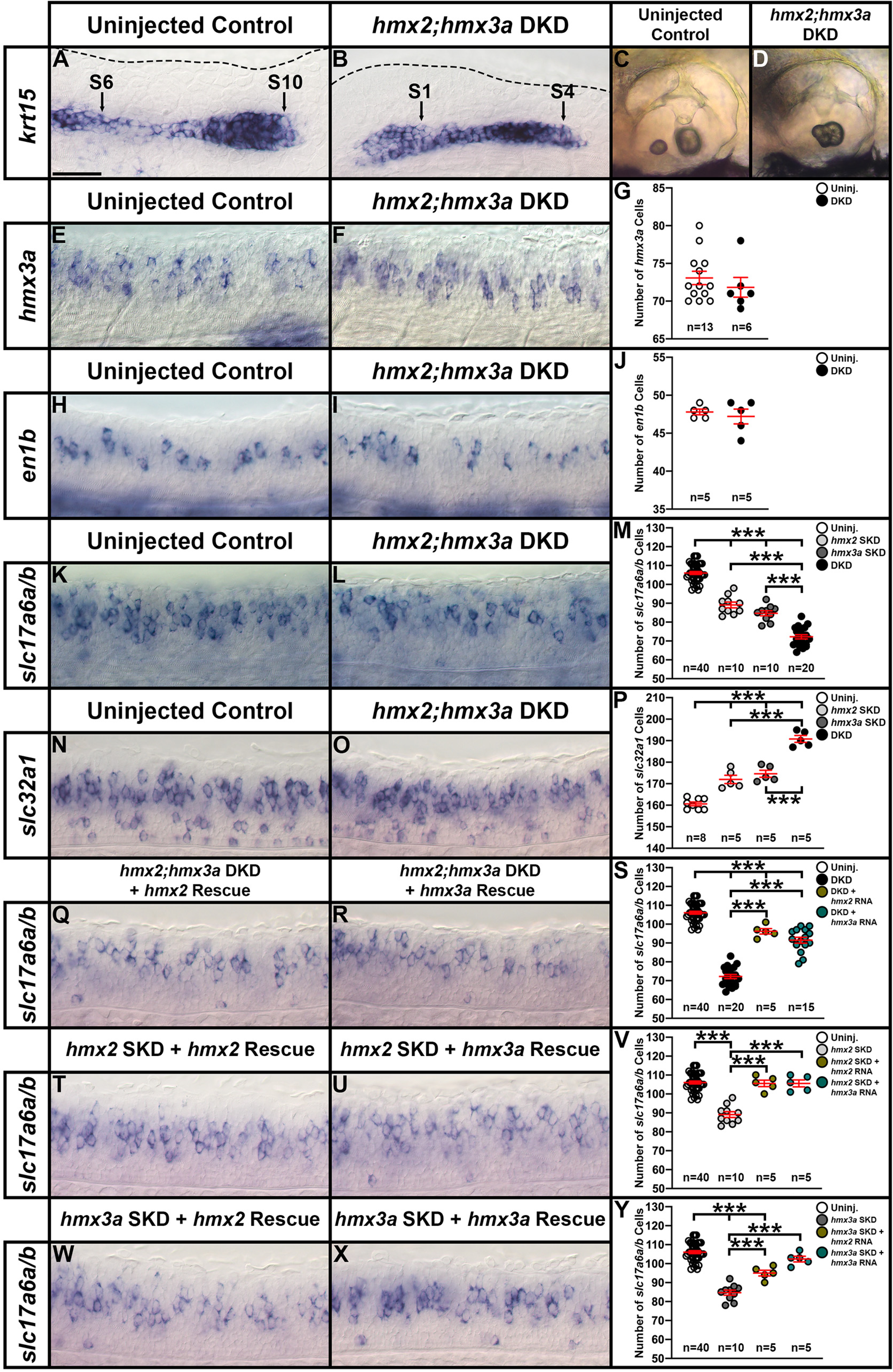
*<u>hmx2/3a</u>* <u>double knockdown (DKD) embryos have fewer excitatory (glutamatergic) and more inhibitory spinal cord interneurons.</u> (**A-D, E-F, H-I, K-L, N-O, Q-R, T-U, W-X**) Lateral views of (**A-B, E-F, H-I, K-L, N-O, Q-R, T-U, W-X**) spinal cord at 27 h and (**C-D**) otic vesicles at 3 d. Rostral, left; Dorsal, up. (**G, J, M, P, S, V, Y**) Mean number of cells expressing *hmx3a* (**G**), *en1b* (**J**), *slc17a6a/b* (**M, S, V, Y**) and *slc32a1* (**P**) in a precisely-defined spinal cord region adjacent to somites 6-10 at 27 h. All counts are an average of at least 5 embryos. Data are depicted as individual value plots and the *n*-values for each genotype are also shown. For each plot, the wider red horizontal bar depicts the mean number of cells and the red vertical bars depict the standard error of the mean (standard error of the mean (S.E.M.) values are listed in Table 1A). Statistically significant (*p* < 0.001) comparisons are indicated with brackets and three asterisks. All data were first analyzed for normality using the Shapiro-Wilk test. Data set in **G** is non-normally distributed and was, therefore, analyzed with the Wilcoxon-Mann-Whitney test. Data sets in **J, M, P, S, V** and **Y** are normally distributed and so for the pairwise comparison shown in J, the F test for equal variances was performed. This data set has equal variances and so a type 2 (for equal variances) student’s t-test was performed. To accurately compare the 4 different data sets each shown in panel **M, P**, **S, V** and **Y**, a one-way analysis of variance (ANOVA) test was performed. All data sets for ANOVA analysis have both normal distributions and homogeneous (homoscedastic, Bartlett’s test p value >0.05) variances and so standard ANOVA analysis was performed. All ANOVA analyses shown are significant (**M**: ANOVA (*F*(3,76) = 231.5, *p* = <0.0001), **P**: ANOVA (*F*(3,19) = 80.64, *p* = <0.0001), **S**: ANOVA (*F*(3,76) = 196.3, *p* = <0.0001), **V**: ANOVA (*F*(3,56) = 34.97, *p* = <0.0001) and **Y**: ANOVA (*F*(3,56) = 61.14, *p* = <0.0001)), and so to determine which specific experimental groups or groups differed, Tukey’s honestly significant difference post hoc test for multiple comparisons was performed. Mean numbers of cells and P-values are provided in Table 1A. In some cases, cell count data were pooled from different experiments (uninjected control data in Fig. 3M, S, V and Y from 12 pooled experiments, *hmx2;hmx3a* DKD data in Fig. 3M & S from 4 pooled experiments, *hmx2* SKD data in Fig. 3M & V from 2 pooled experiments, and *hmx3a* SKD data in Fig. 3M & Y from 2 pooled experiments). As *in situ* hybridization staining can vary slightly between experiments, we only pooled data from different experiments, or compared different morpholino-injection experiments if pairwise comparisons of the counts from corresponding uninjected WT control embryos were not statistically significantly different from each other. (**A-B**) By 27 h, in an uninjected WT control embryo *krt15* mRNA expression shows that the lateral line primordium (LLP) has migrated to its expected position over somite 10 (S10 + black arrow) (**A**). In contrast, at 27 h in *hmx2;hmx3a* DKD embryos, the LLP is stalled beside somites 1-4 (S1, S4, black arrows, **B**). This is identical to the stalled LLP phenotype observed in *hmx3a^SU3^* and *hmx2;hmx3a^SU44^* mutants (see Fig. 5O-P). Dotted line indicates dorsal spinal cord boundary in **A** and dorsal posterior hindbrain and anterior spinal cord boundary in **B**. (**C-D**) Also like *hmx3a^SU3^* and *hmx2;hmx3a^SU44^* mutants (see Fig. 5U-V), *hmx2;hmx3a* DKD embryos have fused otoliths at 3 d (**D**), but uninjected controls do not (**C**). (**E-J**) There is no change in the number of *hmx3a-* or *en1b*-expressing spinal cells in *hmx2/3a* DKD compared to uninjected control embryos, suggesting that V1 and dI2 interneurons do not die or transfate/change into different cell types. (**K-M**) The number of *slc17a6a/b*- expressing (excitatory) spinal cells is reduced in both double and single knockdown (SKD) embryos, with the reduction being more severe in DKD embryos. (**N-P**) Concomitantly, there is a statistically significant increase in the number of *slc32a1*-expressing (inhibitory) cells in both SKD and DKD embryos, with the increase being more profound in DKD embryos. (**Q-S**) Injection of either morpholino-resistant *hmx2* (**Q**) or *hmx3a* mRNA (**R**) partially rescues the number of spinal excitatory cells in DKD embryos. (**T-V**) Injection of either morpholino-resistant *hmx2* (**T**) or *hmx3a* mRNA (**U**) fully rescues the number of spinal excitatory cells in *hmx2* SKD embryos. (**W-Y**) Injection of morpholino-resistant *hmx2* mRNA (**W**) partially rescues the number of spinal excitatory cells in *hmx3a* SKD embryos, but injection of morpholino-resistant *hmx3a* mRNA (**X**) fully rescues the phenotype. Scale bar = 30 µm (**A-B, E-F, H-I, K-L, N-O, Q-R, T-U, W-X**); 80 µm (**C-D**).

To confirm the specificity of these morpholino knock-down results, we first tested whether we saw a similar spinal cord phenotype if we injected splice-blocking morpholinos against *hmx2* and *hmx3a* (see materials and methods). In these experiments we obtained a partial reduction in the correct splicing of these genes (Fig. S1A-D) and a statistically significant reduction in the number of glutamatergic spinal cord cells (Fig S1E and see details in the figure legend). The reduction in the number of glutamatergic cells was less than for the translation-blocking DKD experiments, (14 cells vs 34 cells, see Figs. 3M & S1E, Table 1A), consistent with the fact that we only obtained a partial knock-down of each gene using the splice-blocking morpholinos. We then tested whether co-injecting a morpholino-resistant *hmx2* or *hmx3a* mRNA with the translation-blocking morpholinos could rescue the reduction in the number of spinal glutamatergic cells that occurs in SKD and DKD embryos (see materials and methods for the design of the mRNAs). We found that both *hmx3a* and *hmx2* morpholino-resistant mRNA could completely rescue the translation-blocking morpholino phenotype in *hmx2* SKD embryos (Fig. 3T-V and Table 1A) and *hmx3a* could completely rescue and *hmx2* could partially rescue the translation-blocking morpholino phenotype in *hmx3a* SKD embryos (Fig. 3W-Y and Table 1A). In addition, either *hmx2* or *hmx3a* morpholino-resistant mRNA was able to partially rescue the number of glutamatergic spinal neurons in DKD embryos (Fig. 3Q-S and Table 1A). Injections of higher amounts of mRNA or of both mRNAs at the same time led to embryo death, probably because of the toxic effects of injecting considerable amounts of both mRNA and morpholinos into the embryos during early development.

### Mutational analyses suggest that *hmx2* is not, by itself, required, and that Hmx3a protein may not require its DNA-binding homeodomain, for viability, correct migration of lateral line primordium, or correct development of ear otoliths or a subset of spinal cord interneurons

To further and more robustly test the hypothesis that *hmx2* and *hmx3a* are required for the correct specification of a subset of spinal cord interneuron neurotransmitter phenotypes, we created CRISPR mutants in each of these genes, targeting a region upstream of the homeobox (see materials and methods; Fig. 4). We also obtained a *hmx3a^sa23054^* allele from the Sanger zebrafish mutation project (Kettleborough *et al*. 2013) that introduces a stop codon upstream of the homeobox.

Our analyses of homozygous mutant embryos demonstrate that *hmx3a^SU3^* and *hmx3a^SU43^* mutants have fused otoliths, stalled lateral line progression and are homozygous lethal (Fig. 4, Fig. 5O, U, Table 2 and data not shown, also see (Hartwell *et al*. 2019) for a detailed description of the *hmx3a^SU3^* ear phenotype). When we examined the spinal cords of these mutants, we observed a statistically significant reduction in the number of glutamatergic cells, but in both cases the reduction was smaller than we had previously observed for morpholino-injected DKD embryos (Fig. 5D & I, Table 1B). There was also an increase in the number of inhibitory spinal cord interneurons, although, again, the increase was less than in the DKD morpholino-injected embryos (Fig. 6C, H, M, Table 1B). However, similar to the DKD embryos, there was no change in the number of spinal *hmx3a*- or *en1b*-expressing cells in *hmx3a^SU3^* mutants, suggesting that dI2 and V1 interneurons are forming in normal numbers and not dying or changing into different classes of interneurons (Fig. 6A, B, F, G, K, L, Table 1B).

**Table 2.** Statistical analyses of whether the frequencies of homozygous mutants that survive to adulthood or have abnormal otolith development or lateral line progression phenotypes are Mendelian. Column one indicates the mutant allele. These are listed in the same order as in Fig. 4. Columns two, four and six show the frequency of embryos with otolith fusion phenotypes from incrosses of heterozygous (z) or homozygous (mz) parents, the frequency of viable adult homozygous mutants and the frequency of embryos with stalled lateral line progression respectively. n = total number of animals examined. Columns three, five and seven list the P value, rounded up to 3 decimal places, for a Chi-squared test of the hypothesis that the frequency of embryos with fused otoliths, homozygous mutant adults that are viable or embryos with stalled lateral line progression respectively is Mendelian (25% for crosses of heterozygous parents (z) and 100% for crosses of homozygous parents (mz)). Statistically significant values are indicated in bold. N/A = not applicable (cases where no homozygous mutants survived, had otolith fusions or stalled lateral line phenotypes).

| **Mutant allele** | **% Otolith fusion phenotype** | **P value for otolith fusion phenotype** | **% Survival** | **P value for survival** | **% Stalled lateral line progression** | **P value for stalled lateral line progression** |
| --- | --- | --- | --- | --- | --- | --- |
| *SU35* | 0% (n=100; z) | N/A | 18.03% (n=61) | 0.234 | 0% (n=28) | N/A |
| *SU36* | 0% (n=100; z) | N/A | 27.27% (n=66) | 0.566 | 0% (n=26) | N/A |
| *SU37* | 0% (n=112; z) | N/A | 20.19% (n=104) | 0.258 | 0% (n=48) | N/A |
| *SU38* | 0% (n=56; z)<br>0% (n=77; mz) | N/A | 33.30% (n=54) | 0.214 | 0% (n=104) | N/A |
| *SU39* | 0% (n=49; z)<br>0% (n=228; mz) | N/A | 30.26% (n=76) | 0.289 | 0.69% (n=145) | **<0.001** |
| *sa23054* | 12.35% (n=332; z)<br>27.41% (n=135; mz) | **<0.001**<br>0.552 | 20.78% (n=77) | 0.428 | 0% (n=124) | N/A |
| *SU42* | 0% (n=121; z) | N/A | 20.00% (n=65) | 0.388 | 0% (n=104) | N/A |
| *SU3* | 24.85% (n=2370; z) | 0.850 | NONE | N/A | 20.36% (n=334) | 0.935 |
| *SU43* | 23.33% (n=463; z) | 0.391 | NONE | N/A | 22.94% (n=109) | 0.995 |
| *SU44* | 25.41% (n=1334; z) | 0.752 | NONE | N/A | 25.23% (n=329) | 1.000 |
| *SU45* | 29.52% (n=227; z) | 0.126 | NONE | N/A | 19.44% (n=108) | 0.879 |

**Figure 4.**
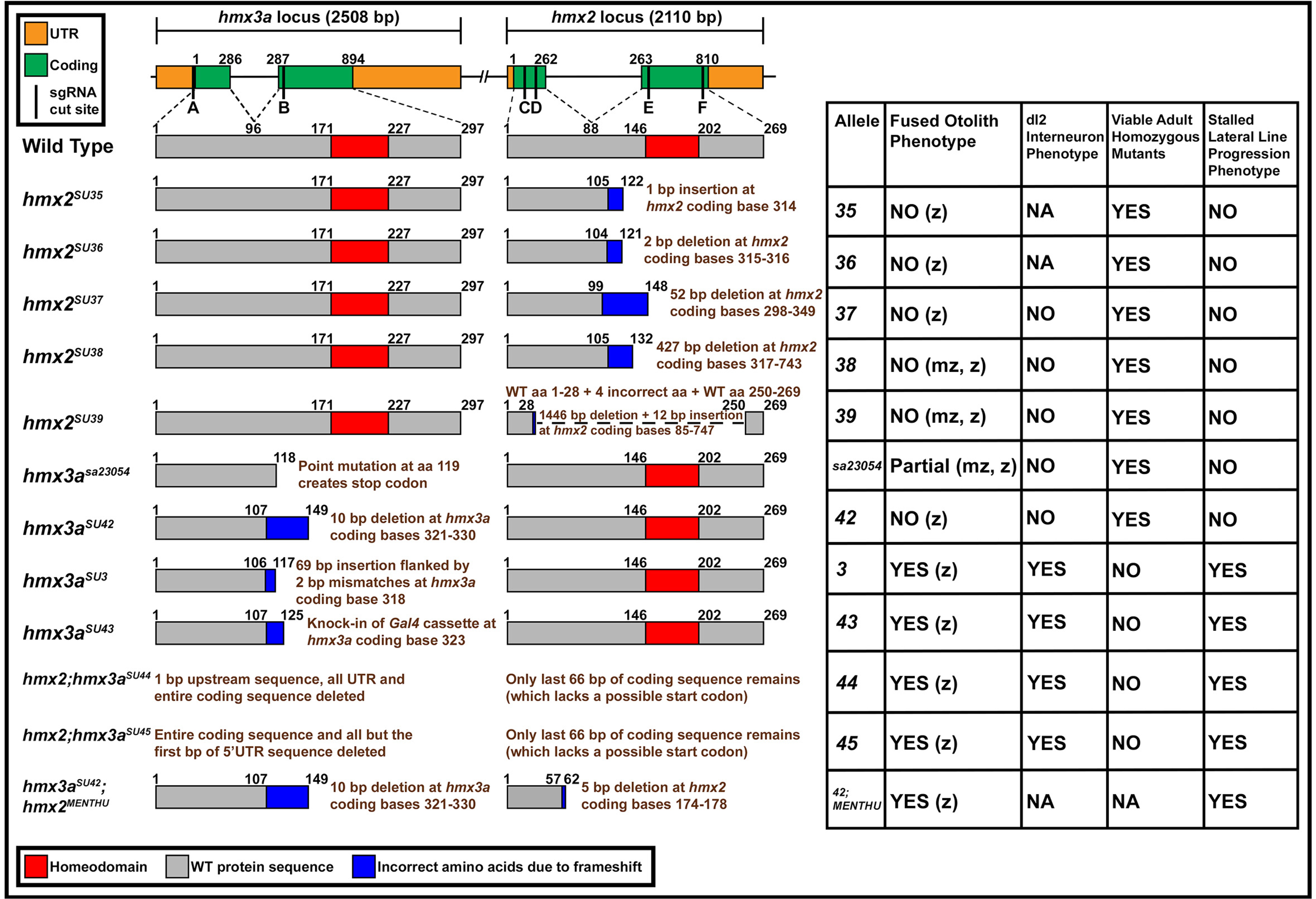
<u>Summary of *hmx2/3a* mutant alleles analyzed and their phenotypes when homozygous.</u> Left-hand side: schematics of 11 mutant alleles and one double mutant analyzed. Top row = genomic locus; lower rows indicate predicted protein products. There is a 6454 bp gap between *hmx2* and *hmx3a*. Vertical black bars on genomic locus indicate locations of sgRNA sequences, A-F, used to generate the mutants shown. These sequences and the combinations of sgRNAs used to generate the mutants shown here, are listed in Table S2. For each mutant allele, the genomic location plus the nature of the mutation or indel size is shown in brown text at the right-hand side of each mutant protein schematic. Coding bases refer to the translated sequence, e.g. coding bases 1-3 correspond to the bases encoding the start methionine. Right-hand side: Column 1: allele number. Column 2: indicates whether embryos with fused otoliths were observed in incrosses of heterozygous (z) or homozygous (mz) parents (also see Table 2 and Fig. 5). Column 3 indicates whether a reduction in the number of spinal excitatory cells was observed at 27 h in homozygous mutants, as assayed by *in situ* hybridization for *slc17a6a/b* (Fig. 5 and data not shown). Column 4: indicates whether viable adult homozygous mutants were recovered. In all cases where adult homozygous mutants were identified, the numbers of these fish are not statistically significantly different (P> 0.214) from expected mendelian ratios, as assayed by a Chi-squared test. P values are provided in Table 2. Column 5: indicates whether embryos with stalled lateral line progression phenotypes were observed in incrosses of heterozygous parents. All *hmx2* stable mutant alleles recovered to date are homozygous viable and homozygous mutants do not have a reduction in the number of glutamatergic spinal cord interneurons or otolith or stalled lateral line progression phenotypes. This is the case, even for embryos from incrosses of fish homozygous mutant for the most severely deleted *hmx2* alleles (*hmx2^SU38^* and *hmx2^SU39^*). *hmx3a^sa23054^* mutants are also homozygous viable. They have variable, incompletely penetrant, otolith fusion phenotypes. Strikingly, only 27.41% of embryos from an incross of homozygous mutant parents have otolith fusion phenotypes (Table 2). *hmx3a^sa23054^* mutants do not have a reduction in the number of glutamatergic spinal cord interneurons or stalled lateral line progression phenotypes. *hmx3a^SU42^* mutants are homozygous viable and do not have any obvious abnormal phenotypes, even though this allele should encode a protein with the same number of WT amino acids as *hmx3a^SU43^* and only one more WT amino acid than *hmx3a^SU3^*. *hmx2;hmx3a^SU44^* and *hmx2;hmx3a^SU45^* differ only in the amount of upstream sequence that is deleted and have identical phenotypes.

**Figure 5.**
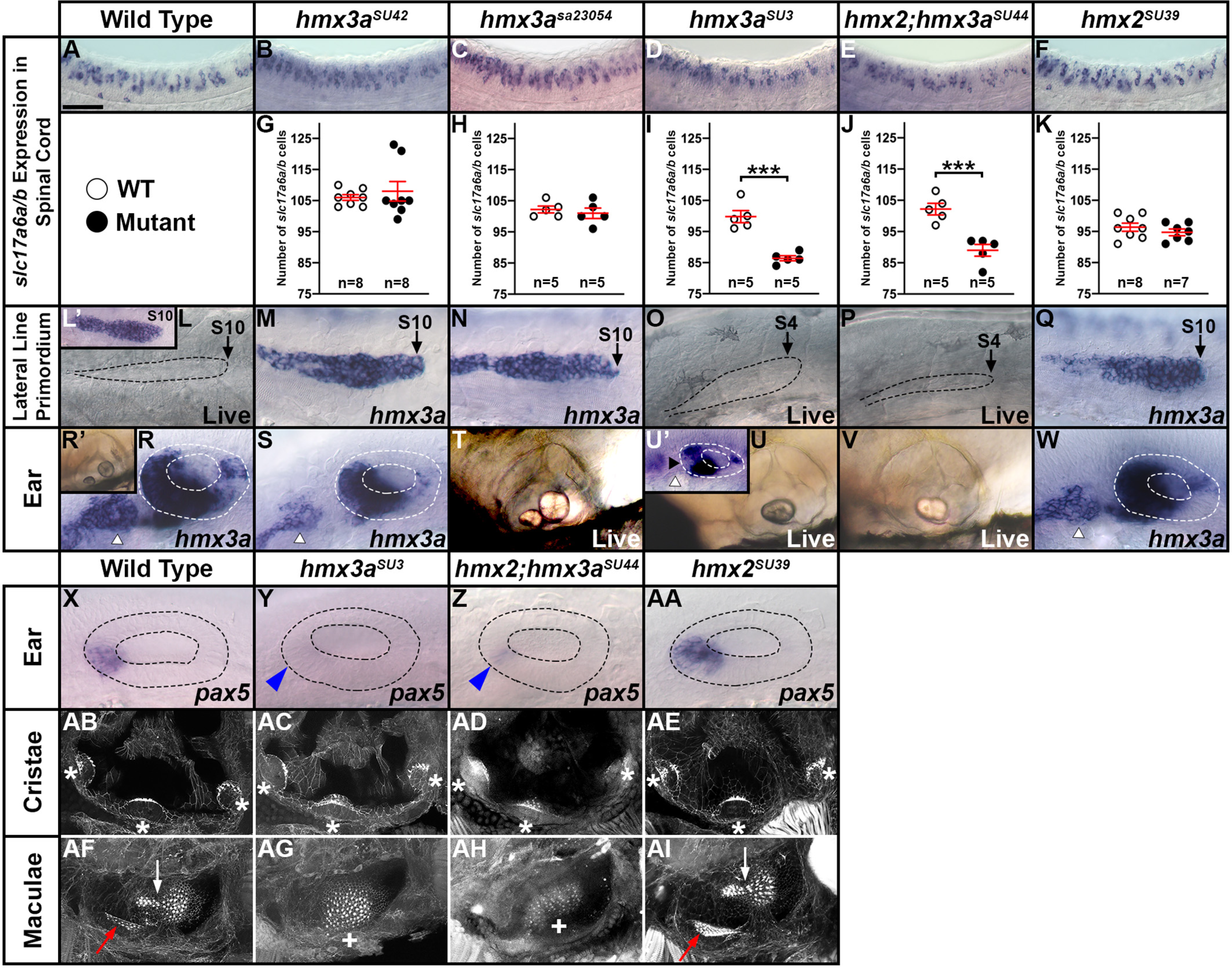
<u>Only some *hmx2/3a* alleles have mutant phenotypes.</u> (**A-F, L-AI, L’, R’ U’**) Lateral views. Rostral, left; Dorsal, up. (**A-F**) Expression of *slc17a6a/b* in spinal cord at 27 h. (**G-K**) Number of cells expressing *slc17a6a/b* in a precisely-defined spinal cord region adjacent to somites 6-10 at 27 h. Data are depicted as individual value plots and the *n*-values for each genotype are also shown. For each plot, the wider red horizontal bar depicts the mean number of cells and the red vertical bars depict the standard error of the mean (standard error of the mean (S.E.M.) values are listed in Table 1B). All counts are an average of at least 5 embryos. Statistically significant (*p* < 0.001) comparisons are indicated with brackets and three asterisks. White circles indicate WT data and black circles the appropriate mutant data as indicated in key under panel A. All data were first analyzed for normality using the Shapiro-Wilk test. Data in **G** is not normally distributed and so a Wilcoxon-Mann-Whitney test was performed. Data sets in **H-K** are normally distributed and so the F test for equal variances was performed, followed by a type 2 student’s t-test (for equal variances). P-values are provided in Table 1B. (**L-Q, L’**) Lateral line primordium phenotypes examined either by *hmx3a* expression (**L’, M-N, Q**) or live (**L, O-P**) at 27 h. (**R-AI, R’, U’**) Ear phenotypes examined either by *hmx3a* expression at 27 h (**R-S, U’ & W**), live at 4 d (**R’, T-V**), *pax5* expression at 24 h (**X-AA**) or phalloidin staining at 4 d (**AB-AI**). (**A-K**). There is no change in the number of spinal excitatory neurons in *hmx3a^SU42^* (**B, G**) and *hmx3a^sa23054^* (**C, H**) and *hmx2^SU39^* (**F, K**) mutant embryos compared to WT (**A**). In contrast, there is a statistically significant reduction in *hmx3a^SU3^* (**D, I**) and *hmx2;hmx3a^SU44^* (**E, J**) mutant embryos. (**L-Q, L’**). At 27 h, the tip of the lateral line primordium (black dotted line, **L**) has reached somite 10 (S10 = somite 10) in WT embryos (**L, L’**). This rate of migration is unchanged in *hmx3a^SU42^*, *hmx3a^sa23054^* and *hmx2^SU39^* mutant embryos (**M-N, Q**). (**O-P**) In contrast, the lateral line primordium fails to migrate in *hmx3a^SU3^* (**O**) and *hmx2;hmx3a^SU44^* (**P**) mutant embryos. Instead, it is stalled adjacent to somites 1-4 (S4 = somite 4). (**R-S, U’ & W**) *hmx3a* expression in the ear (inside white dotted lines) and in presumptive neuroblasts anterior to the ear (white arrowheads) is unchanged in *hmx3a^SU42^* (**S**) and *hmx2^SU39^* (**W**) mutants, compared to WT embryos (**R**), but is severely reduced in both the presumptive neuroblasts anterior to the ear and the anterior ear (black arrowhead) in *hmx3a^SU3^* mutants (**U’**). (**T**) *hmx3a^sa23054^* mutants show incompletely penetrant, variable otolith fusion phenotypes, ranging from no fusion (like WT ear in **R’**), through incomplete fusion (**T**), to complete fusion, like that observed at full penetrance in *hmx3a^SU3^* (**U**) and *hmx2;hmx3a^SU44^* (**V**) mutant embryos. (**X-AA**) The expression of *pax5* in the anterior ear (inside black dotted lines) is unchanged in *hmx2^SU39^* (**AA**) mutants, compared to WT embryos (**X**), but is severely reduced (blue arrowhead) in both *hmx3a^SU3^* (**Y**) and *hmx2;hmx3a^SU44^* (**Z**) mutants. (**AB-AI**) The three cristae of the ear (white asterisks) form normally in WT (**AB**), *hmx3a^SU3^* (**AC**), *hmx2;hmxsa^SU44^* (**AD**) and *hmx2^SU39^* (**AE**) mutants. In contrast, the spatially distinct anterior (utricular, red arrow) and posterior (saccular, white arrow) maculae are unchanged in *hmx2^SU39^* (**AI**) mutants, compared to WT embryos (**AF**), but are fused and are located in a more medio-ventral position (white cross) in *hmx3a^SU3^* (**AG**) and *hmx2;hmx3a^SU44^* (**AH**) mutants. However, there are no obvious differences between *hmx3a^SU3^* and *hmx2;hmx3a^SU44^* mutants. Scale bar = 50 µm (**A-F**), 30 µm (**L-W**), 20 µm (**X-AA**), 60 µm (**L’, R’, U’, AB-AI**). Panels **X-Y, AB-AC** and **AF-AG** are reproduced from (Hartwell *et al*. 2019) as per the Creative Commons Attribution (CC BY) license at *PLoS Genetics*.

**Figure 6.**
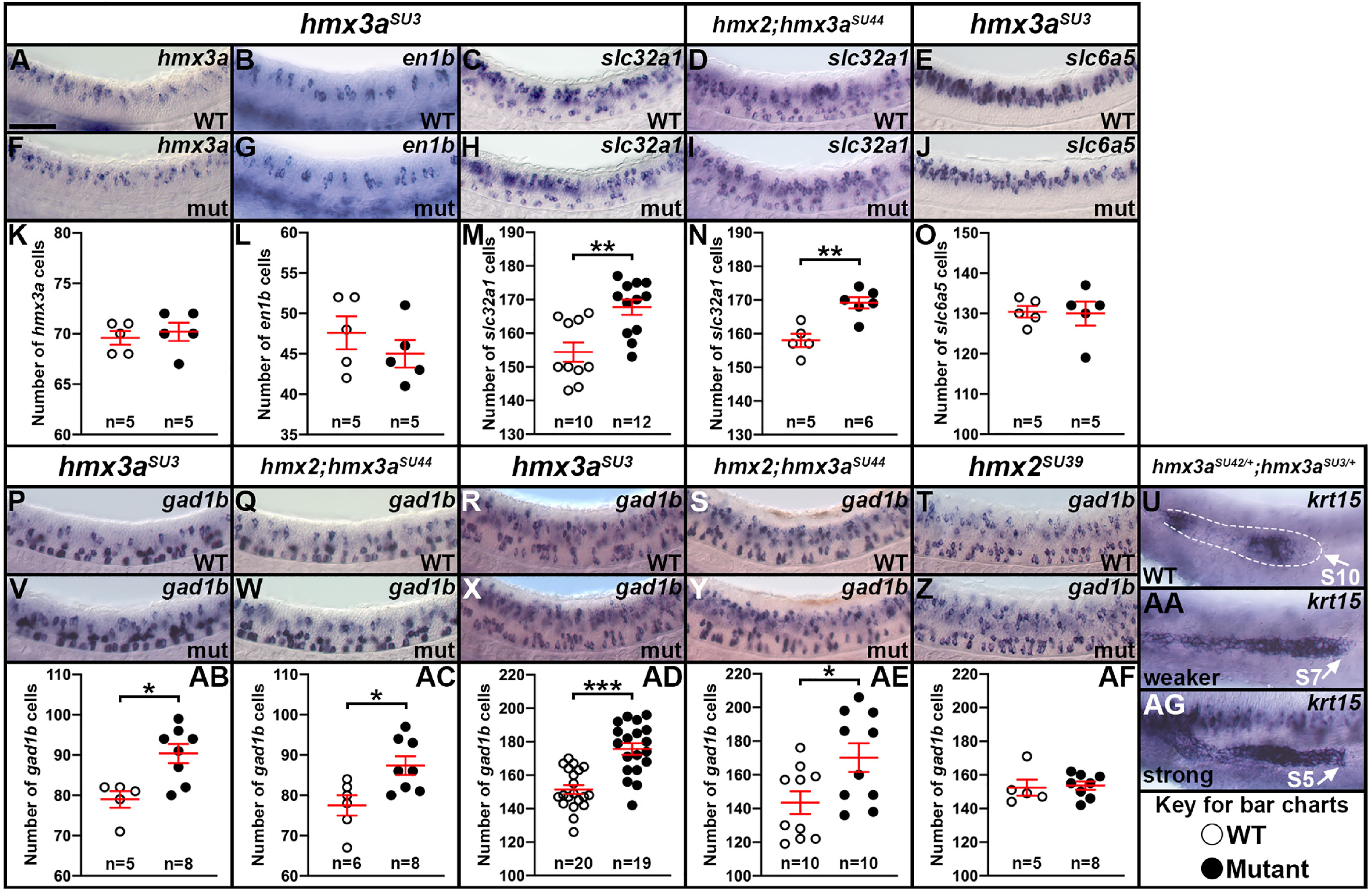
<u>Analysis of *hmx3a* single and *hmx2;hmx3a* deletion mutants.</u> (**A-J, P-AA & AG**) Lateral views of *hmx3a* (**A, F**), *en1b* (**B, G**) *slc32a1* (**C, D, H, I**) *slc6a5* (**E, J**), *gad1b* (**P-T, V-Z**) or *krt15* (**U, AA, AG**) expression in spinal cord (**A-J, P-T, V-Z**) or lateral line primordium (**U, AA, AG**) at 27 h (**A-J, P, Q, U-W, AA, AG**) or 48 h (**R-T, X-Z**). Rostral, left; Dorsal, up. (**K-O, AB-AF**). Number of cells expressing *hmx3a* (**K**), *en1b* (**L**), *slc32a1* (**M & N**), *slc6a5* (**O**) and *gad1b* (**AB-AF**) in a precisely-defined spinal cord region adjacent to somites 6-10 at 27 h (**K-O, AB & AC**) or 48 h (**AD-AF**). Data are depicted as individual value plots and the *n*-values for each genotype are also shown. For each plot, the wider red horizontal bar depicts the mean number of cells and the red vertical bars depict the standard error of the mean (standard error of the mean (S.E.M.) values are listed in Table 1B). All counts are an average of at least 5 embryos. Statistically significant (*p* < 0.05) comparisons are indicated with brackets and asterisks. *p* < 0.05 = *, *p* < 0.01 = **, *p* < 0.001 = ***. White circles indicate WT data and black circles the appropriate mutant data as indicated in key under panel AG. All data were first analyzed for normality using the Shapiro-Wilk test. Data sets in M, AB and AF are non-normally distributed and were analyzed with the Wilcoxon-Mann-Whitney test. Data sets in K, L, N, O, AC, AD, AE are normally distributed and so the F test for equal variances was performed. All of these had equal variances, so a type 2 student’s t-test was performed. P-values are provided in Table 1B. (**A, B, F, G, K, L**) As in DKD embryos (Fig. 3), dI2 and V1 interneurons do not die, nor do dI2 interneurons transfate/change into V1 interneurons in *hmx3a^SU3^* mutant embryos, since the numbers of *hmx3a-* (**A, F** & **K**) and *en1b*-expressing cells (**B, G** & **L**) do not change compared to WT embryos. There is a statistically significant increase in the number of inhibitory, *slc32a1*-expressing cells in *hmx3a^SU3^* mutants (**C, H, M**) and *hmx2;hmx3a^SU44^* mutants (**D, I, N**) compared to WT embryos. (**E, J, O**) However, at 27 h, the number of *slc6a5*-expressing cells is unchanged between WT and *hmx3a^SU3^* mutants, whereas there is an increase in the number of GABAergic (*gad1b*-positive) cells in *hmx3a^SU3^* (**P, V, AB**) and *hmx2;hmx3a^SU44^* mutants (**Q, W, AC**), suggesting that the additional inhibitory cells in the mutant embryos are GABAergic and not glycinergic. (**R, S, X, Y, AD, AE**) There is an equivalent increase in GABAergic (*gad1b*-positive) cells at 48 h in *hmx3a^SU3^* and *hmx2;hmx3a^SU44^* mutant embryos. However, there is no change in the number of GABAergic (*gad1b*-positive) cells at 48 h in *hmx2^SU39^* mutants, compared to WT embryos (**T, Z, AF**). (**U, AA, AG**). *hmx3a^SU42/+^;hmx3a^SU3/+^* trans-het embryos have two different lateral line primordium progression phenotypes at 27 h. Scale bar = 50 µm.

In contrast, embryos homozygous for *hmx3a^SU42^* do not have fused otoliths or stalled lateral line progression (Fig. 4, Fig. 5M, Table 2) and unlike *hmx3a^SU3^* mutants, they have normal expression of *hmx3a* in the anterior otic epithelium and adjacent anterior neuroblasts (Fig. 5S and cf. Fig. 5R & U’). *hmx3a^SU42^* mutants also have normal numbers of spinal cord glutamatergic neurons (Fig. 5B & G, Table 1B) and are homozygous viable (Fig. 4, Table 2). Interestingly, embryos homozygous for *hmx3a^sa23054^* have variable, incompletely penetrant otolith fusion phenotypes that range from no fusion, through incomplete fusion (Fig. 5T), to complete fusion, despite the fact that any protein encoded by this allele should retain more WT sequence than that encoded by *hmx3a^SU42^* (Fig. 4). However, *hmx3a^sa23054^* mutants have normal lateral line progression, no reduction in the number of spinal cord glutamatergic neurons and they are also viable (Fig. 4, Fig. 5C, H & N, Tables 1B & 2).

Surprisingly, we also found that all four of the different *hmx2* alleles we created in these experiments (*hmx2^SU35^*, *hmx2^SU36^*, *hmx2^SU37^*, *hmx2^SU38^*; Fig. 4) are homozygous viable and have no obvious defects in otolith development. We also examined lateral line progression and the number of spinal cord glutamatergic neurons in *hmx2^SU37^* and *hmx2^SU38^* homozygous mutants and did not detect any change compared to WT embryos (Fig. 4, Tables 1B & 2 and data not shown).

To test whether our *hmx2* mutants had no obvious phenotypes because they had retained some Hmx2 function, we created a large deletion allele, *hmx2^SU39^*, that deletes most of the *hmx2* genomic sequence. Only 84 nucleotides of 5’ and 60 nucleotides of the most 3’ coding sequence remain (Fig. 4). However, *hmx2^SU39^* homozygous mutants also lack fused otoliths and are homozygous viable. They also have normal expression of *hmx3a* and *pax5* in the anterior otic epithelium, normal expression of *hmx3a* in the adjacent anterior neuroblasts, the normal complement of three distinct cristae and two distinct maculae in the ear, and normal lateral line progression (Fig. 4, Fig. 5Q, W, AA, AE & AI, Tables 1B & 2). In addition, there is no change in the number of glutamatergic cells in the spinal cords of these mutants, when compared to WT sibling embryos (Fig. 5F & K, Table 1B).

While this was surprising, we hypothesized that the lack of obvious phenotypes could be because Hmx3a protein was compensating for loss of Hmx2 protein. To test this hypothesis, we created double mutant embryos. As *hmx2* and *hmx3a* are adjacent on chromosome 17, it is not possible to create double mutants by breeding single mutants. Both mutations have to exist on the same chromosome. Therefore, we created two different deletion alleles, *hmx2;hmx3a^SU44^* and *hmx2;hmx3a^SU45^*, that lack the entire *hmx3a* coding sequence and all but the last 66 nucleotides of *hmx2* coding sequence (Fig. 4 and materials and methods; the alleles differ only in the amount of remaining sequence upstream of the *hmx3a* locus). However, in embryos homozygous for these large deletion alleles, the reduction in the number of excitatory spinal cord interneurons and the increase in the number of inhibitory spinal neurons is equivalent to that in *hmx3a^SU3^* and *hmx3a^SU43^* single mutants (Fig. 5E & J, Fig. 6D, I & N and Table 1B). There are also no obvious differences in the otolith fusion or lateral line progression phenotypes between these single mutants and embryos homozygous for the double deletion alleles (Fig. 5P & V, compare to Fig. 5O and U). To further interrogate whether there might be subtle differences in ear phenotypes between *hmx3a^SU3^* and double deletion mutants, we examined the expression of *pax5* in the anterior otic epithelium and the presence and integrity of the cristae and maculae of the ear using phalloidin staining. Both the reduction in *pax5* expression in the anterior otic epithelium (cf. Fig. 5Z & Y), the fusion/juxtaposition of the maculae within a more ventro-medial position in the ear (cf. Fig. 5AH & AG) and the size and number of cristae in the ear (Fig. 5AB-AD) are equivalent in *hmx2;hmx3a^SU44^* double deletion and *hmx3a^SU3^* single mutants.

To determine whether the increase in the number of spinal inhibitory interneurons reflects an increase in glycinergic or GABAergic neurons, we examined the expression of genes expressed exclusively by cells with these inhibitory neurotransmitter phenotypes (*slc6a5* for glycinergic and *gad1b* for GABAergic, see materials and methods). While we found no statistically significant difference in the number of spinal cord glycinergic cells in *hmx3a^SU3^* mutants or *hmx2;hmx3a^SU44^* deletion mutants, there was a statistically significant increase in the number of GABAergic cells in *hmx3a^SU3^* and *hmx2;hmx3a^SU44^* mutants compared to WT sibling embryos (Fig. 6E, J, O, P, Q, V, W, AB, AC, Table 1B).

As *hmx2* spinal expression is initially weaker than *hmx3a* expression (Fig.1), we also analyzed *hmx2^SU39^* single mutants at 48 h, to determine if there was a spinal cord neurotransmitter phenotype at this later stage of development. We examined the number of GABAergic cells, as these are easier than glutamatergic cells to count at this stage. However, there was no change in the number of GABAergic spinal cells in *hmx2^SU39^* single mutants compared to WT embryos (Fig. 6 T, Z, AF, Table 1B). We also tested whether the spinal phenotype of embryos homozygous for the double deletion alleles was more severe than *hmx3a^SU3^* single mutants at 48 h. However, while the number of GABAergic cells was increased in both *hmx3a^SU3^* single mutants and *hmx2;hmx3a^SU44^* double deletion mutants, there was no statistically significant difference between these two phenotypes (Fig. 6R, S, X, Y, AD, AE, Table 1B).

### Trans-heterozygous crosses suggest that *hmx3a^SU42^* is a hypomorphic allele

*hmx3a^SU3^*, *hmx3a^SU42^* and *hmx3a^SU43^* mutant alleles all introduce a frameshift within 4 nucleotides of each other (nucleotides 319, 320, and 323 of the coding sequence respectively). Assuming that all three of these alleles are translated into truncated proteins, *hmx3a^SU3^* would retain 106 WT amino acids whereas the other two alleles would retain 107 WT amino acids (Fig. 4). However, despite the similarity of these mutant alleles, embryos homozygous for *hmx3a^SU3^* and *hmx3a^SU43^* have fused otoliths, stalled lateral line progression, altered spinal interneuron neurotransmitter phenotypes and do not survive to adulthood whereas embryos homozygous for *hmx3a^SU42^* are viable and lack all of these phenotypes. In addition, *hmx3a^sa23054^* mutants have variable otolith fusion phenotypes, despite retaining more WT sequence than *hmx3a^SU42^* alleles, and embryos homozygous for *hmx2^SU39^,* which almost completely deletes the *hmx2* coding sequence, have no obviously abnormal phenotypes. Given the surprising nature of these results, we decided to further investigate these alleles by creating trans-heterozygous animals. To do this we performed different pair-wise crosses between fish heterozygous for *hmx2^SU39^, hmx3a^SU3^*, *hmx3a^SU42^*, *hmx3a^SU43^*, *hmx3a^sa23054^* and *hmx2;hmx3a^SU44^* and analyzed ear and lateral line development in the resulting embryos.

Interestingly, when we crossed fish heterozygous for *hmx2;hmx3a^SU44^* with fish heterozygous for *hmx2^SU39^* the resulting embryos had normal otolith development and lateral line progression (Table 3). Given that approximately a quarter of these embryos should lack almost all of the coding sequence for both alleles of *hmx2* (*hmx2;hmx3a^SU44^* lacks all but the last 66 bp and *hmx2^SU39^* lacks all but the first 84 and last 60 bp of *hmx2a* coding sequence, see Fig. 4) as well as all of the coding sequence for one allele of *hmx3a*, this demonstrates that one WT allele of *hmx3a* is sufficient for normal otolith development and lateral line progression.

**Table 3:** Trans-heterozygous mutant analyses identify an allelic series of *hmx3a* mutants. Parents heterozygous for different mutant alleles were mated and otolith and lateral line progression phenotypes were assayed in their progeny. Otoliths were assayed at 3 d by visual inspection down a stereomicroscope. Lateral line progression was assayed by *in situ* hybridization for *krt15* at 27 h. Column one indicates the allele combination tested. Column two shows the percentage of embryos with otolith phenotypes, column three indicates the total number of embryos analyzed. In contrast to incrosses of *hmx3a^SU3^* or *hmx3a^SU43^,* where mutant embryos have fused otoliths in both ears (“severe” phenotype), in some trans-heterozygous crosses we observed two additional types of ear phenotypes. Phenotypes were classified as “moderate” if otoliths were adjacent but not fused in both ears, and as “weak” if there was an otolith fusion or adjacent otolith phenotype in only one ear. The percentage of embryos with each of these phenotypes is provided in columns five (severe), six (moderate) and seven (weak). Column eight indicates the percentage of embryos with stalled lateral line progression and column nine shows the total number of embryos analyzed. In contrast to incrosses of *hmx3a^SU3^* or *hmx3a^SU43^*, where mutant embryos lack any migration of the lateral line primordium along the trunk (“strong” phenotype), in some trans-heterozygous crosses, we observed embryos where the lateral line primordium had migrated slightly more caudally (“weaker” phenotype). Column 11 shows the percentage of embryos with the strong phenotype, column 12 shows the percentage of embryos with the slightly weaker phenotype. Chi-squared tests were performed to test if the frequency of embryos with otolith fusion or lateral line phenotypes was Mendelian and the P values for these tests are provided in columns 4 or 10 respectively. Statistically significant values are indicated in bold. We also performed a binomial distribution test, using the cumulative distribution function, to test whether the number of embryos from the *SU39 x SU44* cross that had fused otoliths was statistically significantly different from zero. P=0.264 for this test.

| Allele combination | % Ear phenotypes | # Embryos analyzed | P value | % Severe phenotype | % Moderate phenotype | % Weak phenotype | % Stalled lateral line progression | # Embryos analyzed | P value | % Strong phenotype | % Weaker phenotype |
| --- | --- | --- | --- | --- | --- | --- | --- | --- | --- | --- | --- |
| *SU39 x SU44* | 0.7% | 268 | <0.001 | 0.0% | 0.0% | 0.7% | 0.0% | 38 | <0.001 | 0.0% | 0.0% |
| *SU3 x SU43* | 22.6% | 623 | 0.165 | 22.5% | 0.0% | 0.1% | 32.4% | 37 | 0.250 | 32.4% | 0.0% |
| *SU3 x SU44* | 26.8% | 276 | 0.487 | 26.8% | 0.0% | 0.0% | 19.4% | 36 | 0.441 | 19.4% | 0.0% |
| *SU43 x SU44* | 28.3% | 279 | 0.214 | 28.0% | 0.0% | 0.3% | 29.8% | 114 | 0.192 | 26.3% | 3.5% |
| *sa23054 x SU44* | 22.4% | 303 | 0.289 | 22.4% | 0.0% | 0.0% | 26.3% | 38 | 0.703 | 18.4% | 7.9% |
| *sa23054 x SU3* | 26.6% | 139 | 0.696 | 26.6% | 0.0% | 0.0% | 17.6% | 34 | 0.244 | 0.0% | 17.6% |
| *SU42 x SU44* | 22.1% | 208 | 0.337 | 19.7% | 1.4% | 1.0% | 18.9% | 37 | 0.703 | 0.0% | 18.9% |
| *SU3 x SU42* | 17.1% | 1794 | <0.001 | 4.7% | 6.0% | 6.4% | 23.3% | 192 | 0.614 | 2.1% | 21.2% |

As expected, as both alleles produce homozygous mutant phenotypes, when we crossed fish heterozygous for *hmx3a^SU3^* with fish heterozygous for *hmx3a^SU43^* we observed mendelian ratios of embryos with fused otoliths and stalled lateral line progression (Table 3). Similarly, we obtained mendelian ratios of embryos with fused otoliths and stalled lateral line progression when we crossed fish heterozygous for *hmx3a^SU3^* with fish heterozygous for *hmx2;hmx3a^SU44^*, or fish heterozygous for *hmx3a^SU43^* with fish heterozygous for *hmx2;hmx3a^SU44^*.

More interestingly, when we crossed fish heterozygous for *hmx3a^sa23054^* with fish heterozygous for either *hmx2;hmx3a^SU44^* or *hmx3a^SU3^,* we also obtained mendelian ratios of embryos with fully penetrant otolith fusion phenotypes and stalled lateral line progression, as opposed to the variable otolith fusion phenotypes and normal lateral line progression that occurs in *hmx3a^sa23054^* homozygous mutants (cf. Table 2 & Table 3). Some of the embryos with stalled lateral line progression had the strong phenotype that we observe in *hmx3a^SU3^* and *hmx2;hmx3a^SU44^* homozygous mutants, and the rest had a slightly weaker phenotype (Table 3, similar to Fig. 6AA). This suggests that while two *hmx3a^sa23054^* alleles provide sufficient Hmx3a activity for normal lateral line progression and, in some cases, normal otolith development, this is not the case for either the combination of one *hmx3a^sa23054^* and one *hmx3a^SU3^* allele or the combination of one *hmx3a^sa23054^* allele over a *hmx3a* deletion.

Even more surprisingly, when we crossed fish heterozygous for *hmx3a^SU42^* with fish heterozygous for either *hmx2;hmx3a^SU44^* or *hmx3a^SU3^* we also obtained embryos with fused otoliths and stalled lateral line progression, although most of the embryos had the slightly weaker lateral line phenotype mentioned above (Fig. 6 AA & AG, Table 3). For the combination of *hmx3a^SU42^* and *hmx2;hmx3a^SU44^,* these phenotypes occurred in mendelian ratios. However, for the combination of *hmx3a^SU3^* and *hmx3a^SU42^,* while we observed a mendelian ratio of embryos with stalled lateral line progression, only 17% of embryos had abnormal otolith phenotypes and in most of these cases the otoliths were either adjacent but not fused in both ears or there was an abnormal otolith phenotype in only one ear (Table 3). When we genotyped a subset of these embryos, we found that all of the embryos with abnormal otolith phenotypes (n=18, 6 embryos each with either fused otoliths in both ears, adjacent otoliths in both ears, or a fused or adjacent otolith phenotype in only one ear) were *hmx3a^SU3/+^;hmx3a^SU42/+^* trans-hets. Interestingly, when we genotyped embryos with WT otolith phenotypes (two normal otoliths per ear, n=173), 5.20% were actually *hmx3a^SU3/+^;hmx3a^SU42/+^* trans-hets, 29.48% were only heterozygous for *hmx3a^SU3^*, 31.79% were only heterozygous for *hmx3a^SU42^* and 33.53% were homozygous wild type. Taken together, these data suggest that even though *hmx3a^SU42^* homozygous mutants have no obvious abnormal phenotypes, *hmx3a^SU42^* is a hypomorphic allele: while two alleles of *hmx3a^SU42^* provide sufficient Hmx3a activity for normal otolith development and lateral line progression, one *hmx3a^SU42^* allele combined with either one *hmx2;hmx3a^SU44^* or one *hmx3a^SU3^* allele does not.

### Loss of *hmx2* function can enhance hypomorphic *hmx3a* phenotypes

Our comparisons of *hmx3a^SU3^*, *hmx3a^SU43^*, *hmx2;hmx3a^SU44^* and *hmx2;hmx3a^SU45^* mutant phenotypes (Figs 5 & 6 and Table 1B) suggested that *hmx2* does not act redundantly with *hmx3a* in zebrafish otolith development, lateral line progression or specification of correct spinal interneuron neurotransmitter phenotypes, as complete removal of both genes does not result in more severe phenotypes than loss of just *hmx3a* function. However, if loss of *hmx3a* function already produces maximal mutant phenotypes, we wouldn’t detect stronger phenotypes in embryos homozygous for the double deletion alleles. Therefore, a more sensitive way to test if *hmx2* functions in these developmental processes, would be to remove *hmx2* function in embryos homozygous for a “weaker” hypomorphic *hmx3a* mutant allele, such as *hmx3a^SU42^*. Unfortunately, we cannot mate fish with the *hmx2^SU39^* and *hmx3a^SU42^* mutant alleles to make double mutants, as *hmx2* and *hmx3a* are adjacent on the same chromosome, so each mutant allele is tightly linked to a WT allele for the other gene. Therefore, we decided to knock-down Hmx2 function in *hmx3a^SU43^* mutants using CRISPR-mediated mutagenesis.

We injected CRISPR reagents to mutate *hmx2* into embryos from an incross of fish that were heterozygous for *hmx3a^SU42^*. We used the Microhomology-mediated End joining kNockout Target Heuristic Utility (MENTHU) tool to identify a sgRNA target site that should predominantly result in the same 5 bp deletion frame-shift allele (Fig. 4, Fig. 7C), being generated through microhomology-mediated end joining (MMEJ; Ata *et al*. 2018; Mann *et al*. 2019). We also used a two-part crRNA + tracrRNA system + Cas9 protein ribonucleoprotein complex for the injections, as this can produce a high efficiency of biallelic mutations and F0 phenotypes (DiNapoli *et al*. 2019; Lewis Lab unpublished data; Hoshijima *et al*. 2019). When we did this, we found that at ∼3.5 d, 28.25% (n=807) of *hmx2* CRISPR-injected embryos had an abnormal otolith phenotype (21.31% had fused otoliths in both ears and 6.94% had fused or adjacent otoliths in only one ear; Fig.7B, cf. to uninjected control, Fig.7A). In comparison, only 0.6% (n=670) of uninjected embryos and 2% (n=347) of embryos injected with a CRISPR crRNA ribonucleoprotein complex that we have used successfully to make mutations in an unrelated gene, had abnormal otolith phenotypes. These control experiments were performed at the same time as the *hmx2* CRISPR injections, using embryos obtained from the same heterozygous *hmx3a^SU42^* parent fish.

**Figure 7.**
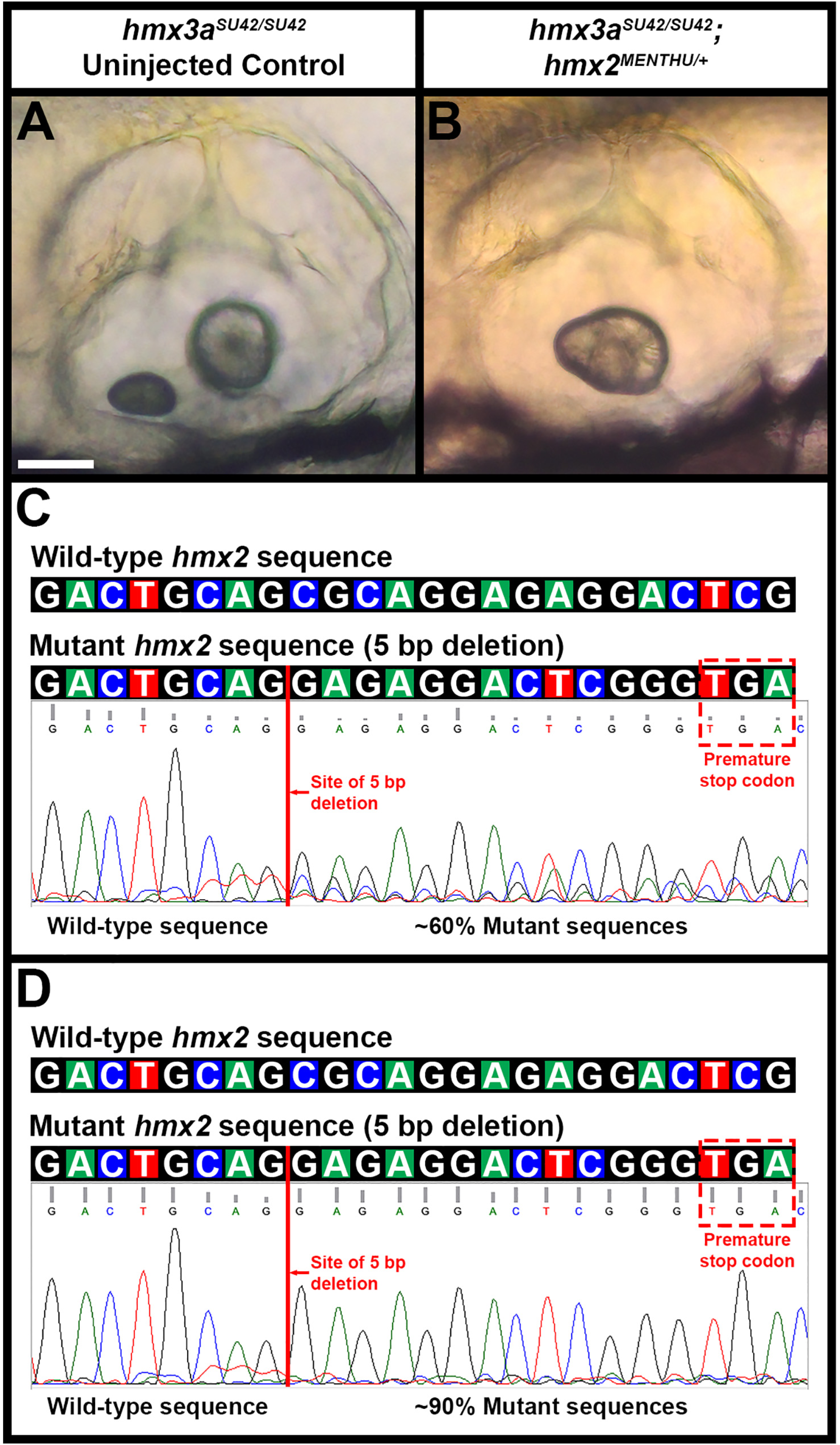
<u>Phenotypic and genotypic analysis of embryos from an incross of *hmx3a*</u>*<u>^SU42/+^</u>* <u>parents injected with *hmx2*</u>*<u>^MENTHU^</u>* <u>CRISPR reagents.</u> (**A-B**) Lateral views of ear phenotypes at 4 d in live uninjected (**A**) and *hmx2^MENTHU^* CRISPR-injected (B) *hmx3a^SU42^* homozygous mutant embryos. Rostral, left; Dorsal, top. The uninjected *hmx3a^SU42^* homozygous mutant embryo has two normal otoliths in each ear (**A**). In contrast, the *hmx2^MENTHU^* CRISPR-injected *hmx3a^SU42^* homozygous mutant embryo has fused otoliths in both ears. (**C-D**) Wild type (top row) and *hmx2^MENTHU^* (bottom row) genomic sequences on top. Each colored box represents a specific nucleotide in the *hmx2* coding sequence: A = green, C = blue, G = black, T = red. The *hmx2^MENTHU^* mutant sequence contains a 5 bp deletion (CGCAG (red line)) which introduces a premature stop codon (red dashed box) 14-16 bases after the deletion. Sequencing traces (below) from individual embryos from an incross of *hmx3a^SU42/+^* parents injected with *hmx2^MENTHU^* CRISPR reagents. In the injected embryos, *hmx2^MENTHU^* mutant sequences (with the 5 bp deletion) comprise approximately 60% (C) to 90% (**D**) of all amplified sequences at this locus. Scale bar = 50 µm (A& B).

We examined 40 of the *hmx2* CRISPR-injected embryos at ∼30 h for lateral line progression phenotypes and then let these embryos develop to ∼3.5 d, so we could correlate lateral line and otolith phenotypes. 25% of the embryos had strong or medium stalled lateral line progression phenotypes and all of these embryos also developed fused otoliths in both ears (Table 4A). A few additional embryos had a weaker lateral line progression defect (migration of the primordium was only delayed by 2-3 somites compared to stage-matched injected siblings) and two of these also developed fused otoliths in both ears. When we genotyped these embryos for *hmx3a^SU42^*, we found that all of the embryos with fused otoliths were homozygous for *hmx3a^SU42^* (Table 4B).

**Table 4.** Lateral line primordium and otolith phenotypes in embryos from incrosses of hmx3a^SU42/+^ parents, injected with hmx2^MENTHU^ CRISPR reagents. Table 4A. Embryos from an incross of *hmx3a^SU42/+^* parents were injected with *hmx2^MENTHU^* CRISPR reagents at the one-cell stage and assayed for lateral line primordium and otolith phenotypes at 30 h and 72 h respectively. Rows 2-5 show stalled lateral line primordium migration phenotypes: normal = primordium in expected position (over somite 15 at 30h); weak = primordium migration stalled by 2-3 somites; medium = primordium moderately stalled (over somite 6-7 at 30 h); strong = primordium not detected. Columns 2-3 show otolith phenotypes. Column 4 = total number of embryos with each lateral line phenotype. Row 6 = total number of embryos with each otolith phenotype. Table 4B. *hmx3a^SU42^* genotypes of the 40 injected embryos screened for lateral line primordium and otolith phenotypes in Table 4A. Rows 2-7 show combinations of lateral line primordium phenotype (normal, weak, medium, strong, as in Table 4A) and otolith phenotype (fused in both ears, two normal otoliths in both ears, as in Table 4A). Columns 2-4 show *hmx3a^SU42^* genotypes. Column 5 = total number of embryos with each combination of lateral line primordium and otolith fusion phenotypes. Row 7 = total number of embryos with each *hmx3a^SU42^* genotype. Table 4C. *hmx3a^SU42^* genotypes of embryos from a second incross of *hmx3a^SU42/+^* parents injected with *hmx2^MENTHU^* CRISPR reagents at the one-cell stage and assayed for otolith fusion phenotypes at 72 h. Rows 2-4 show otolith phenotypes. Columns 2-4 show *hmx3a^SU42^* genotypes. Column 5 = total number of embryos with each otolith phenotype. Row 5 = total number of embryos with each *hmx3a^SU42^* genotype. Table 4D. A subset of 23 embryos screened in Table 4C were also sequenced to assess whether they contained any *hmx2^MENTHU^* mutant sequences. Rows 2-4 indicate otolith phenotypes. Columns 2-4 indicate *hmx3a^SU42^* genotypes. Column 5 = approximate percentage of *hmx2^MENTHU^* mutant sequences detected in the PCR amplicons for embryos with each otolith phenotype. In all cases, the *hmx2^MENTHU^* mutant sequences represented at least 60% of all *hmx2* sequences in each PCR amplicon.

| **Table 4A. Lateral line and otolith phenotypes of 40 embryos from *hmx3a*<sup>SU42/+</sup> parents injected with *hmx2* CRISPR reagents** |  |  |  |  |
| --- | --- | --- | --- | --- |
| Lateral line phenotype | Fused otoliths both ears | Normal otoliths both ears | Total |  |
| Strong | 6 | 0 | 6 |  |
| Medium | 4 | 0 | 4 |  |
| Weak | 2 | 5 | 7 |  |
| Normal | 1 | 22 | 23 |  |
| Total | 13 | 27 | 40 |  |
| **Table 4B. *hmx3a*<sup>SU42</sup> genotypes of the 40 embryos included in Table 4A** |  |  |  |  |
| Lateral line /Otolith phenotype | *hmx3a*<sup>SU42</sup> homozygous | *hmx3a*<sup>SU42</sup> heterozygous | *hmx3a*<sup>SU42</sup> WT | Total |
| Strong/Fused | 6 | 0 | 0 | 6 |
| Medium/Fused | 4 | 0 | 0 | 4 |
| Weak/Fused | 2 | 0 | 0 | 2 |
| Normal/Fused | 1 | 0 | 0 | 1 |
| Weak/Normal | 0 | 4 | 1 | 5 |
| Normal/Normal | 0 | 16 | 6 | 22 |
| Total | 13 | 20 | 7 | 40 |
| **Table 4C. Genotyping embryos with abnormal and normal otolith phenotypes** |  |  |  |  |
| Otolith Phenotype | *hmx3a*<sup>SU42</sup> homozygous | *hmx3a*<sup>SU42</sup> heterozygous | *hmx3a*<sup>SU42</sup> WT | Total |
| Fused both ears | 32 | 1 | 0 | 33 |
| Fused or adjacent one ear | 3 | 1 | 0 | 4 |
| Normal both ears | 1 | 26 | 8 | 35 |
| Total | 36 | 28 | 8 | 72 |
| **Table 4D. *hmx3a*<sup>SU42</sup> genotypes and otolith phenotypes of embryos genotyped for *hmx2*** |  |  |  |  |
| Otolith phenotype | *hmx3a*<sup>SU42</sup> homozygous | *hmx3a*<sup>SU42</sup> heterozygous | *hmx3a*<sup>SU42</sup> WT | % of *hmx2* mutant sequences |
| Fused both ears | 9 | 1 | 0 | 60-90% |
| Fused one ear | 1 | 1 | 0 | 60-75% |
| Normal both ears | 0 | 4 | 7 | 60-90% |

We also genotyped 72 additional *hmx2* CRISPR-injected embryos, just over half of which had otolith defects. The vast majority of the embryos with otolith phenotypes were homozygous for *hmx3a^SU42^* (Fig. 7B, Table 4C, one embryo with fused otoliths in both ears and one embryo with an otolith defect in one ear only were heterozygous). In contrast, all except one of the 35 embryos that did not have obvious defects in otolith development were heterozygous for *hmx3a^SU42^* or WT (Table 4C).

Taken together, these results suggest that we obtained a high efficiency of *hmx2* mutations in our injected embryos and that CRISPR-mediated knock-down of Hmx2 causes *hmx3a^SU42^* mutants to have defects in otolith development and lateral line progression. To test this, we sequenced the *hmx2* allele from 23 of the embryos that we had genotyped for *hmx3a^SU42^* that had different otolith phenotypes (Table 4D). We found that all of these embryos had a substantial frequency of *hmx2* nonsense alleles. As predicted by the MENTHU algorithm, the mutated alleles all contained a 5 bp deletion, although some also had additional mismatches in the three bases immediately prior to the deletion and the location of the deletion differed by 1 bp in a few cases. In all cases, we estimate that at least 60% of the amplified *hmx2* sequences were mutant (Table 4D; Fig. 7C, D). In one of the WT embryos that lacked a phenotype, approximately 90% of the amplified *hmx2* sequences were mutant, suggesting that, consistent with the lack of abnormal phenotypes in *hmx2^SU39^* mutants, CRISPR mutagenesis of *hmx2* is not sufficient for abnormal otolith development (Fig. 7C, D).

### *hmx2* and *hmx3a* are not expressed maternally

One possible explanation for why the spinal cord phenotype is less severe in *hmx2/3a* deletion mutants than in morpholino-injected DKD embryos would be if *hmx2* and/or *hmx3a* are maternally expressed, as in this case the morpholinos might knock-down both maternal and zygotic function whereas the mutants would only remove zygotic function. In addition, maternal expression of *hmx2* might explain the lack of any obvious abnormal phenotypes in *hmx2* single mutants. To test this, we performed *in situ* hybridization for *hmx2* and *hmx3a* at the 16-cell stage. However, we did not detect any maternal expression of *hmx2*, *hmx3a* or any of the other *hmx* genes (Fig. 8A-E). We also performed qRT-PCR for *hmx2* and *hmx3a* on whole embryos at different developmental stages. We did not observe expression of either gene at either the 16-cell stage or at 6 h, suggesting that neither *hmx2* nor *hmx3a* are maternally expressed (Fig. 8F). At 14 h, shortly after when both of these genes start to be expressed in the ear and spinal cord, we observed low levels of expression and, for both genes, as expected, this became more abundant at 27 h and 48 h (Fig. 8F). Finally, we also generated embryos from adults that were homozygous mutant for *hmx2^SU38^, hmx2^SU39^* and *hmx3a^sa23054^*. However, even though half of the embryos in each of these crosses should have been maternal zygotic mutants, we still did not observe any embryos with fused otoliths (Fig. 4, Table 2).

**Figure 8.**
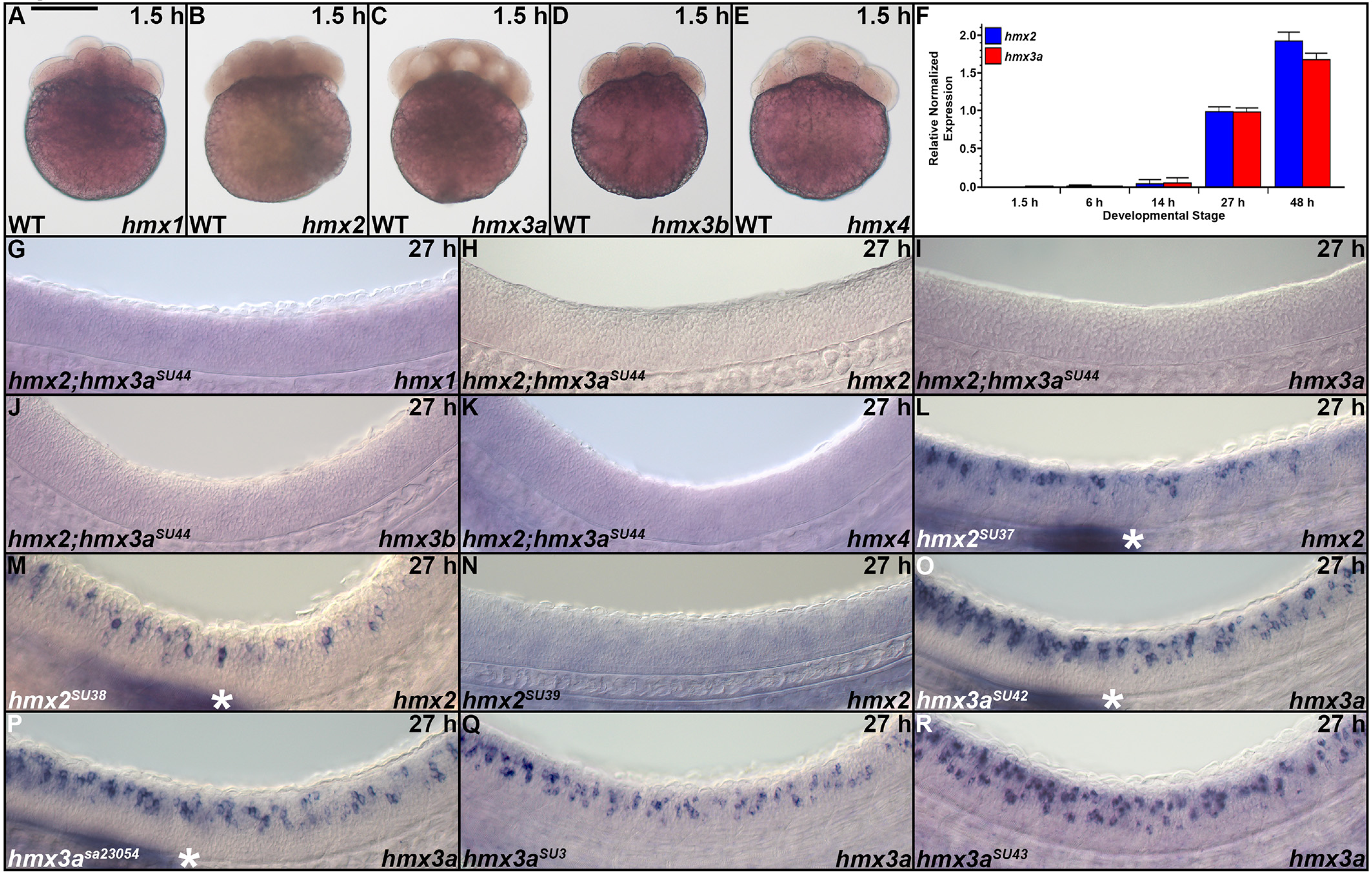
<u>Expression of *hmx* genes in mutant zebrafish embryos and before the mid-blastula transition.</u> (**A-E, G-R**) Lateral views of expression in whole embryos at 1.5 h (16-cells, **A-E**) or the spinal cord (**G-R**) at 27 h or. (**A-E)** Animal pole, up. (**G-R**) Rostral, left; Dorsal, up. (**L-M, O-P**) White asterisk = expression in the lateral line primordium. None of the *hmx* genes are maternally expressed at 1.5 h, as assessed by *in situ* hybridization (**A-E**), and, in the case of *hmx2* and *hmx3a*, qRT-PCR on whole embryos (**F**). No maternal expression of *hmx2* and *hmx3a* was detected and zygotic expression was not observed via qRT-PCR until 14 h (**F**). *hmx1* (**G**), *hmx3b* (**J**) and *hmx4* (**K**) are not expressed in the spinal cord of *hmx2;hmx3a^SU44^* deletion mutants. However, *hmx1* and *hmx4,* were still expressed in the head as shown in Fig. 1 (data not shown), confirming that the *in situ* hybridization experiment had worked. We never detect expression of *hmx3b* in WT embryos at 27 h (see Fig. 1). (**H-I**) As expected, given the deletion of the entire *hmx3a* coding sequence and all but the last 66 bp of *hmx2* coding sequence in *hmx2;hmx3a^SU44^* mutants (Fig, 4), we did not detect any *hmx2* (**H**) or *hmx3a* (**I**) transcripts in these mutants. (**L-M**) *hmx2* mRNA does not exhibit nonsense-mediated decay (NMD) in *hmx2^SU37^* or *hmx2^SU38^* mutants. (**N**) In *hmx2^SU39^* mutants, deletion of all but the first 84 and the last 60 bases of *hmx2* coding sequence (Fig. 4) generates a severely truncated *hmx2* transcript that cannot be detected by our *hmx2* ISH probe. Generation of a short ISH probe targeted to the predicted truncated transcript product of *hmx2^SU39^* mutants also failed to detect *hmx2* expression in these mutants (data not shown). (**O-R**) *hmx3a* mRNA does not exhibit NMD in *hmx3a^SU42^* (**O**), *hmx3a^sa23054^* (**P**), *hmx3a^SU3^* (**Q**) or *hmx3a^SU43^* (**R**) mutant embryos. Scale Bar = 280 µm (**A-E**), 50 µm (**G-R**).

### *hmx3b*, *hmx1* and *hmx4* expression is not upregulated in *hmx2;hmx3a^SU44^* deletion mutants

Even though *hmx3b*, *hmx1* and *hmx4* are not normally expressed in the spinal cord (Fig.1), it was theoretically possible that they are upregulated in response to the absence, or reduced levels, of either Hmx2 and/or Hmx3a protein function, in which case they could partially substitute for the loss of *hmx2* and/or *hmx3a*. To test this, we performed *in situ* hybridization for these genes in *hmx3a^SU3^* and *hmx2;hmx3a^SU44^* mutants at 27 h. In both cases, we did not observe any spinal cord expression of these genes in either genotyped mutants or their sibling embryos, although, as observed previously in WT embryos (Fig. 1), *hmx1* and *hmx4* were expressed in the eye, ear and anterior lateral line neuromasts in both mutants and WT sibling embryos (Fig. 8G, J, K and data not shown). As expected, given the deletion of the entire *hmx3a* coding sequence and all but the last 66 bp of *hmx2* coding sequence in *hmx2;hmx3a^SU44^* mutants (Fig, 4), we did not detect any *hmx2* or *hmx3a* transcripts in these mutants (Fig. 8H-I).

### *hmx2^SU39^* mutants do not lack a spinal cord phenotype because of genetic compensation

Recent reports have demonstrated that genetic compensation (upregulation of other genes that can compensate for loss of the mutated gene) can result in loss-of-function mutants having a less severe phenotype than embryos injected with a morpholino against the same gene (Rossi *et al*. 2015; El-Brolosy and Stainier 2017; Zhu *et al*. 2017; Sztal *et al*. 2018; El-Brolosy *et al*. 2019; Peng 2019). If this is the case, then morpholino knock-down should have less effect on mutant embryos than on WT sibling embryos, because the morpholino will not affect upregulated compensating genes (e.g. see Rossi *et al*. 2015; Sztal *et al*. 2018). Therefore, to test whether the lack of a spinal cord phenotype in *hmx2^SU39^* deletion mutants is due to genetic compensation, we injected the translation-blocking *hmx2* morpholino into embryos from a cross of fish heterozygous for *hmx2^SU39^* and performed *in situ* hybridization for *slc17a6a/b* to label glutamatergic spinal cord interneurons. We predicted that if there was genetic compensation in *hmx2^SU39^* mutants, morpholino-injected WT sibling embryos should have a reduced number of spinal cord glutamatergic interneurons, whereas morpholino-injected *hmx2^SU39^* homozygous mutant embryos should have normal numbers of these cells. However, if there is no genetic compensation, we would expect a similar frequency of morpholino-injected homozygous mutant and WT embryos to have spinal cord phenotypes and those phenotypes to be roughly equivalent in severity. In contrast, if morpholino-injected homozygous mutant embryos have more severe phenotypes than morpholino-injected WT siblings, this might suggest that *hmx2^SU39^* is a hypomorphic allele. However, this seemed highly unlikely given that the *hmx2* gene is almost completely deleted in this allele (Fig. 4) and concordantly we do not detect any *hmx2* transcripts by *in situ* hybridization (Fig. 8N).

We initially examined the injected embryos down a stereomicroscope and divided them into two groups: those that had an obvious reduction in glutamatergic cells and others that either lacked a phenotype or had a more subtle phenotype. When we genotyped these embryos, in both groups we found homozygous mutant and WT embryos at frequencies that were not statistically significantly different from Mendelian ratios (Table 5). In addition, when we compared the average number of glutamatergic cells in morpholino-injected WT and mutant embryos, there was no statistically significant difference between them, <u>regardless</u> of whether we compared all of the morpholino-injected embryos of each genotype or just compared embryos within the same phenotypic group (Table 5). This suggests that the lack of an abnormal spinal cord phenotype in *hmx2^SU39^* homozygous mutant embryos is not due to genetic compensation and that the differences that we observed between the two groups of injected embryos instead probably reflect exposure to different levels of morpholino (see materials and methods).

**Table 5.** The lack of spinal cord phenotypes in *hmx2^SU39^* mutants is not due to genetic compensation. Spinal cord phenotypes were assessed by *slc17a6a/b in situ* hybridization. *hmx2* translation-blocking morpholino-injected embryos from an incross of *hmx2^SU39/+^* parents were visually inspected on a stereomicroscope and categorized as resembling either a “WT-like” or “morphant-like” spinal cord phenotype. Embryos were genotyped to identify homozygous WTs and mutants. The spinal cord phenotypes of these embryos were then analyzed on a compound microscope while blind to genotype. Values in columns 2 and 3 of lower table indicate the mean number of labelled cells in the spinal cord region adjacent to somites 6-10 ± S.E.M. P value in upper table is from Chi-squared test. P value in bottom table is either from a Wilcoxon-Mann-Whitney test (^^^, performed when data was not normally distributed) or from a type 2 Student’s t-test (^+^, performed when data was normally distributed and variances were equal) for the comparison of homozygous WT embryos to homozygous *hmx2^SU39^* mutants for a particular classification (values on same row). All P values are rounded up to 2 decimal places. See materials and methods for more information on statistical tests.

| **<u>Genotype</u>**<br>**Embryo class** | **WT** | **Het** | ***hmx2*<sup>SU39</sup>**<br>**mutants** | **N value** | **P value** |
| --- | --- | --- | --- | --- | --- |
| Uninjected control | 33.3% | 57.6% | 9.1% | 33 | 0.11 |
| WT-like | 44.4% | 22.2% | 33.3% | 9 | 0.12 |
| Morphant-like | 32.3% | 41.9% | 25.8% | 31 | 0.68 |
| **<u>Genotype</u>**<br>**Embryo class** | **WT** | ***hmx2*<sup>SU39</sup>**<br>**mutants** |  | **P value** |  |
| Uninjected control | 106.36 ± 1.75 | 109.67 ± 0.67 |  | 0.21<sup>^</sup> |  |
| WT-like | 104.25 ± 5.30 | 98.33 ± 4.06 |  | 0.86<sup>^</sup> |  |
| Morphant-like | 78.10 ± 2.50 | 71.25 ± 3.25 |  | 0.11<sup>+</sup> |  |
| All injected | 85.57 ± 3.96 | 78.64 ± 4.56 |  | 0.26<sup>+</sup> |  |

### *hmx3a^SU42^* mutant alleles do not lack a spinal cord phenotype because of genetic compensation

We also tested whether the spinal cord phenotype of *hmx3a^SU42^* mutants is less severe than embryos injected with a *hmx3a* morpholino because of genetic compensation. As above, we injected the translation-blocking *hmx3a* morpholino into embryos from a cross of fish heterozygous for *hmx3a^SU42^* and performed *in situ* hybridization for *slc17a6a/b*. When we examined the injected embryos down a stereomicroscope, we were able to separate them into one group that had an obvious reduction in spinal glutamatergic cells and another group that either lacked, or had a more subtle, phenotype. However, when we genotyped the embryos in these two groups, we found similar numbers of homozygous mutant and WT embryos in each group and the frequencies of different genotypes were not statistically significantly different from Mendelian ratios (Table 6). In addition, when we compared the average number of glutamatergic cells in morpholino-injected WT and mutant embryos, there was no statistically significant difference between them, <u>regardless</u> of whether we compared all of the morpholino-injected embryos of each genotype or just compared embryos within the same phenotypic group (Table 6). These data suggest that the lack of an abnormal spinal cord phenotype in *hmx3a^SU42^* homozygous mutant embryos is not due to genetic compensation and that the differences that we observed between the two groups of injected embryos just reflect exposure to different levels of morpholino (see materials and methods). Consistent with this, we also do not observe any nonsense-mediated decay (NMD) of *hmx3a* mRNA in *hmx3a^SU42^* mutants (Fig. 8O; NMD has been suggested to play a key role in at least some instances of genetic compensation; El-Brolosy *et al*. 2019).

**Table 6.** The lack of spinal cord phenotypes in *hmx3a^SU42^* mutants is not due to genetic compensation. Spinal cord phenotypes were assessed by *slc17a6a/b in situ* hybridization. *hmx3a* translation-blocking morpholino-injected embryos from an incross of *hmx3a^SU42/+^* parents were visually inspected on a stereomicroscope and categorized as resembling either a “WT-like” or “morphant-like” spinal cord phenotype. Embryos were genotyped to identify homozygous WTs and mutants. The number of glutamatergic cells in the spinal cord region adjacent to somites 6-10 was then counted in each embryo, using a compound microscope, while blind to genotype. Values in columns 2 and 3 of the lower table indicate the mean number of labelled cells ± S.E.M. The P value in the upper table is from a Chi-squared test to determine if the frequencies of different genotypes were Mendelian. The P value in the bottom table is either from a type 2 (^+^, performed when data was normally distributed and variances were equal), or type 3 Student’s t-test (^§^, performed when data was normally distributed and variances were unequal) for the comparison of homozygous WT embryos to homozygous *hmx3a^SU42^* mutants for a particular classification (values on same row). All P values are rounded up to 2 decimal places. See materials and methods for more information on statistical tests.

| **Genotype**<br>**Embryo class** | **WT** | **Het** | ***hmx3a*<sup>SU42</sup>**<br>**mutants** | **N value** | **P value** |
| --- | --- | --- | --- | --- | --- |
| Uninjected control | 22.5% | 52.5% | 25.0% | 40 | 0.93 |
| WT-like | 31.25% | 31.25% | 37.5% | 16 | 0.30 |
| Morphant-like | 20.5% | 56.4% | 23.1% | 39 | 0.61 |
| **Genotype**<br>**Embryo class** | **WT** |  | ***hmx3a*<sup>SU42</sup>**<br>**mutants** | **P value** |  |
| Uninjected control | 117.56 ± 2.26 |  | 116.10 ± 1.38 | 0.58<sup>+</sup> |  |
| WT-like | 116.80 ± 2.89 |  | 114.67 ± 2.69 | 0.60<sup>+</sup> |  |
| Morphant-like | 86.38 ± 2.15 |  | 88.22 ± 5.07 | 0.74<sup>§</sup> |  |
| All injected | 98.08 ± 4.58 |  | 98.80 ± 4.67 | 0.91<sup>+</sup> |  |

### *hmx3a^SU3^* mutants also do not have genetic compensation

We also tested whether *hmx3a^SU3^* mutants have a less severe spinal cord phenotype than morpholino knock-down embryos because of genetic compensation. As above, we injected the translation-blocking *hmx3a* morpholino into embryos from a cross of fish heterozygous for *hmx3a^SU3^* and performed *in situ* hybridization for *slc17a6a/b*. When we examined these embryos down a stereomicroscope, some of the embryos appeared to have a severe reduction in glutamatergic cells that resembled the morpholino knock-down phenotype, whereas in the other embryos any reduction was more subtle. However, when we genotyped these embryos, we again found similar numbers of homozygous mutant and WT embryos in both groups and the frequencies of the different genotypes in each group were not statistically significantly different from Mendelian ratios (Table 7). In addition, when we compared the average number of glutamatergic cells in morpholino-injected WT and mutant embryos, there was no statistically significant difference between them, <u>regardless</u> of whether we compared all of the morpholino-injected embryos, or just compared the injected embryos within a particular phenotypic group (Table 7). For the “weaker” phenotypic group, the difference between WT and mutant embryos approached statistical significance. However, this is probably because some of the WT embryos in this category had almost no reduction in the number of glutamatergic cells, whereas all of the mutants in this category had at least their normal mutant phenotypes. These results suggest that the differences that we observed between the two groups of injected-embryos probably just reflect exposure to different levels of morpholino (see materials and methods), and that embryos in the “weaker” phenotypic group did not receive sufficient morpholino to effectively knock-down *hmx3a* mRNA or cause the more severe morpholino phenotype. Taken together, these data suggest that the spinal cord phenotype in *hmx3a^SU3^* homozygous mutant embryos is not less severe than the morpholino knock-down phenotype because of genetic compensation. Consistent with this, we also do not detect any NMD of *hmx3a* mRNA in *hmx3a^SU3^* mutants (Fig. 8Q).

**Table 7.**
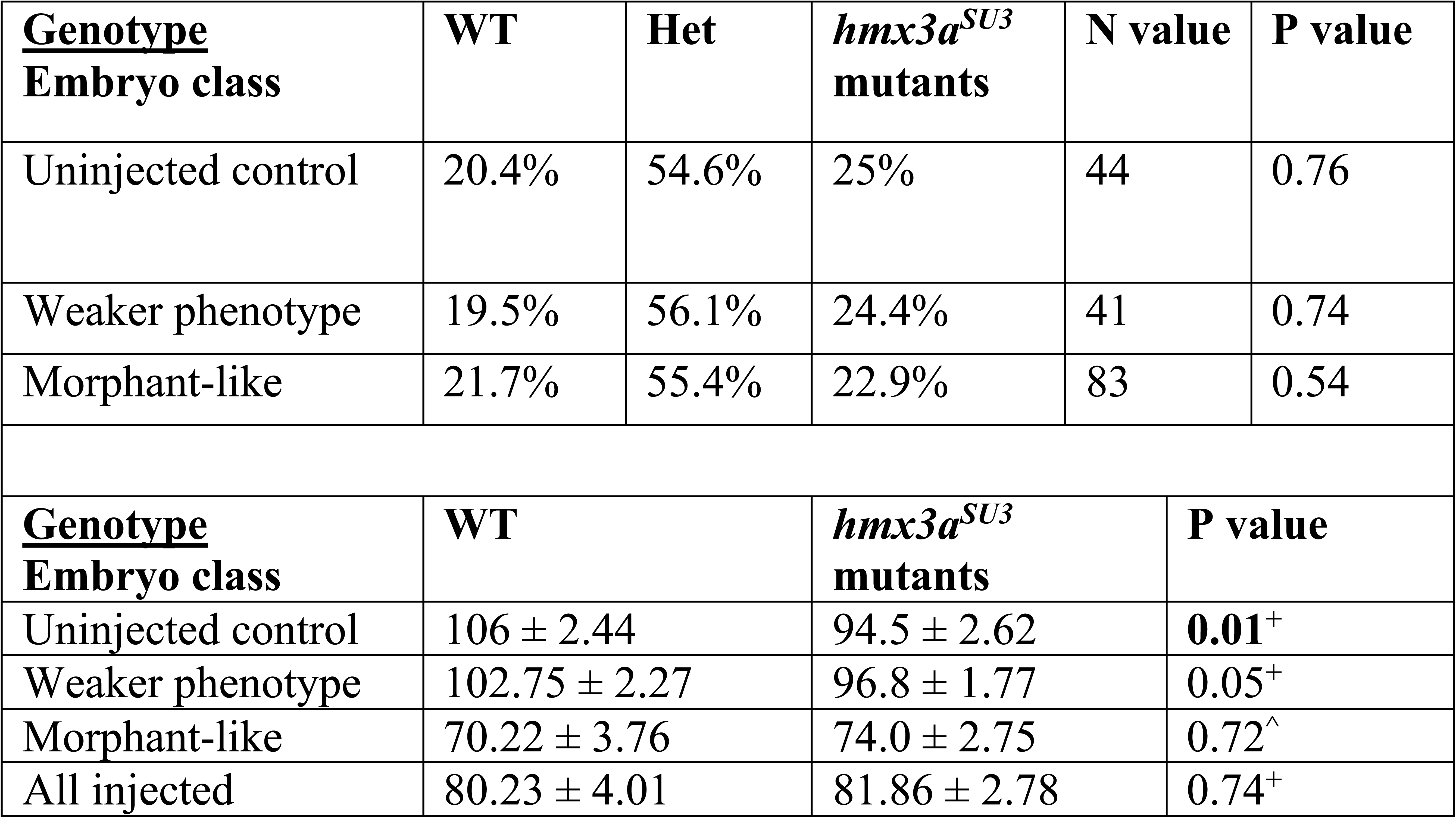
The incomplete penetrance of the spinal cord phenotype in *hmx3a^SU3^* mutants is not due to genetic compensation. Spinal cord phenotypes were assessed by *slc17a6a/b in situ* hybridization. *hmx3a* translation-blocking morpholino-injected embryos from an incross of *hmx3a^SU3/+^* parents were visually inspected on a stereomicroscope and categorized as resembling either a “weaker” or more severe, “morphant-like” spinal cord phenotype. Embryos were genotyped to identify homozygous WTs and mutants. The number of glutamatergic cells in the spinal cord region adjacent to somites 6-10 was then counted in each embryo, using a compound microscope, while blind to genotype. Values in columns 2 and 3 of the lower table indicate the mean number of labelled cells ± S.E.M. The P value in the upper table is from a Chi-squared test to determine if the frequencies of different genotypes were Mendelian. The P value in the bottom table is either from a Wilcoxon-Mann-Whitney test (^^^, performed when data was not normally distributed) or from a type 2 Student’s t-test (^+^, performed when data was normally distributed and variances were equal) for the comparison of homozygous WT embryos to homozygous *hmx3a^SU3^* mutants for a particular classification (values on same row). All P values are rounded up to 2 decimal places. See materials and methods for more information on statistical tests. Statistically significant values are indicated in bold.

## Discussion

### *hmx3a* is required for correct neurotransmitter phenotypes of a subset of spinal cord interneurons

In this paper, we identify for the first time, a requirement for *hmx3a* in spinal cord interneuron development. We demonstrate that *hmx2* and *hmx3a* are expressed by V1 and dI2 interneurons, which is consistent with very recent scRNA-Seq data from mouse spinal cord (Delile *et al*. 2019). Of these cell types, only dI2 interneurons are glutamatergic. Therefore, the most likely explanation for the reduction in the number of glutamatergic spinal cord cells in *hmx3a* mutants is that some dI2 interneurons are losing their glutamatergic phenotypes. Given that we also detect a corresponding increase in GABAergic spinal cord cells, but the number of V1 cells (indicated by *en1b* expression) does not change, it is likely that the dI2 interneurons that are losing their glutamatergic phenotypes are becoming GABAergic instead. Unfortunately, the respective *in situ* hybridization probes are not strong enough to formally confirm with double-labeling experiments that dI2 interneurons switch their neurotransmitter phenotype from glutamatergic to GABAergic. However, unless Hmx3a is acting in a cell non-autonomous manner, which we think is unlikely as this protein has a nuclear localization sequence and no obvious signal peptide, this is the most likely explanation of our data.

### *hmx3a* is required for progression of the posterior lateral line primordium

Feng and Xu previously reported that the number of posterior lateral line primordium neuromasts was either severely reduced or completely lost at 3 d in *hmx2/3a* DKD animals (Feng & Xu, 2010). Intriguingly, the few neuromasts that sometimes persisted were located very rostrally in the embryo, close to the earliest forming somites. Our analyses demonstrate that at 27 h, when the posterior lateral line primordium has migrated to somite 10 in WT embryos, in *hmx3a^SU3^, hmx3a^SU43^, hmx2;hmx3a^SU44^* and *hmx2;hmx3a^SU45^* mutants the primordium is stalled adjacent to somites 1-4. This suggests that the previously reported loss of neuromasts at 3 d is probably caused by the posterior lateral line primordium failing to migrate and deposit neuromasts. Feng & Xu also described reduced cell proliferation (at 15 h) and reduced *hmx3a* expression (at 24 h) in the posterior lateral line primordium of *hmx2/3a* DKD animals. Whilst we cannot rule out the possibility that the lateral line primordium fails to migrate because it has not formed correctly, we observe persistent expression of both *hmx3a* and *krt15* in the stalled primordium of our *hmx3a^SU3^, hmx3a^SU43^, hmx2;hmx3a^SU44^* and *hmx2;hmx3a^SU45^* mutants (data not shown), suggesting that some other mechanism, possibly chemosensory, might underlie the stalled migration.

### Hmx3a protein may not require its homeodomain for its functions in viability and otolith, lateral line and spinal cord interneuron development

Our results also suggest that Hmx3a protein may not require its homeodomain for either its role in viability or its essential functions in otolith development, lateral line progression and correct specification of a subset of spinal cord neurotransmitter phenotypes. This is surprising because most homeodomain proteins act as transcription factors and use their homeodomain to bind DNA and regulate gene expression. Instead, our data suggest that there may be at least one other, as yet undiscovered, crucial functional domain in the N-terminal region of Hmx3a, that is required for its functions in embryo development and viability, as embryos homozygous for *hmx3a^SU42^* are viable and have no obvious abnormal phenotypes and embryos homozygous for *hmx3a^sa23054^* are also viable, have normal lateral line progression and spinal cord interneuron neurotransmitter phenotypes and produce viable progeny. It is highly unlikely that the lack of obvious abnormal phenotypes in these two different mutants is due to an alternative translation start site creating a truncated Hmx3a protein that contains the homeodomain, as the only downstream methionine in *hmx3a* is more than a third of the way through the homeodomain, and also, in this case we would expect the *hmx3a^SU3^* and *hmx3a^SU43^* alleles to also make this truncated protein. The lack of obviously abnormal phenotypes in *hmx3a^SU42^* and *hmx3a^sa23054^* homozygous mutants also can not be explained by alternative splicing, as these mutations are in the second of two coding exons and also, when we sequenced cDNA made from homozygous *hmx3a^SU42^* mutants we obtained the sequence that we expected (see materials and methods). It is still theoretically possible that there is translational read-through in these two alleles and not in the other *hmx3a* mutant alleles that have obvious abnormal phenotypes. While this seems unlikely given how similar these different alleles are, we cannot rule out this possibility as we have not been able to identify an antibody that is specific to Hmx3a and we could not detect any Hmx3a peptides in SWATH analysis (see discussion below). However, the most parsimonious explanation of our data so far is that Hmx3a does not need its homeodomain for its functions in viability and otolith, lateral line and spinal cord interneuron development. In this case, while Hmx3a may still bind to other DNA-binding proteins and function in transcriptional complexes, unless the N-terminal of Hmx3a contains a novel DNA-binding domain, Hmx3a is not acting as a classic transcription factor (defined in the strict sense as a protein that binds DNA and regulates transcription) during these developmental processes. Nevertheless, as the homeodomain is highly conserved, suggesting that it is still under evolutionary pressure to be maintained, it is possible that Hmx3a has additional functions that do require this domain, either in adult fish or in aspects of development that we did not assay. However, if this is the case, it is still striking that these functions are not required for such fundamental processes as embryonic development and adult viability.

There are a few other examples of homeodomain proteins that can function in some contexts without their homeodomain. For example, protein interaction and over-expression experiments suggest that Lbx2 does not require its homeodomain to enhance Wnt signaling during gastrulation in zebrafish embryos (Lu *et al*. 2014). Instead it sequesters TLE/Groucho, preventing this protein from binding to TCF7L1 and reducing TLE/TCF co-repressor activity. In addition, *homothorax* (*hth*) does not require its homeobox for its functions during Drosophila head development and proximo-distal patterning of the appendages, although the homeodomain is required for antennal development (Noro *et al*. 2006). *hth* has 16 exons and three alternative splice forms. Only one of these isoforms contains the homeobox, but all three contain a protein interaction domain, called the HM domain, that binds to, and can induce the nuclear localization of, Extradenticle (Noro *et al*. 2006). However, in contrast to *hth*, zebrafish *hmx3a* has only two exons and one splice form. While relatively rare, there also examples of transcription factors from other families that only need to bind DNA for some of their functions. For example, Scl/Tal1 has both DNA-binding dependent and DNA-binding independent functions in hematopoietic and vascular development (Porcher *et al*. 1999; Ravet *et al*. 2004).

### Zebrafish *hmx2* may not, by itself, be required for viability or correct development of otoliths, lateral line or spinal cord neurotransmitter phenotypes

The experiments described in this paper also show that *hmx2* single mutants, with progressively larger deletions of the *hmx2* coding sequence from *hmx2^SU37^* mutants (with a 52 bp deletion), through *hmx2^SU38^* mutants (with a 427 bp deletion) to *hmx2^SU39^* mutants (that lack almost all *hmx2* coding sequence), do not exhibit NMD (which can trigger genetic compensation in some circumstances, El-Brolosy *et al*. 2019, Fig. 8L-N) are viable and have no obvious otolith, lateral line or spinal cord interneuron neurotransmitter mutant phenotypes. In addition, *hmx2;hmx3a^SU44^* and *hmx2;hmx3a^SU45^* deletion mutants do not have more severe phenotypes than *hmx3a^SU3^* or *hmx3a^SU43^* single mutants. These results are surprising because zebrafish *hmx2* and *hmx3a* have very similar expression domains during embryonic development (although the spinal cord expression of *hmx3a* does briefly precede that of *hmx2*, Fig. 1B & C), these overlapping expression domains are highly conserved in different vertebrates and studies in other animals suggest that *hmx2* and *hmx3* often act redundantly during development (Wang *et al*. 2004; Wang and Lufkin 2005; Wotton *et al*. 2010). Most notably, previous analyses demonstrated that mouse *Hmx2* mutants had defects in ear development, although interestingly these were more severe in about 70% of homozygous mutants than in the other 30%, showing that there was some variability in the requirement for Hmx2 (Wang *et al*. 2001). In addition, mouse *Hmx2;Hmx3* double mutants had more severe ear phenotypes than either single mutant, as well as defects in hypothalamus and pituitary development that were not found in either single mutant and most of the double mutants died around the 5^th^ day after birth, whereas the single mutants were viable (Wang *et al*. 2004). When considered in combination, these data suggest that mouse *Hmx2* has important functions in ear and brain development, although some of these are redundant with *Hmx3*. In contrast, the only mutational analysis where we detected any function for zebrafish *hmx2*, was when we introduced *hmx2* mutations into hypomorphic *hmx3a^SU42^* mutants. Taken together, our data suggest that while zebrafish Hmx2 protein can function in otolith and lateral line development, it may only impact the development of these structures in embryos with significantly reduced Hmx3a function (less activity than that provided by one functional *hmx3a* allele, as embryos trans-heterozygous for *hmx2;hmx3a^SU44^* and *hmx2^SU39^* develop normally) but more Hmx3a activity than in *hmx3a^SU3^* or *hmx3a^SU43^* mutants.

Our experiments also suggest that the lack of any obvious abnormal phenotypes in *hmx2^SU39^* single mutants is not due to genetic compensation or maternal expression of *hmx* genes. One possible explanation for why zebrafish Hmx2 might have a diminished role in development compared to mouse Hmx2 or zebrafish Hmx3a could be if zebrafish Hmx2 has evolved to be less conserved with mouse Hmx2 and Hmx3 than zebrafish Hmx3a. However, a comparison of all four proteins only reveals six residues that are shared between mouse Hmx2 and Hmx3 and zebrafish Hmx3a but not zebrafish Hmx2, and four of these residues are upstream of the *hmx3a^SU3^* mutation (Fig. S2). Our prior research identified Hmx2 and Hmx3 in all of the different vertebrates that we analyzed, including five different teleost species, and our phylogenetic analyses of these proteins did not suggest that Hmx2 has evolved any faster in zebrafish than in other species, or that Hmx2 has evolved faster than Hmx3 (Wotton *et al*. 2010). Taken together, these observations suggest that there is still considerable evolutionary pressure to maintain zebrafish Hmx2, which in turn suggests that it should have an important role(s) in zebrafish survival and/or reproduction. Therefore, it is surprising that we did not detect more severe consequences from loss of Hmx2. It is possible that Hmx2 has important functions later in development and/or in aspects of development that we did not assay. However, if this is the case, these functions are not required for viability or reproduction as even *hmx2^SU39^* homozygous mutants survive to adulthood and produce viable progeny. It is also possible that *hmx2* has important function(s) in adult fish, as our assays would not have detected this. It would be interesting to investigate these possibilities in future studies.

### Very similar *hmx3a* mutant alleles have different homozygous mutant phenotypes

It is currently unclear why *hmx3a^SU42^* retains more WT function than *hmx3a^SU43^* when both should encode proteins with only 107 WT amino acids. As discussed above, we are confident that this is not due to alternative splicing or exon skipping. We also do not observe any NMD of *hmx3a* mRNA for any of our *hmx3a* single mutant alleles (Fig. 8O-R; the double deletion mutants lack all *hmx3a* coding sequence, so there is no mRNA to assess, Fig. 8I), so the difference in allelic strength is not due to some of the mutant mRNAs being degraded. However, it is possible that different mutant alleles result in different amounts of truncated protein due to differences in translation efficiency or protein stability. Unfortunately, we were not able to test this as there are currently no antibodies that uniquely recognize Hmx3a and we would require an antibody that recognizes the N-terminal region of Hmx3a that should be conserved in our single mutant Hmx3a proteins. In addition, we were unable to detect any Hmx3a peptides in a SWATH analysis (data not shown; Hmx3a has also not been detected in other SWATH analyses Blattmann *et al*. 2019; Lin *et al*. 2019), presumably because, like many transcription factors, it is expressed in either two few cells and/or at too low a level.

It is also possible that the overall length of the mutant protein is important for retaining function and that the additional abnormal amino acids after the frameshift but before the premature stop codon in *hmx3a^SU42^* help this allele retain more WT function. A longer protein sequence might facilitate a required protein conformation and/or binding with other proteins or molecules. If this is the case, then it could also explain why Hmx3a^SU42^ protein (which is predicted to contain 107 WT + 42 abnormal amino acids, Fig.4) appears to retain more WT function than Hmx3a^sa23054^ protein (which should contain only 118 WT amino acids; Fig.4). Currently, there are no known binding partners of Hmx3a. However, if future analyses identify any it would be interesting to test if they can bind to Hmx3a^SU42^ and Hmx3a^SU3^.

Another related possibility is that the different stretches of abnormal amino acids after the frameshift in *hmx3a^SU3^*, *hmx3a^SU42^* and *hmx3a^SU43^* might introduce sequences that influence protein stability, degradation or function. Using a variety of online analysis tools, we did not detect any sumoylation, or PEST sequences in any of the predicted mutant protein sequences and the only ubiquitination motifs that we identified are located in the WT sequence present in all four mutant proteins (Rice *et al*. 2000; Sarachu and Colet 2005; Brameier *et al*. 2007; Radivojac *et al*. 2010; Zhao *et al*. 2014). However, the eukaryotic linear motif prediction tool identified a monopartite variant of a classic, basically-charged nuclear localization signal in Hmx3a^SU4*2*^ protein that is not present in any of the other predicted mutant protein sequences (Via *et al*. 2009; Gould *et al*. 2010; Kumar *et al*. 2020), although this domain was not detected using default parameters with cNLS Mapper or NucPred (Brameier *et al*. 2007; Kosugi *et al*. 2009). This is potentially very interesting as WT Hmx3a has a nuclear localization signal located between amino acids 167-177, overlapping the start of the homeodomain at amino acid 171, which is downstream of the mutations in all of these alleles.

In addition, online tools that identify disordered versus ordered protein structure suggest that both the WT amino acids in Hmx3a^sa23054^ that are not found in the other predicted Hmx3a mutant proteins and the non-WT amino acids in the predicted protein product of *hmx3a^SU42^* may provide longer stretches of disordered sequence than are present at the end of the predicted protein products of *hmx3a^SU3^* or *hmx3a^SU43^* (Linding *et al*. 2003; Dosztanyi *et al*. 2005; Ishida and Kinoshita 2007; Dosztanyi 2018; Meszaros *et al*. 2018; Erdos and Dosztanyi 2020). These findings raise the intriguing possibility that *hmx3a^SU42^* might retain Hmx3a function because it can still localize to the nucleus and/or that *hmx3a^sa23054^* and *hmx3a^SU42^* might retain some WT activity because of the disordered sequences at the end of their predicted proteins. As disordered protein regions can switch between disordered and ordered states in the presence of a binding partner, it is tempting to speculate that these disordered stretches at the C-termini of Hmx3a^SU42^ and Hmx3a^sa23054^ proteins may still be able to bind proteins essential for Hmx3a function that the other alleles cannot (Meszaros *et al*. 2018). Investigation of these possibilities is outside the scope of the current study but would be interesting to address in future work.

### *hmx2* and *hmx3a* morpholino injections produce more severe spinal interneuron phenotypes than *hmx2* and *hmx3a* mutants

Our original morpholino data suggested that all dI2 interneurons might be switching their neurotransmitter phenotypes as the increase in the number of spinal inhibitory cells and the reduction in the number of excitatory spinal cells in DKD embryos were both roughly equal to the number of dI2 interneurons (glutamatergic *hmx3a*-expressing cells). However, even in our *hmx2;hmx3a* deletion mutants, the number of cells changing their neurotransmitter phenotypes is lower than this. The differences between *hmx3a* morpholino-injected embryos and mutant embryos can be seen clearly in our experiment to test whether genetic compensation occurs in *hmx3a^SU3^* homozygous mutants. These data directly compare uninjected mutants with morpholino-injected mutants and WT siblings from the same experiment. While there were a range of morpholino-injected phenotypes, overall the *hmx3a* morpholino-injected WT and *hmx3a^SU3^* mutant embryos had a more severe reduction of glutamatergic cells than uninjected mutants (Table 7). While it is possible that *hmx3a^SU3^* mutants may be slightly hypomorphic, their spinal cord phenotype is the same as *hmx2;hmx3a^SU44^* mutants, in which the *hmx3a* coding sequence is completely deleted. Therefore, the more severe phenotypes in some of the *hmx3a^SU3^* mutants injected with *hmx3a* morpholino cannot be explained by the morpholino removing any residual Hmx3a function. This experiment also suggests that the less severe phenotype in uninjected *hmx3a^SU3^* mutants is not caused by genetic compensation. Consistent with this, we have also shown that this less severe mutant phenotype is not due to other *hmx* genes being upregulated in these mutants or maternal expression of *hmx3a* or any other *hmx* genes. These results are very puzzling. There are several reasons to suggest that the more severe spinal cord phenotype in DKD embryos is not due to non-specific effects of either the *hmx3a* or *hmx2* morpholino. First, we were able to rescue more glutamatergic spinal neurons in our DKD embryos, with co-injection of either *hmx2* or *hmx3a* morpholino-resistant mRNA, than are lost in any of our mutants (*hmx2;hmx3a* deletion mutants have a reduction of ∼ 14 glutamatergic neurons in the region of the spinal cord that we assayed, whereas both mRNA and morpholino co-injection experiments “rescued” about 20 glutamatergic neurons in DKD embryos (Tables 1A & B)). Second, we were able to fully rescue the *hmx3a* SKD phenotype by co-injecting a morpholino-resistant *hmx3a* mRNA, and the *hmx2* SKD phenotype by co-injecting either a morpholino-resistant *hmx2* mRNA or a morpholino-resistant *hmx3a* mRNA. Third, it is unlikely that the more severe phenotype is due to cell death or a delay in embryo development (which are common non-specific side effects of morpholino injections), as there was no change in the number of *hmx3a*- or *en1b*-expressing spinal cord cells in DKD embryos and there was an increase in the number of inhibitory spinal cord interneurons equivalent to the reduction in glutamatergic neurons. Finally, it is also unclear, why a non-specific effect of a morpholino would exacerbate the real loss-of-function phenotype, causing additional spinal cord interneurons to lose their glutamatergic phenotypes and instead become inhibitory. This suggests that if the more severe morpholino-injection phenotypes are due to non-specific effects of the morpholinos, these non-specific effects produce an identical phenotype to the specific knock-down effect, namely a switch in neurotransmitter phenotype.

It is also puzzling why *hmx2* morpholino-injected SKD embryos have reduced numbers of spinal cord glutamatergic cells and an increase in the number of inhibitory spinal interneurons, while *hmx2^SU39^* mutants do not, and our experiments suggest that this is also not due to genetic compensation. In addition, co-injection of a morpholino-resistant *hmx2* mRNA rescues DKD embryos as well as *hmx3a* co-injection, even though WT *hmx2* is not sufficient for normal development in *hmx3a^SU3^* or *hmx3a^SU4^*^3^ mutants. The latter result could be explained if the RNA injection provides higher levels of Hmx2 function than is normally found endogenously. However, the first result is harder to explain. As discussed above, our data suggest that it is unlikely that the *hmx2* morpholino has non-specific effects on neurotransmitter phenotypes in the spinal cord. These results are also not due to cross-reactivity of the *hmx2* translation-blocking morpholino with *hmx3a*. The *hmx3a* and *hmx2* translation-blocking morpholino sequences are completely different from each other and there is no homology between the *hmx2* translation-blocking morpholino and *hmx3a* upstream or coding sequence. When we BLAST the *hmx2* translation-blocking morpholino sequence against the zebrafish genome the only homology is with *hmx2* (25/25 residues) and with intronic sequence for a gene *si:dkey-73p2.3* on chromosome 3 (18/25 residues), which is predicted to encode a protein with GTP-binding activity. Similarly, when we blast the *hmx3a* translation-blocking morpholino sequence against the zebrafish genome the only homology is with *hmx3a* (25/25 residues).

One intriguing alternative possibility that could explain the apparent specificity of the additional phenotype in the morpholino-injected embryos could be that Hmx3a and Hmx2 are acting as fate guarantors, to make the normal neurotransmitter phenotype more robust, as has previously been described for some transcription factors (Topalidou *et al*. 2011; Zheng *et al*. 2015; Zheng and Chalfie 2016). In these cases, mutating the gene usually caused a partially penetrant phenotype in ideal conditions but a more severe phenotype in stressed conditions. In this case, the injection of morpholinos could be such a stressed condition, and this could account for the more severe phenotypes that we see in morpholino knock-down experiments compared to mutational analyses. Future analyses could investigate this possibility by testing if other stressed conditions increase the severity of *hmx3a^SU3^* or *hmx2^SU39^* single mutant or *hmx2;hmx3a* deletion mutant phenotypes. Even if this is not the case, our results suggest that something other than just non-specific effects from the morpholinos may be occurring. Therefore, we felt that it was crucial to report this morpholino injection data, as an intriguing and hopefully thought-provoking contribution to the continuing discussion about the pros and cons of using morpholinos to investigate gene function.

### *hmx3b* has very limited expression during early zebrafish embryogenesis

The data in this paper also provide the first characterization of zebrafish *hmx3b* expression. Surprisingly, given the expression of all of the other four *hmx* genes during zebrafish embryonic development, the only expression of *hmx3b* that we have been able to detect, is very weak expression in the hindbrain from 36-48 h (Fig. 1 S’, X’ & AC’). When we performed our earlier analyses of vertebrate NK genes, we did not find *hmx3b* in either the Zv7 or Zv8 versions of the zebrafish genome (Wotton *et al*. 2010). It did not appear until Zv9. Interestingly, despite *Hmx2* and *Hmx3* being closely linked in all vertebrates examined so far, only one *hmx2* gene has been found in the zebrafish genome and zebrafish *hmx3b* is not located within any of the previously described duplicated NK homeobox clusters, including the teleost duplications: it is located on chromosome 12, separate from any other *nk* genes (Wotton *et al*. 2010). However, *hmx3b* is flanked by two genes, *bub3* and *acadsb*, that flank *Hmx2* and *Hmx3* in the human, mouse, chicken and xenopus genomes, suggesting that part of the NK cluster previously identified on chromosome 1 may have translocated to chromosome 12. This is maybe not surprising as zebrafish *hmx2* and *hmx3a* have also translocated from the rest of their NK cluster on chromosome 13 to chromosome 17 and zebrafish *hmx1* and *hmx4* have translocated from the rest of their cluster on chromosome 14 to chromosome 1 (Wotton *et al*. 2010). It is possible that this translocation and the loss of surrounding sequences may explain the different expression pattern of *hmx3b* compared to *hmx3a*. For example, in previous analyses we identified three highly conserved non-coding regions in the vicinity of *hmx2* and *hmx3a* (two upstream of *hmx3a* and one in between *hmx3a* and *hmx2*) that are conserved in mammals, frog and teleosts, (Wotton *et al*. 2010) but none of these regions are present near *hmx3b* (data not shown).

In conclusion, in this paper we provide the first description of zebrafish *hmx3b* expression. Our results also identify the spinal cord cells that express *hmx2* and *hmx3a* (dI2 and V1 interneurons) and uncover novel functions for *hmx3a* in correct specification of a subset of spinal cord neurotransmitter phenotypes and in lateral line progression. Our data suggest that while *hmx3a* is required for viability, correct otolith development, lateral line progression and specification of a subset of spinal neurotransmitter phenotypes, *hmx2* is not, by itself, required for any of these developmental processes, although it can act partially redundantly with *hmx3a* in situations where *hmx3a* function is significantly reduced, but not completely eliminated. Finally, our results also suggest that Hmx3a may not require its homeodomain for its roles in viability or embryonic development. Taken together, these findings significantly enhance our understanding of spinal cord, ear and lateral line development and suggest that, intriguingly, more homeodomain proteins may not require their homeodomain for many of their essential functions.

## Acknowledgments

We thank ZFIN for providing information on nomenclature and other essential zebrafish resources and Henry Putz, Jessica Bouchard, Paul Campbell, Annika Swanson and several SU undergraduate fish husbandry workers for help with maintaining zebrafish lines. We also thank Professor Jason Fridley and Associate Professor Qiu Wang for help with statistical analyses and William Haws and Arshi Mustafa for comments on previous drafts of the manuscript. This research was primarily funded by NINDS R01:NS077947 with some support from New York State Spinal Cord Injury Fund Contract # C32253GG and NSF IOS 1755354.

## Author Contributions

SJE created all the *hmx* mutants described in this study except *hmx3a^sa23054^*, performed most of the morpholino and mutant experiments, all of the statistical analyses and prepared the figures and some of the tables; GAC performed most of the double-labeling experiments and some of the morpholino analyses; AK, TS and GG performed some of the genotyping and *in situ* hybridization experiments and AK also performed some of the double-labeling experiments; KEL conceptualized and directed the study, acquired the financial support for the project, contributed to data analysis, made some of the tables and wrote the paper. All authors read and commented on drafts of the paper and approved the final version.

---

**Figure S1.**
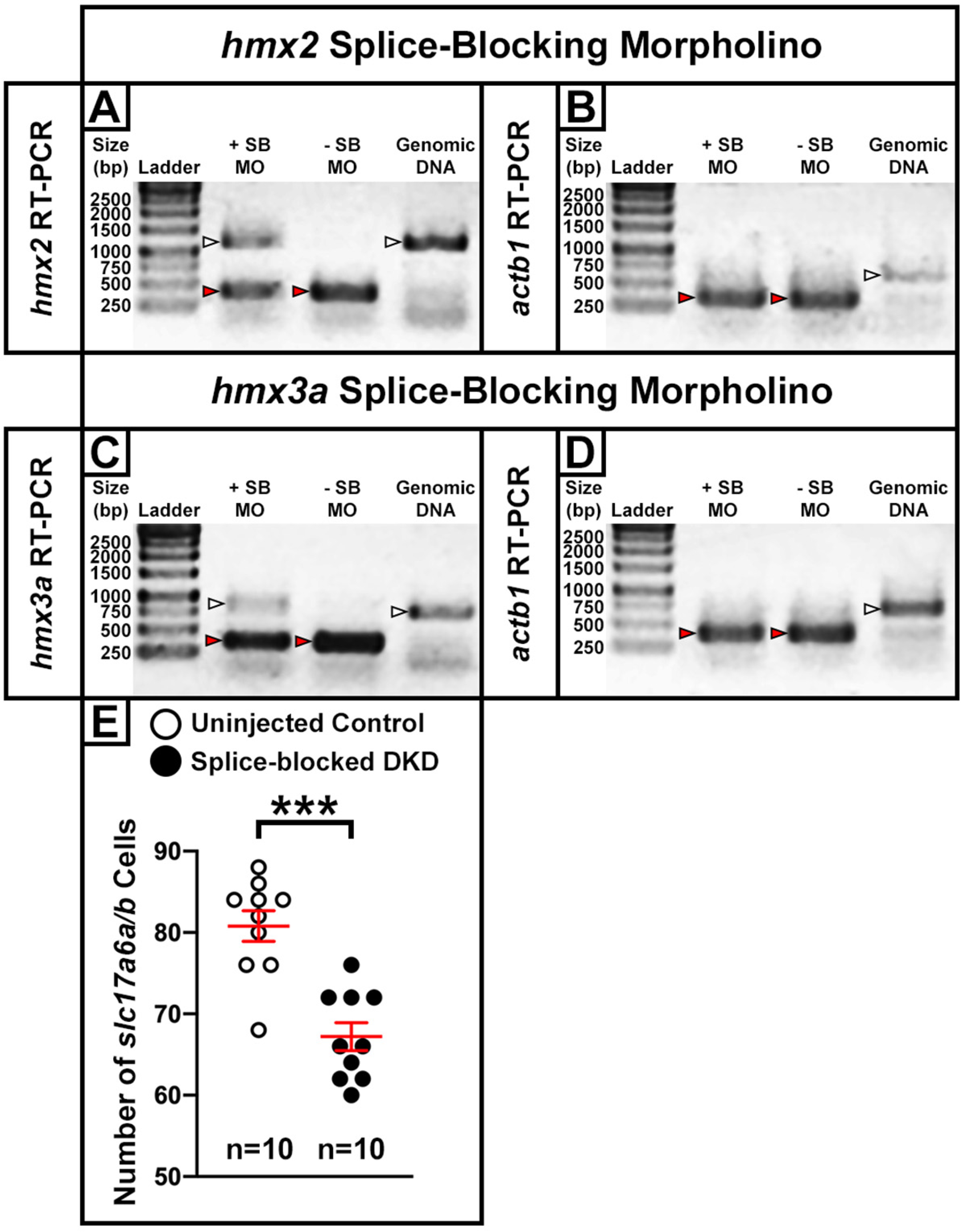
***hmx2;hmx3a* double knock-down with splice-blocking morpholinos. (A-D)** RT-PCR analysis of embryos injected at the one cell stage with 20 ng of *hmx2* **(A-B)** or *hmx3a* **(C-D)** splice-blocking morpholino, plus 35 ng of the zebrafish control *p53* morpholino (+ SB). RT-PCR analysis was also performed on uninjected WT control embryos (- SB). Controls without reverse transcriptase, using non-DNase-treated templates (genomic DNA) are included to help identify the unspliced amplicon (see materials and methods for further information). Molecular weight marker (ladder) fragment sizes are shown in base pairs on the LHS. **(A, C)** Even at the maximum dose tolerated by the embryos, neither injection of *hmx2* **(A)** or *hmx3a* **(C)** splice-blocking morpholino is sufficient to block splicing of all *hmx2* or *hmx3a* transcripts. White arrowheads: unspliced product *(hmx2* = 1204 bp, *hmx3a* = 779bp); red arrowheads: spliced product *(hmx2* = 426 bp, *hmx3a* = 393 bp). **(B, D)** Neither *hmx* splice-blocking morpholino exerts non-specific effects on the splicing of transcripts of the housekeeping gene *actb1* (white arrowheads: unspliced product= 697 bp, red arrowheads: spliced product= 387 bp). **(E)** Number of cells expressing *s/c17a6a/b* in a precisely-defined spinal cord region on one side of the body adjacent to somites 6-10 at 27 h. For ease of comparison with data throughout the rest of the paper, the cell counts shown here have been doubled to represent both sides of the spinal cord. Data is depicted as an individual value plot and the n-values for each genotype are also shown. The wider red horizontal bar depicts the mean number of cells (uninjected control: 80.8, *hmx2;hmx3a* DKD: 67.2) and the red vertical bars depict the standard error of the mean (standard error of the mean (S.E.M.), uninjected control: ± 1.9, *hmx2;hmx3a* DKD: ± 1.7). All counts are an average of 10 embryos. The statistically significant (p < 0.001) comparison is indicated with brackets and three asterisks. All data were first analyzed for normality using the Shapiro-Wilk test. The data are normally distributed and so the F test for equal variances was performed. Since the variances were equal, a type 2 student’s t-test was performed (p = <0.0001). These data show that following co-injection of *hmx2* and *hmx3a* splice-blocking morpholinos, there is a statistically significant reduction in the number of excitatory (s/c17a6a/b-expressing) cells in *hmx2;hmx3a* DKD embryos compared to uninjected control embryos (14 fewer cells).

**Figure S2.**
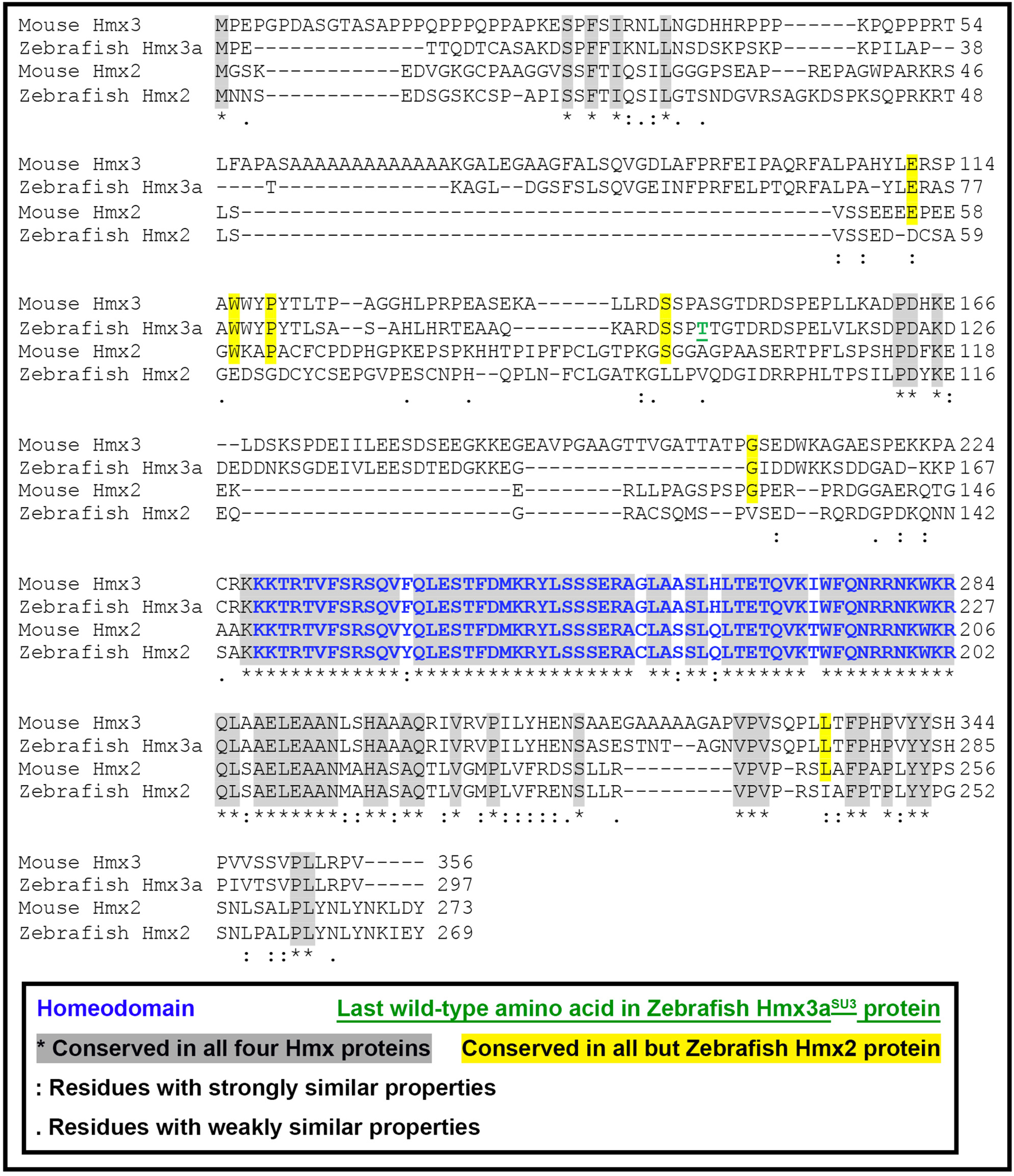
**Alignment of mouse and zebrafish Hmx2 and Hmx3(a) protein sequences.** Alignment of full-length mouse and zebrafish Hmx2 and Hmx3(a) protein sequences. Alignment performed using Clustal Omega version 1.2.4 and default parameters (Madeira et al. 2019). Blue text = homeobox domain. Green underlined text = last WT amino acid of Hmx3asu ^3^protein. Gray shaded residues are conserved between all four Hmx proteins. Yellow shaded residues are conserved between both mouse Hmx proteins and zebrafish Hmx3a protein, but not zebrafish Hmx2 protein. : = amino acids with highly similar properties; . = amino acids with weakly similar properties. Other than the start methionine, upstream of the homeobox domain only 8 other amino acids are conserved between all four Hmx proteins. In contrast, 26 amino acids are conserved between all four Hmx proteins downstream of the homeobox domain. As expected, the homeobox domain is highly conserved (91%, 52/57 amino acids) across all four Hmx proteins.

**Table S1.**
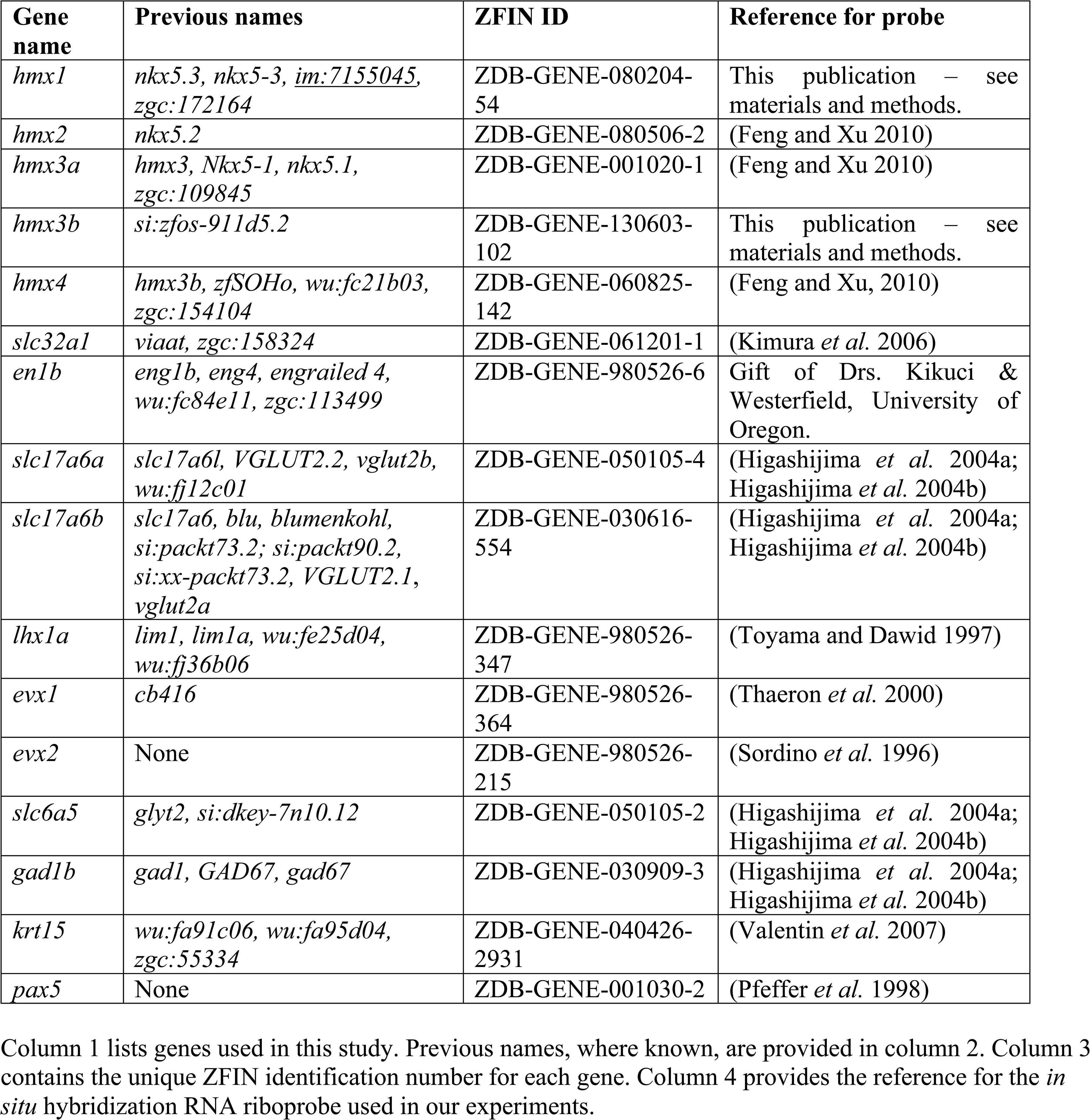
Gene names, previous names, ZFIN identifiers and references for *in situ* hybridization probes.

**Table S2.**
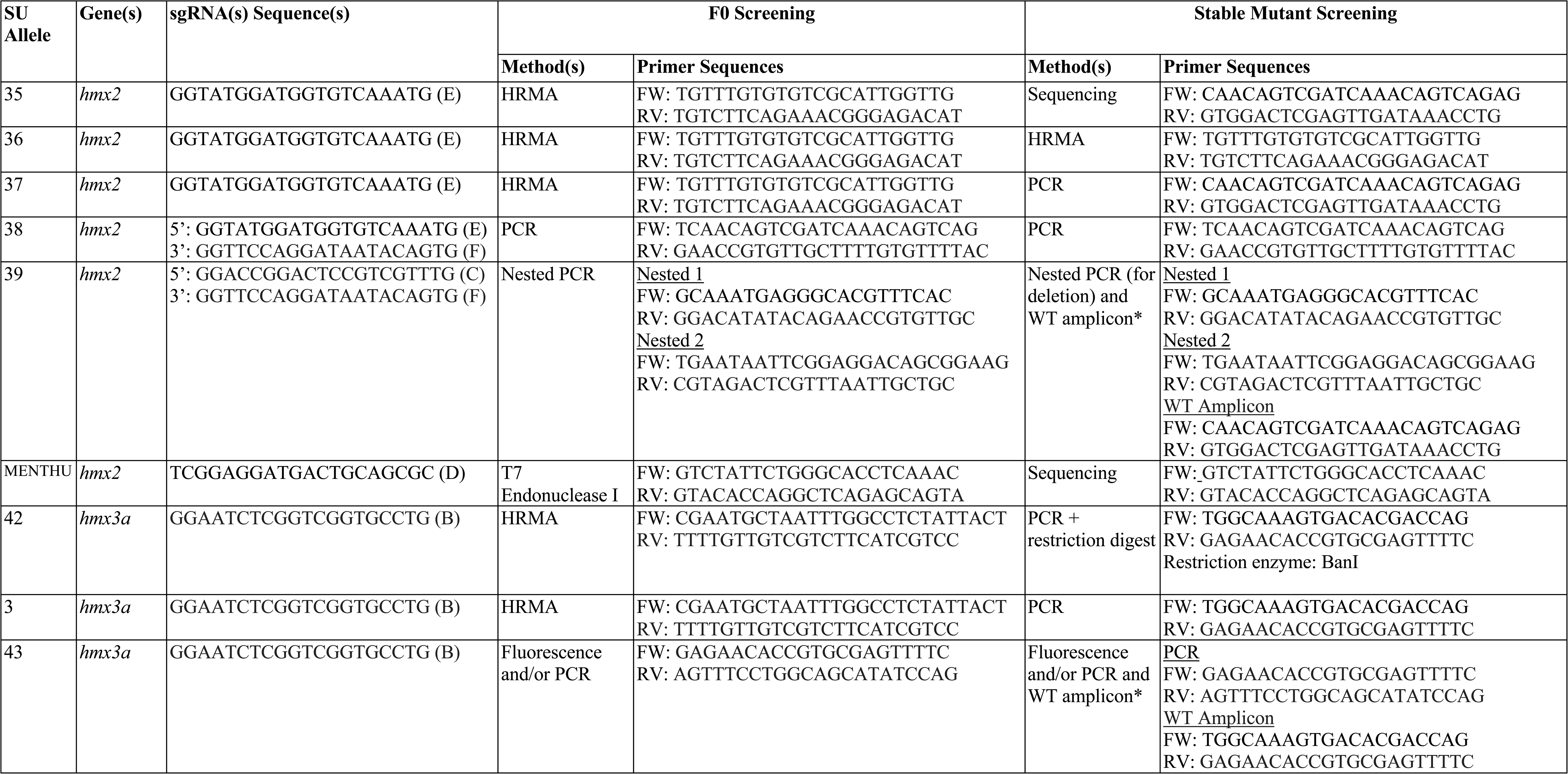
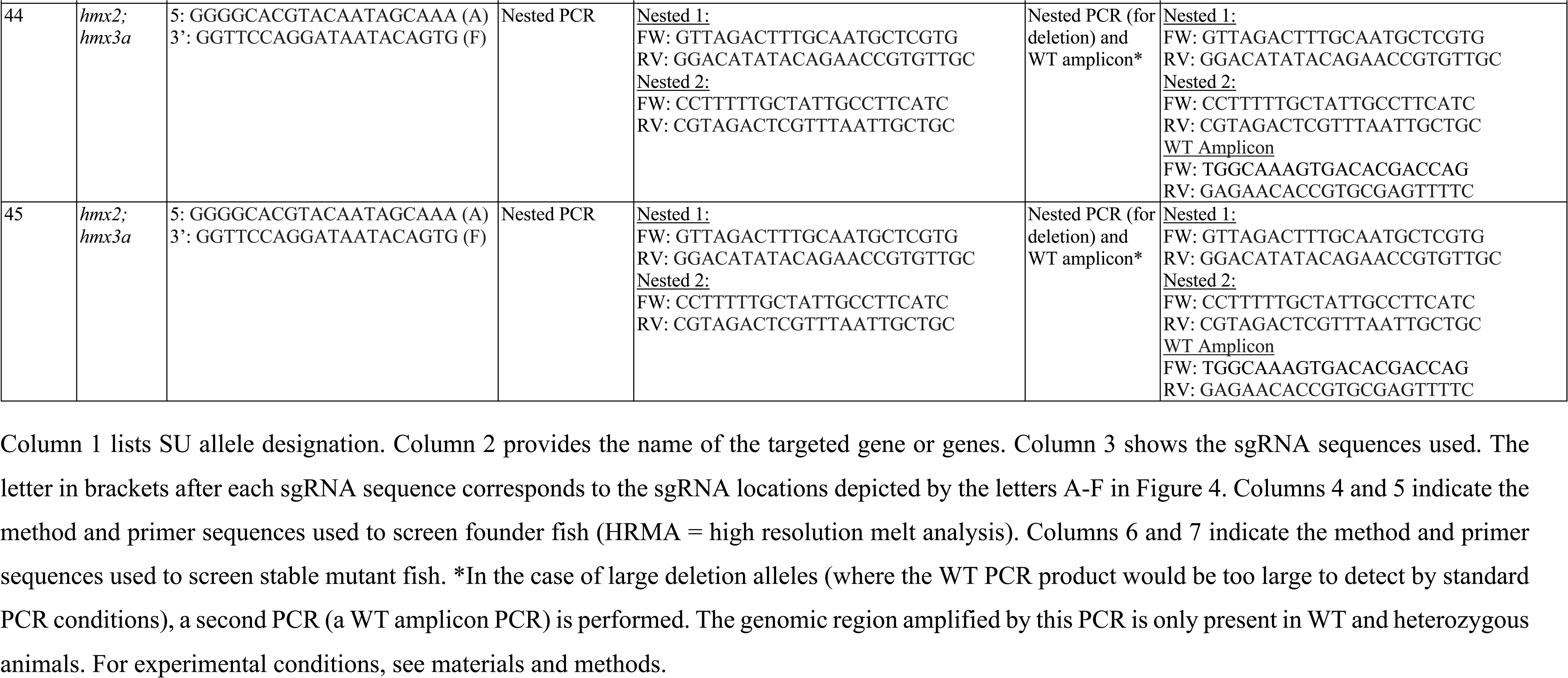
sgRNA and primer sequences used for *hmx*2, *hmx3a* and *hmx2;hmx3a* CRISPR mutagenesis.

## Notes

### Competing Interest Statement

The authors have declared no competing interest.

### Summary of Updates

This version of the manuscript has been revised to make the figures next to figure legends and correct some typos.

